# Endogenous opioid dynamics in the dorsal striatum sculpt neural activity to control goal-directed action

**DOI:** 10.1101/2024.05.20.595035

**Authors:** Raajaram Gowrishankar, Madelyn Hjort, Abigail J Elerding, Sofia E Shirley, Josie Van Tilburg, David J Marcus, Pranav Senthilkumar, Khalid A Abrera, Dustin Sumarli, Kat Motovilov, Valerie Lau, Adam A Gordon-Fennell, Zhe C Zhou, Chunyang Dong, Lin Tian, Garret D Stuber, Michael R. Bruchas

## Abstract

Endogenous neuropeptides are uniquely poised to regulate neuronal activity and behavior across multiple timescales. Traditional studies ascribing neuropeptide contributions to behavior lack spatiotemporal precision. The endogenous opioid dynorphin is highly enriched in the dorsal striatum, known to be critical for regulating goal-directed behavior. However, the locus, the precise timescale, or functional role of endogenous dyn-KOR signaling on goal-directed behavior is unknown. Here, we report that local, time-locked dynorphin release from the dorsomedial striatum is necessary and sufficient for goal-directed behavior using a suite of high resolution modern approaches including *in vivo* two-photon imaging, neuropeptide biosensor detection, conditional deletions and time-locked optogenetic manipulations. We discovered that glutamatergic axon terminals from the basolateral amygdala evoke striatal dynorphin release, resulting in retrograde presynaptic GPCR inhibition during behavior. Collectively, our findings isolate a causal role for endogenous neuropeptide release at rapid timescales, and subsequent GPCR activity for tuning and promoting fundamental goal-directed behaviors.

## 1 INTRODUCTION

Most of the behaviors that animals perform are defined as “goal-directed”. At its simplest, goal-directed behavior can be summarized as an animal’s ability to associate, sustain and flexibly update the predictive relationship between an action and a subsequent outcome. This behavior is fundamental to survival and relies on stable learning across multiple timescales. This includes – a timescale of seconds in which animals must prospectively look for causal relationships between actions and outcomes, and days where animals learn the associated probabilities between these causal actions and outcomes. Dysfunctions in goal-directed behavior have been implicated in numerous neuropsychiatric disorders including obsessive compulsive disorder (Balleine et al. 2007), depression (Ironside et al. 2020), chronic stress (Yoshida et al. 2021) and substance use disorders (SUDs) (Hogarth 2020). The dorsal striatum plays an integral role in enabling goal-directed behavior in rodents (Balleine et al. 2007), non-human primates (Hikosaka et al. 1989); and humans (O’Doherty et al. 2002) with the medial portion of dorsomedial striatum (DMS) implicated most specifically (Yin et al. 2005, Corbit and Janak 2010).

Canonically, functional roles for goal-directed behavior have been ascribed to largely two cell-types – the direct pathway medium spiny projection neurons (MSNs) expressing the dopamine D1 receptor/dynorphin (D1 SPNs) and the indirect pathway spiny projection neurons expressing the dopamine D2 receptor/Enkephalin (D2 SPNs) (Gerfen and Surmeier 2011, Shan et al. 2014). Prior studies have implicated both MSN populations in the acquisition and maintenance of goal-directed action-outcome behaviors (Gremel and Costa 2013, Bloem et al. 2017, Peak et al. 2020). However, either the timescale of manipulation studies do not match the real-time timescale of the behavior, or lack the capabilities to determine activity across large populations of DMS neurons (thousands), and across multiple timescales. Performing second-by-second *in vivo* monitoring of these cell-types longitudinally across behavior facilitates an understanding of how goals, actions, and outcomes are encoded within the DMS neurocircuitry.

Neuropeptides are expressed widely in the brain and can be released in a diffuse manner, thereby exerting their effects at both fast (seconds) and slow (minutes, hours and days) timescales (**?**). Signaling via their cognate G protein-coupled receptors (GPCRs), neuropeptides can control neural activity by engaging a variety of downstream effectors, notably ion channels (fast), and cAMP production to impact neuronal plasticity and gene transcription (slow) (van den Pol 2012). The neuropeptide dynorphin is expressed exclusively in D1 SPNs in the striatum (Reiner and Anderson 1990) and has been implicated in stress regulation, dysphoria, anxiety-like behavior and drug-seeking behavior (Shippenberg et al. 2007, Bruchas et al. 2010, Tejeda and Bonci 2019, Limoges et al. 2022). Dyn causes long-lasting changes in brain activity via its cognate inhibitory Gai-coupled G protein-coupled receptor, the kappa opioid receptor (KOR) (Chavkin et al. 1982) and dampens neuronal activity over slower timescales (minutes to hours) (Bruchas et al. 2010). However, Gai-coupled G protein-coupled receptor signaling, (for eg. dyn-KOR signaling) can also work at faster timescales due to their coupling to calcium channels and potassium channels (Al-Hasani and Bruchas 2011, Corder et al. 2018). This inhibitory function, via neuropeptide-GPCR signaling, could be highly relevant for creating, shaping and stabilizing animal behavioral sequences, as well as establishing learning through retrograde feedback (Marder 2012, Nair et al. 2023).

Studies have shown that Gi-coupled GPCRs are expressed widely across different neuronal compartments and can influence neuronal activity via both pre and post-synaptic mechanisms (Atwood et al. 2014). KOR can be expressed on dendrites, cell bodies, and presynaptic axon terminals, suggesting dyn-KOR signaling could modulate post-synaptic neuron activity and/or presynaptic neurotransmitter release (Drake et al. 1996, Svingos et al. 1999). In vitro electrophysiology studies have shown that KOR activation in the striatum inhibits the presynaptic release from a variety of inputs (Hjelmstad and Fields 2003, Mu et al. 2011), and more recently, onto D1 SPNs via dyn-KOR control of the basolateral amygdala (BLA) inputs (Tejeda et al. 2017). KOR mRNA is enriched in >50% of BLA neurons (Nygard et al. 2016). Decades of work has established the BLA as an important hub for action-outcome associations and goal-directed learning (Baxter and Murray 2002, Ostlund and Balleine 2008, Parkes and Balleine 2013, Wassum and Izquierdo 2015, Courtin et al. 2022). Importantly, studies in rodents (Pan et al. 2010, Corbit et al. 2013, Wall et al. 2013) and non-human primates (Cho et al. 2013) have established anatomically the existence of direct, excitatory projection from the BLA to the DMS. More recently, studies have suggested a role for DMS-projecting BLA neurons in goal-directed behavior (Courtin et al. 2022, Giovanniello et al. 2023); yet how BLA-DMS projections influence fundamental goal-directed behaviors has been surprisingly unclear.

Here we sought to decode the potential feedback neuromodulation mechanisms of inhibitory peptidergic control of a novel amygdalar-basal ganglia pathway. We aimed to provide a clearer fundamental understanding of how neuropeptides dynamically control essential inputs to the basal ganglia to shape goal-directed behavior. We achieved recordings of over 14,000 DMS neurons using specialized implantable microprisms in conjunction with *in vivo* 2-photon calcium imaging. We report that DMS pdyn neurons are preferentially engaged during learning behavior, and discovered that they specifically encode goal-directed action or cued anticipation of reward. Next, using a dyn biosensor we recently developed, we found that dyn is released in a rapid dynamic manner specifically during the cued anticipation of reward, following action. Finally, we use a suite of high-resolution approaches including conditional deletion, optogenetics, multiplexed with *in vivo* fiber photometry and 2-photon calcium imaging to isolate a foundational mechanism by which inhibitory neuropeptide-GPCR signaling influences neuronal activity and stabilizes goal-directed behaviors.

## RESULTS

### DMS^pdyn^ neurons encode goal-directed actions and cues

To measure the activity of DMS neurons during goal-directed action-outcome behavior, we imaged the activity of over 14,000 neurons using 2-photon calcium imaging through implanted microprisms (we recently developed and characterized this specialized approach, see (Hjort et al. 2024)) during head-fixed operant behavior. We injected D1R-TdTomato mice with a virus packaging a genetically-encoded calcium sensor, AAVDJ-hsyn-GCaMP6s and implanted a 1.5×1.5×8 mm microprism in the DMS (**Fig. 1A, S1A**). Following lick training for 10% sucrose and Pavlovian conditioning to associate a tone to sucrose delivery, mice were next trained to rotate a wheel in an “active” direction to obtain the cue and the reward, with rotations in the “inactive” direction yielding nothing. We previously showed that mice learn operant responding and the reversal of conditions (Gordon-Fennell et al. 2023a). Mice progressed to making significantly more active rotations across 8 days of learning (**Fig. S1B**), and increase the frequency of their overall operant responses that achieve reward (decreasing the distance between reward receipts) (**Fig. S1C**,**D**). Supporting that these responses are goal-directed, when sucrose delivery upon an active rotation was omitted in an extinction regime, mice rapidly reduce their behavior (**Fig. S1B**). In order to collectively capture all the variables in the progression of action-outcome sequences across individual animal variance in a single metric, we calculated an operant index (See **Methods**). This combinatorial operant index displays the expected characteristics of operant learning and extinction and accounts for action vigor (total actions), action discrimination (difference between active and inactive actions) and outcome consumption (whether the reward was actually consumed). In the case of extinction, we used the number of licks the animals engaged in an empty sipper (**Fig. 1B**).

**FIGURE 1.**
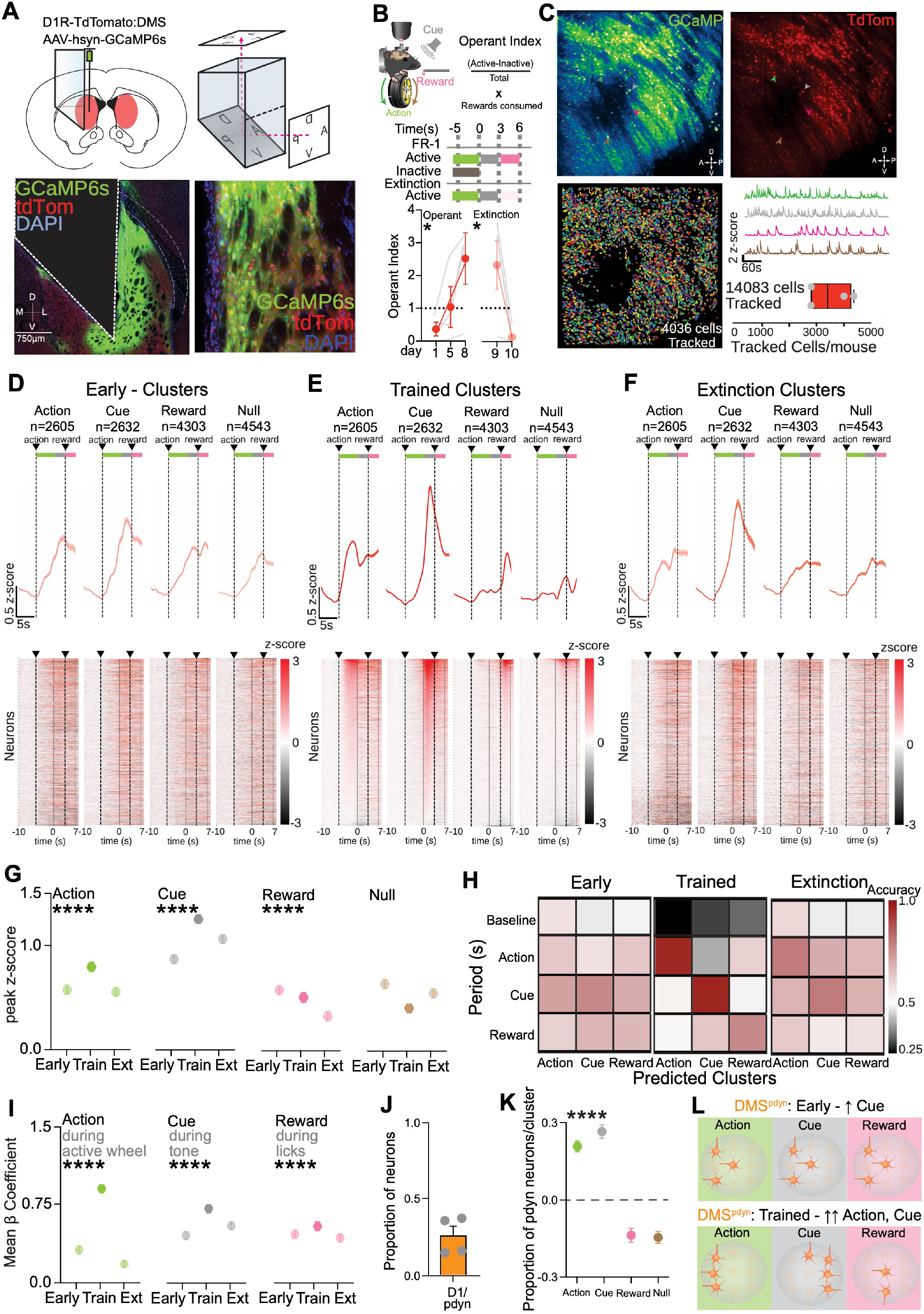
DMS^pdyn^ neurons encode goal-directed actions and cues. (**A**) Top - Schematic of viral injection and prism implantation in the DMS. Bottom – 20X Confocal image of prism implant in the DMS (left) and 40X confocal image of cells stained with DAPI, expressing GCaMP6s and Td-Tomato. (**B**) Top - Schematic of experiment and operant behavior schedule using a head-fixed wheel-based task. Bottom - Operant learning and extinction behavior. Left – Operant Learning (n=4mice; Simple Linear Regression, R^2^=0.4158, *p=0*.*0236\**, F=7.117). Right – Operant Extinction (n=4mice; Simple Linear Regression, R^2^=0.5882, *p=0*.*0264\**, F=8.571). (**C**) Top – Representative standard deviation images from videos of GCaMP6s and Td-tomato fluorescence (from multiple sessions). Bottom - tracked cells from Suite2p and average tracked cells across 4 mice. (**D**) Spectral clustering classification of early operant data across 4 mice showing fluorescence activity traces (top) and heat maps (bottom) of cells from all clusters across time. (**E**) Spectral clustering classification of trained operant data across 4 mice showing fluorescence activity traces (top) and heat maps (bottom) of cells from all clusters across time (n=4mice). (**F**) Spectral clustering classification of operant extinction data across 4 mice showing fluorescence activity traces (top) and heat maps (bottom) of cells from all clusters across time. (**G**) Peak z-score quantification of **Action** clusters (n=4mice; One Way ANOVA, *p<0*.*0001\*\*\*\**, F(2,7812)=32.76. Multiple comparisons – early vs. trained, *p<0*.*0001\*\*\*\**, trained vs. extinction, *p<0*.*0001\*\*\*\**). Peak z-score quantification of **Cue** clusters (n=4mice; One Way ANOVA, *p<0*.*0001\*\*\*\**, F(2,7893)=139.9. Multiple comparisons – early vs. trained, *p<0*.*0001\*\*\*\**, trained vs. extinction, *p<0*.*0001\*\*\*\**, early vs. extinction, *p<0*.*0001\*\*\*\**). Peak z-score quantification of **Reward** clusters (n=4mice; One Way ANOVA, *p<0*.*0001\*\*\*\**, F(2,12906)=30.4. Multiple comparisons – trained vs. extinction, p<0.0001****, early vs. extinction, *p<0*.*0001\*\*\*\**). Peak z-score quantification of **Null** clusters (n=4mice; One Way ANOVA, *p<0*.*0001\*\*\*\**, F(2,13626)=59.21. Multiple comparisons – early vs. trained, *p<0*.*0001\*\*\*\**, trained vs. extinction, *p<0*.*0001\*\*\*\**, early vs. extinction, *p<0*.*0001\*\*\*\**). (**H**) Recurring Neural Network (RNN) modelling of accuracy of decoding to predict cluster classification of early, trained and extinction clusters across their respective time periods (n=4mice). (**I**) Generalized Linear Regression Modeling (GLM) quantification of mean b coefficients of **Action** clusters during **active wheel rotations** (n=4mice; One Way ANOVA, *p<0*.*0001\*\*\*\**, F(2,1653)=417.1. Multiple comparisons – early vs. trained, *p<0*.*0001\*\*\*\**, trained vs. extinction, *p<0*.*0001\*\*\*\**, early vs. extinction, p<0.0001****), **Cue** clusters during **tone** (n=4mice; One Way ANOVA, p<0.0001****, F(2,6552)=31.22. Multiple comparisons – early vs. trained, *p<0*.*0001\*\*\*\**, early vs. extinction, *p<0*.*0001\*\*\*\**), and **Reward** clusters during **licks** (n=4mice; One Way ANOVA, *p<0*.*0001\*\*\*\**, F(2,6552)=31.22. Multiple comparisons – early vs. trained, p<0.0001****, trained vs. extinction, *p<0*.*0001\*\*\*\**, early vs. extinction, *p=0*.*011\**). (**J**) Proportion of td-tomato positive (DMS^pdyn^) cells (n=4mice; paired t test, *p=0*.*0046\*\**, t=7.657, df=3). (**K**) Proportion of DMS^pdyn^ cells enriched in behavioral clusters (n=4mice; One Way ANOVA, *p<0*.*0001\*\*\*\**, F(3,12)=80.96. Multiple comparisons – Cue vs. Reward, *p<0*.*0001\*\*\*\**, Cue vs. Null, *p<0*.*0001\*\*\*\**, Action vs. Reward, *p<0*.*0001\*\*\*\**, Action vs. Null, *p<0*.*0001\*\*\*\**). (**L**) Summary schematic of interpretation of results.

Simultaneous to the behavior described above, we imaged the activity of a total of 14,083 individually-tracked neurons across multiple weeks using 2-photon imaging of GCaMP activity. Furthermore, we identified D1/dyn neurons by obtaining a static image of td-Tomato fluorescence at the end of every session to co-register neurons with their calcium imaging data (**Fig. 1C**). From trial-averaged data when the normalized operant index was greater than 1, we qualitatively observed DMS neurons with activity correlated to three behavioral variables – action, cued anticipation or reward. To quantify this, we used spectral clustering (Hirokawa et al. 2019, Namboodiri et al. 2019) to group individual neurons based on their activity on the trained operant day (day 8) in an unbiased manner. Unbiased spectral clustering revealed distinct clusters of neurons with activity that centered around defined behavioral variables – 18.5% of DMS neurons were active during action, 18.7% of DMS neurons were active during cue, 30.6% of DMS neurons were active during reward consumption, and 32.2% of DMS neurons were relatively inactive during these periods, suggesting a role in other behaviors (**Fig. 1E**). To determine if this grouping was inherent to the neurons, or evolved across goal-directed learning, we applied the same spectral clustering classification to the early (day 1) and extinction (extinction day 1) days. Remarkably, we found that individually-tracked DMS neurons displayed significantly different and/or reduced patterns of activity across the same action-outcome trials early in operant learning compared to when the animal was well trained in acquiring the behavior (**Fig. 1D,G**). This functional heterogeneity further developed on day 5, where animals were familiar with the task, but still hadn’t reached optimal operant responding (**Fig. S1E,F**). Furthermore, this correlated activity to behavior diminished when the animals extinguished their behavior as rewards were omitted (**Fig. 1F,G**). To strengthen the link between activity in clustered DMS populations and task-related behavior, we then trained a recurrent neural network (RNN) model to classify neurons into their respective clusters based on their activity on baseline, action, cue or reward epochs. The model performed poorly on shuffled data (**Fig. S1G**) but accurately decoded cluster classification across training days using only active neurons in the relevant window (**Fig. 1H**), supporting the specialization of DMS neuron activity during operant learning. This result strengthens the evidence for DMS neuron activity specialization across operant learning. To provide insight into whether discrete task windows are predictive of activation in clustered subpopulations of DMS neurons, we used a linear regression/generalized linear model (GLM) fit of the fluorescent activity, trained on active wheel rotations (action), tone delivery (cue) and licks (reward). 32% of action neurons, 83% of cue neurons and 37% of reward neurons showed significant mean b coefficients related to their corresponding behavioral events, indicating that their activity was modulated during each respective behavior (**Fig. S1H**). In the trained operant session, the neurons associated with action, cue, and reward exhibited a significantly higher mean beta coefficient relative to the corresponding behavioral event (**Fig. S1I**). This was notably higher compared to the early and extinction sessions (**Fig. 1I**), indicating that the relationship between DMS neural activity and specific task events strengthens over the course of learning. Taken together, these results suggest that a significant proportion of DMS neurons encode discrete behavioral variables, i.e., action, cue, or reward.

Next, we isolated the specific relative contribution of D1/dyn neurons to these discrete behavioral events during action-outcome behavior. Using a static image of td-Tomato (labelling D1/dyn neurons) at the end of each imaging session, we determined that 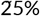 of the individually-tracked neurons were D1/dyn-positive (DMS^pdyn^) (**Fig. 1J**). We then measured the relative classification of these neurons, based on their identities, to each of the established behavioral neuronal clusters. Importantly, we found that DMS^pdyn^ neurons were enriched in action and cue clusters, suggesting that DMS^pdyn^ neurons encode action and cue (**Fig. 1K**).Taken together, these results establish DMS neurons as critical encoders of distinct components of goal-directed behavior across the slower temporal learning domain, while DMS^pdyn^ neurons encode preferentially responses to action and cue with a more dynamic moment-to-moment engagement (**Fig. 1L**).

### DMS dynorphin is released upon cued anticipation of reward

Our results indicated that that DMS^pdyn^ neurons are active during action and cue as animals learn to optimize goal-directed behavior. To determine whether dyn is dynamically released in the DMS during these behavioral epochs, we used a genetically-encoded fluorescent sensor (kLight1.3a) which is a modified KOR that detects ligand binding. We recently characterized this tool in our group to be selective for dyn, and other KOR agonists (Tian et al. 2023). We injected KOR-cre mice with AAV5-CAG-DIO-kLight1.3a and implanted with a 400 mm optic fiber (**Fig. 2A,B, S2A**) into the DMS. Before undergoing the behavioral task, we first characterized klight1.3a sensor performance. When mice received an injection with a KOR agonist U50,488 (i.p., 10 mg/kg) or U50,488 and a short-acting, reversible KOR antagonist aticaprant (Lowe et al. 2014) (i.p.,5 mg/kg), we observed significant, agonist-induced increases in kLight-mediated fluorescence that were blocked by aticaprant (**Fig. S2B-D**). Mice that showed significant elevations to U50 injections (blocked by aticaprant) were food-restricted and underwent Pavlovian conditioning to associate a houselight to sucrose pellet delivery. Mice were then trained in an operant conditioning paradigm to make nosepokes into an “active” port to obtain the cue and the reward, with “inactive” nosepokes yielding nothing. Mice progressed toward making significantly higher active and lower inactive nosepokes on day 5 (trained), compared to day 1 (early) (**Fig. S2E**), while consistently consuming sucrose on all trials (**Fig. S2E**). Collectively, these features were well represented in their operant index, showing an increase across days signifying learning (**Fig. 2D**). Following this, mice underwent extinction training over 5 days, during which sucrose delivery upon an active poke was omitted on all trials. Mice rapidly learned to decrease their active nose poke operant responses and approaches to the sucrose receptacle across 5 days of extinction, resulting in a reduction in their operant index (**Fig. 2D, S2E**).

**FIGURE 2.**
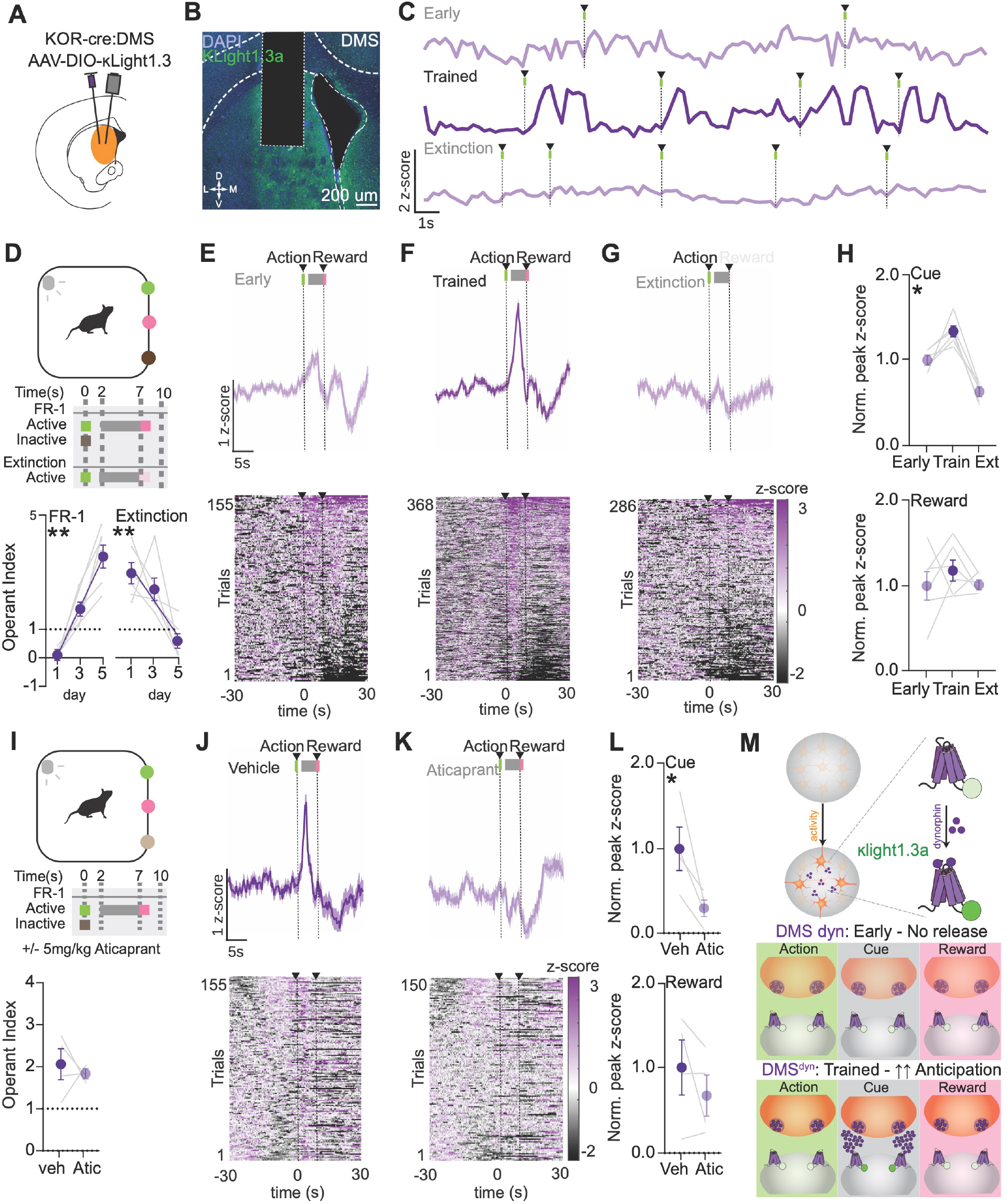
DMS dynorphin is released upon cued anticipation of reward during goal-directed behavior. (**A**) Schematic of viral injection and optic fiber implantation in the DMS. (**B**) 20X Confocal image of optic fiber implant in the DMS and cells stained with DAPI, expressing kLight 1.3a. (**C**) Mean Fluorescence during early (top) and trained (bottom) operant behavior time-locked to reinforced active nosepokes from a representative animal. (**D**) Top - Operant behavior schedule during photometry. Bottom - Operant learning and extinction. Left – Operant Learning (n=6mice; Simple Linear Regression, R^2^=0.8382, *p<0*.*0001*, F=82.86). Right – Operant Extinction (n=6mice; Simple Linear Regression, R^2^=0.5800, *p=0*.*0002*, F=22.10). (**E**) Mean fluorescence and heatmap raster plots during early operant behavior (n=6mice). (**F**) Mean fluorescence and heatmap raster plots during trained operant behavior (n=6mice). (**G**) Mean fluorescence and heatmap raster plots during operant extinction behavior (n=6mice). (**H**) Top - Normalized Peak z-score values of early, trained and extinction during **cue** period (2-7s) (n=6mice; One Way ANOVA, *p<0*.*0001*\*\*\*\*, F(1.863,9.314)=38.24. Multiple comparisons – early vs. trained, *p=0*.*0307\**, trained vs. extinction, *p=0*.*0006\*\*\**, early vs. extinction, *p=0*.*0094\*\**). Bottom - Normalized Peak z-score values of early, trained and extinction during **reward** period (7-30s) (n=6mice; One Way ANOVA, *p>0*.*05*). (**I**) Top - Operant behavior schedule during photometry and aticaprant injection. Bottom - Operant behavior during vehicle vs. Aticaprant (n=4mice; paired t test, *p>0*.*05*). (**J**) Mean fluorescence and heatmap raster plots following vehicle injection during operant behavior (n=4mice). (**K**) Mean fluorescence and heatmap raster plots following aticaprant injection during operant behavior (n=4mice). (**L**) Top - Normalized Peak z-score values of vehicle vs. Aticaprant during **cue** period (2-7s) (n=4mice; paired t test, *p=0*.*0457\**, t=3.301, df=3). Bottom - Normalized Peak z-score values of vehicle vs. Aticaprant during **reward** period (7-30s) (n=4mice; paired t test, *p>0*.*05*). (**M**) Summary schematic of interpretation of results.

Simultaneously during goal-directed behavior, we measured kLight-dependent fluorescence across the learning of the task. We observed no appreciable changes in fluorescence during behavioral events on day one of learning (**Fig. 2E,H**); however, once the animals showed an operant index significantly greater than 1, we observed a significant increase in dyn release after mice nosepoked and approached the reward port, during the cue period (**Fig. 2F,H**). Furthermore, this fluorescence significantly diminished following extinction learning, even though mice continued to approach the reward receptacle (**Fig. 2G,H**). To determine selectivity of kLight-dependent fluorescence as dynorphin-sensitive, we injected mice with either vehicle or aticaprant (KOR antagonist, i.p., 5 mg/kg) as they engaged in operant behavior. Although there were no changes in the maintenance of goal-directed behavior as has previously been observed with systemic injections of KOR antagonists (Farahbakhsh et al. 2023) (**Fig. 2I, S2F**), we show that KOR blockade eliminated fluorescence dynamics (**Fig. 2J-L**), suggesting that fluctuations in fluorescent activity were kLight-dependent. Interestingly, as mice consumed more rewards following cue delivery during Pavlovian conditioning (**Fig. S2E**), we did not observe an increase in dyn release during the cue period (**Fig. S2G-J**). This suggests that the dyn release in the DMS is specific to goal-directed action-outcome behavior, and thus may relate specifically to shaping action-outcome associations. Altogether, our results show that dyn release in the DMS evolves across days of operant learning, and is locally released within the timescale of seconds, specifically following action and a cued anticipation for the reward. (**Fig. 2M**).

### DMS dynorphin is necessary and sufficient for acquiring goal-directed behavior

To whether DMS dynorphin release is required for acquiring goal-directed behavior, we determined the contribution of dynorphin release during anticipation of reward following an operant response (**Fig. 3A**). We injected Pdyn-cre (DMS^pdyn^) or WT mice with AAV5-EF1a-DIO-ChR2-YFP and implanted them with a 200 mm optic fiber (**Fig. 3B, S3A**). Following training on the Pavlovian task and then operant conditioning while tethered to a fiber optic cable, mice received 20 Hz, 5 ms pulse width, 465 nm light stimulation for 7 seconds following an active nose-poke, terminating at reward delivery, to mimic the period of evoked klight fluorescent activity we previously observed (**Fig 2**). DMS^pdyn^ mice showed a significant increase in their operant index compared to WT upon photo-stimulation during the reward delivery window (**Fig. 3C, S3B**). Importantly, sucrose-naïve DMS^pdyn^ or WT mice did not display reinforcement behavior when nosepoking just for stimulation for 7 seconds (**Fig. S3B-left**); however, in accordance with prior studies using D1-cre mice (Vicente et al. 2016, Lalive et al. ????), 1 second stimulation triggered by the nosepoke itself did produce robust self-stimulation behavior (**Fig. S3B-right**). To further measure whether the sufficiency for pdyn neuron stimulation emerged following operant conditioning for sucrose, we split the DMS^pdyn^ mice into two groups for extinction – one that received photo-stimulation of DMS^pDyn^ neurons during reward omission and the other that did not. We found that the group that received photo-stimulation maintained their operant responding despite the lack of any reward delivery over multiple days (**Fig. 3D-left, S3D**). Upon switching the two groups, we observed that the group that previously received stimulation readily extinguished operant behavior, while the group that displayed prior extinction now regained their operant responding (**Fig. 3D-right, S3D**). Finally, to determine whether activation of KOR mediated this effect, we i.p injected DMS^pdyn^ mice with the KOR antagonist aticaprant (5 mg/kg) and found that this eliminated the increase in operant index upon stimulation (**Fig. 3E**). Interestingly, this effect was largely mediated by a reduction in reward consumption as animals still increased their active nosepokes upon stimulation following KOR antagonism (**Fig. S3E**), suggesting that dyn release and subsequent dyn-KOR following an action during the cued anticipation of a reward (like observed in **Fig. 2**) is essential for sustaining goal-directed behavior.

**FIGURE 3.**
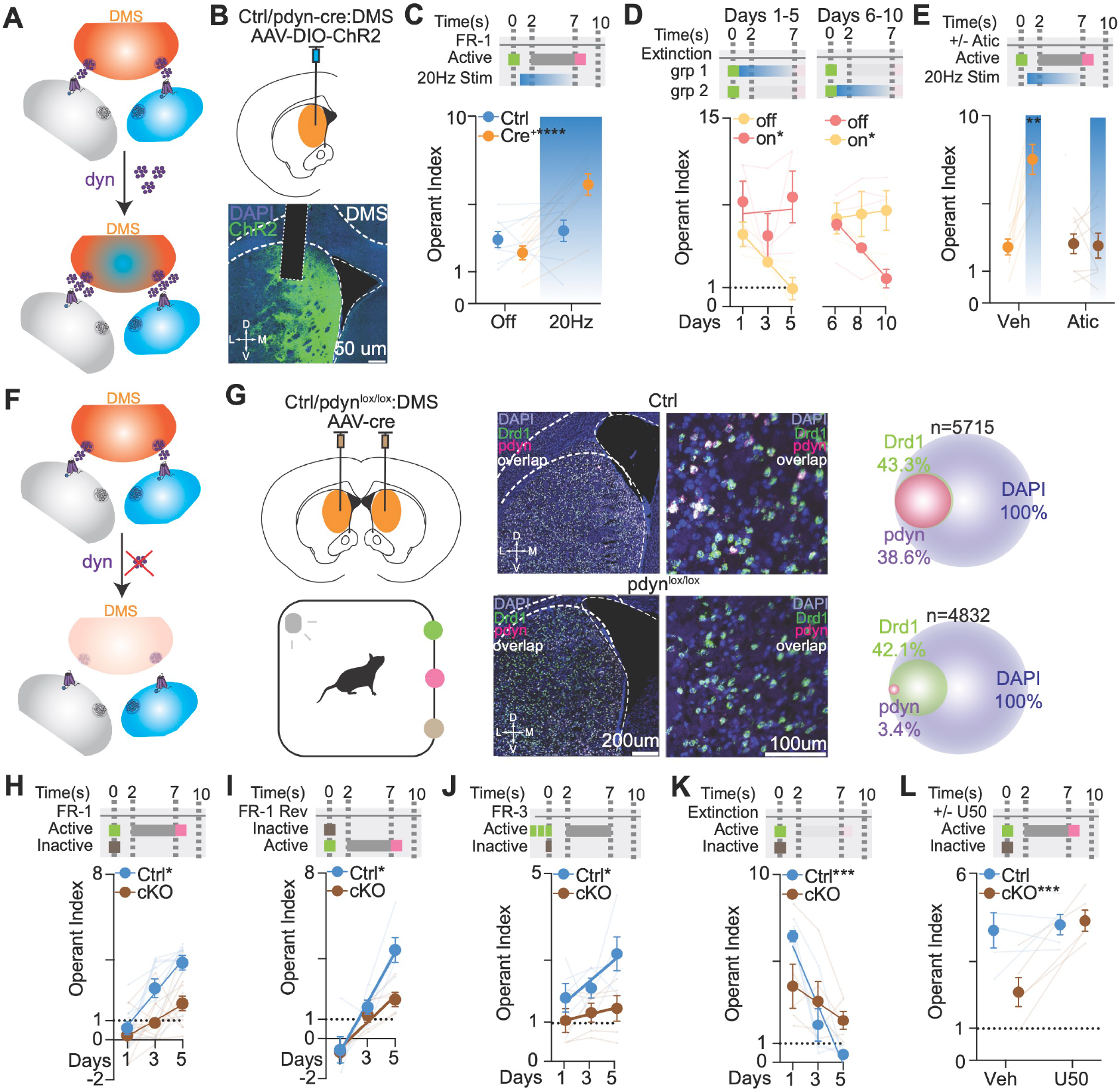
DMS dynorphin is sufficient and necessary for goal-directed behavior. (**A**) Schematic of experimental design to test whether dyn release from DMS dyn neurons via optogenetics is sufficient for goal-directed behavior. (**B**) Top - Schematic of viral injection and optic fiber implantation in the DMS. Bottom - 20X Confocal image of optic fiber implant in the DMS and cells stained with DAPI, expressing ChR2. (**C**) Operant behavior during optogenetic stimulation in Ctrl and DMS^pdyn^ mice (n=8 Ctrl,8 dyn-cre mice; Two Way ANOVA, stim x genotype *p<0*.*0009\*\*\**, F(1,14)=17.75. Multiple comparisons – ctrl off vs. on, *p>0*.*05;* dyn-cre off vs. on, *p<0*.*0001\*\*\*\**). (**D**) Left - Extinction behavior during days 1-5 (n=4mice; Off - Simple Linear Regression, R^2^=0.6401, *p=0*.*0018\*\**, F=17.78. On – Simple Linear Regression, R^2^=0.001827, *p>0*.*05*, F=0.01830. Difference in intercepts – *p=0*.*0035*^*#*^). Right - Extinction behavior during days 6-10 (n=4mice; Off - Simple Linear Regression, R^2^=0.6401, *p<0*.*0001\*\*\*\**, F=39.05. Difference in Slopes – *p=0*.*0356\**. On – Simple Linear Regression, R^2^=0.008760, *p>0*.*05*, F=0.08837). (**E**) Operant behavior during optogenetic stimulation in DMS^pdyn^ mice with aticaprant injection (n=8 dyn-cre mice; Two Way ANOVA, stim x treatment *p<0*.*0059\*\**, F(1,7)=15.18. Multiple comparisons – veh off vs. on, *p=0*.*0007***;* aticaprant off vs. on, *p>0*.*05*). (**F**) Schematic of experimental design to test whether dyn in the DMS is necessary for goal-directed behavior. (**G**) Left - Schematic of viral injection in the DMS and schematic of operant behavior. Right top - 20X (left) and 40X (middle) Confocal image of **Control** DMS section with ISH for DAPI, DRD1, Pdyn and overlap, and (right) Quantification of ISH. Right bottom - 20X (left) and 40X (middle) Confocal image of **DMS**^**pdyn-cKO**^ DMS section with ISH for DAPI, DRD1, Pdyn and overlap, and (right) Quantification of ISH. (**H**) Operant learning (n=9 Ctrl, 12 DMS^pdyn-cKO^ mice; Ctrl - Simple Linear Regression, R^2^=0.5782, *p<0*.*0001\*\*\*\**, F=34.27. DMS^pdyn-cKO^ – Simple Linear Regression, R^2^=0.2808, *p=0*.*0009\*\*\**, F=13.27. Difference in Slopes – *p=0*.*0229\**). (**I**) Operant Reversal learning (n=5 Ctrl, 5 DMS^pdyn-cKO^ mice; Ctrl - Simple Linear Regression, R^2^=0.7663, *p<0*.*0001\*\*\*\**, F=42.63. DMS^pdyn-cKO^ – Simple Linear Regression, R^2^=0.7333, *p<0*.*0001\*\*\*\**, F=35.75. Difference in Slopes – *p=0*.*0133\**). (**J**) Operant Fixed Ratio 3 learning (n=5 Ctrl, 5 DMS^pdyn-cKO^ mice; Ctrl - Simple Linear Regression, R^2^=0.2814, *p=0*.*0419\**, F=5.091. DMS^pdyn-cKO^ – Simple Linear Regression, R^2^=0.04635, *p>0*.*05*, F=0.6319. Difference in Intercepts – *p=0*.*0029*^*#*^). (**K**) Operant Extinction learning (n=5 Ctrl, 5 DMS^pdyn-cKO^ mice; Ctrl - Simple Linear Regression, R^2^=0.7928, *p<0*.*0001\*\*\*\**, F=49.75. DMS^pdyn-cKO^ – Simple Linear Regression, R^2^=0.1324, *p>0*.*05*, F=1.984. Difference in Slopes – *p=0*.*0086\*\**). (**L**)Operant behavior in Ctrl and DMS^pdyn-cKO^ mice with U50,488 injection (n=5 Ctrl, 5 DMS^pdyn-cKO^ mice; Two Way ANOVA, genotype x treatment *p<0*.*0037\*\**, F(1,8)=16.41. Multiple comparisons – Ctrl veh vs. U50, *p>0*.*05;* DMS^pdyn-cKO^ veh vs. U50, *p=0*.*0003\*\*\**).

Next, we determined the necessity for dyn during acquiring goal-directed behavior in our task (**Fig. 3F**). Pdyn from the DMS was knocked out in Pdyn^lox/lox^ or WT mice were injected with AAV5-Cre recombinase (DMS^pdyn-cKO^)(**Fig. 3G**). Surprisingly, although DMS^pdyn-cKO^ mice consumed the same amount of sucrose under food-restriction in their homecage suggesting no effect on the appetitive or sucrose consumption behavior by itself (**Fig. S3G**). These mice did however consume fewer pellets during Pavlovian conditioning (**Fig. S3H**). During operant conditioning, DMS^pdyn-cKO^ mice were slower to learn goal-directed behavior as evidenced by their operant index (**Fig. 3H**), performing fewer operant actions, and consuming less rewards compared to the controls (**Fig. S3I**). To isolate whether this deficit in learning translates to other operant contingencies, mice underwent operant reversal and fixed ratio-3, where they performed 3 active nosepokes to receive a sucrose reward. Again, the DMS^pdyn-cKO^ displayed deficits in their operant index in both operant reversal learning (**Fig.3I, S3J**) and fixed ratio-3 responding (**Fig.3J, S3K**). Additionally, we observed that DMS^pdyn-cKO^ mice extinguished their operant responding slower than control animals during extinction (**Fig.3K, S3L**). Strikingly, an i.p injection of U50, 488 (5 mg/kg) to engage KOR signaling rescued the deficits in operant responding, rewards consumed and subsequently, the operant index in DMS^pdyn-cKO^ mice (**Fig.3L, S3M**) while behavior in control mice remained unaffected. This result indicates that dynorphin tone engages KOR to maintain goal-directed behaviors. Altogether, these results demonstrate that local dyn-KOR signaling in the DMS is necessary and sufficient for the learning, maintenance and extinction of goal-directed action-outcome behavior.

### BLA KOR-expressing neurons project onto DMS D1/dyn neurons and are necessary for goal-directed behavior

To determine the locus of action of dyn-KOR signaling in the DMS, we first characterized the impact of KOR deletion from all putative inputs into the DMS. We injected KOR^lox/lox^ mice with AAV2retro-Cre recombinase in the DMS (DMS^retroKOR-cKO^). DMS^retroKOR-cKO^ mice showed deficits in goal-directed behavior compared to WT mice, specifically pertaining to behavioral learning (**Fig. S4A**), even though they consumed the same number of rewards as controls in their homecage and during Pavlovian conditioning (**Fig. S4D,E**). Interestingly, DMS^retroKOR-cKO^ mice displayed heightened action vigor during the early sessions of operant conditioning but did not improve their action-outcome behavior across learning, like controls did (**Fig. S4F**).

To determine circuit-specific contributions of dyn-KOR signaling, we injected WT mice with AAV2retro-Cre recombinase and used fluorescent in situ hybridization in regions that project to the DMS (**Fig. 4A**). We found that over 50% of all neurons in the BLA express KOR mRNA; upon determining the levels of Cre mRNA in the BLA in this experiments, Cre mRNA was found largely in CamKII+ neurons with 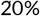 of BLA neurons also expressing Cre mRNA. Interestingly, over 60% of Cre+ neurons also expressed KOR mRNA (**Fig. 4A-top**). Additionally, we found that over 80% of Cre+ neurons also expressed Vglut1 mRNA (**Fig. 4A-bottom)**, which is the predominant vesicular glutamate transporter found in the BLA (Fremeau *et al*., 2001). Anterograde viral tracing in either KOR-Cre or Vglut1-Cre mice injected with AAV5-EF1a-DIO-ChR2-YFP showed YFP+ terminals in the DMS from both BLA KOR- (**Fig. 4B-top**) and Vglut1-expressing neurons (**Fig. 4B-bottom**), further corroborating our findings.

**FIGURE 4.**
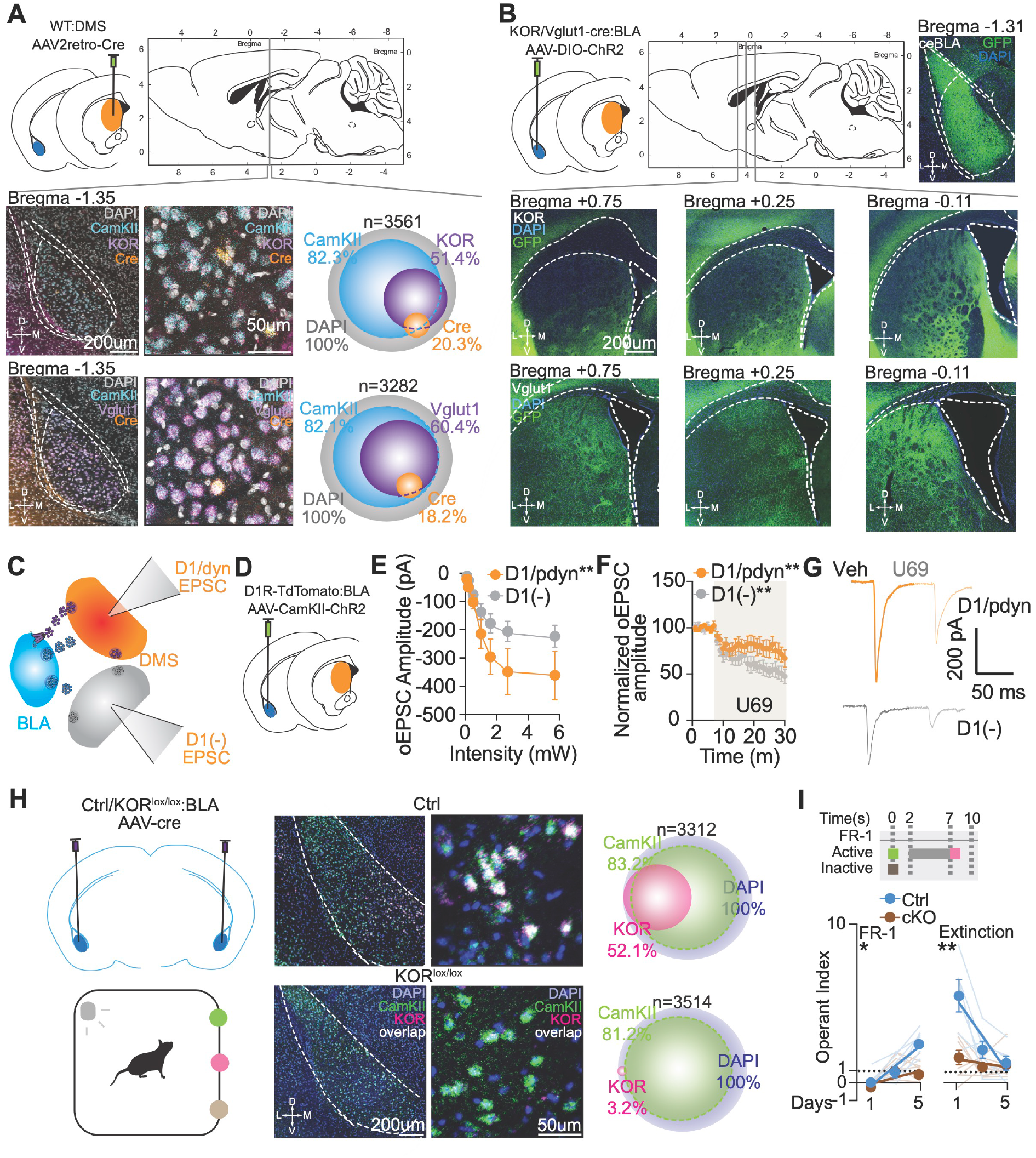
BLA KOR-expressing neurons preferentially project to DMS D1/dyn neurons and are necessary for goal-directed behavior. (**A**) Top - Schematic of viral injection in the DMS. Middle - 20X (left) and 40X (middle) Confocal image of BLA section with ISH for DAPI, CamKII, KOR and Cre, and (right) Quantification of ISH. Bottom - 20X (left) and 40X (middle) Confocal image of BLA section with ISH for DAPI, CamKII, Vglut1 and Cre, and (right) Quantification of ISH. (**B**) Top - Schematic of viral injection in the BLA and 20X Confocal image of BLA section expressing ChR2. Middle - 20X Confocal image of DMS sections stained for DAPI, with KOR^+^ BLA fibers expressing ChR2 across the anterior-posterior axis. Bottom - 20X Confocal image of DMS sections stained for DAPI, with Vglut1^+^ BLA fibers expressing ChR2 across the anterior-posterior axis. (**C**) Schematic of viral injection in the BLA (**D**) Schematic of ex vivo electrophysiology from DMS^pdyn^ and D1(−) neurons. (**E**) Input-output curve of optically-evoked EPSCs from DMS^pdyn^ and D1(−) DMS neurons (n=16 cells, 5 mice; Two Way ANOVA, Cell-type *p<0*.*0014\*\**, F(1,203)=10.5. Multiple comparisons – D1(+) vs. D1(−) at 2.7 mW, *p=0*.*0240*, at 5.7 mW *p=0*.*0282\**). (**F**) Normalized optically-evoked EPSCs from DMS^pdyn^ and D1(−) DMS neurons following U69 (n=8 cells, 4 mice; D1(+) - paired t test, *p<0*.*0313\**, t=2.686, df=7; D1(−) - paired t test, *p<0*.*0002\*\*\**, t=6.599, df=7). (**G**) Representative traces for Veh and U69 in DMS^pdyn^ (orange) and D1(−) (grey) neurons. (**H**) Left - Schematic of viral injection in the BLA and schematic of operant behavior. Right top - 20X (left) and 40X (middle) Confocal image of **Control** BLA section with ISH for DAPI, CamKII, KOR and overlap, and (right) Quantification of ISH. Right bottom - 20X (left) and 40X (middle) Confocal image of **BLA**^**KOR-cKO**^ BLA section with ISH for DAPI, CamKII, KOR and overlap, and (right) Quantification of ISH. (**I**) Left – Operant learning (n=6 Ctrl, 8 BLA^KOR-cKO^ mice; Ctrl - Simple Linear Regression, R^2^=0.6587, *p<0*.*0001\*\*\*\**, F=30.89. BLA^KOR-cKO^ – Simple Linear Regression, R^2^=0.2152, *p=0*.*0224\**, F=6.033. Difference in Slopes – *p=0*.*0043\*\**). Right – Extinction learning (n=6 Ctrl, 8 BLA^KOR-cKO^ mice; Ctrl - Simple Linear Regression, R^2^=0.5057, *p=0*.*0009\*\*\**, F=16.37. BLA^KOR-cKO^ – Simple Linear Regression, R^2^=0.0440, *p>0*.*05*, F=1.013. Difference in Slopes – *p=0*.*001\*\**).

To further isolate if BLA terminals are functionally connected to the DMS, we performed *ex vivo* whole-cell patch clamp electrophysiology experiments (**Fig. 4C**). We injected D1R-TdTomato mice with AAV5-CamKIIa-ChR2-YFP in the BLA to stimulate BLA terminals and record optically evoked EPSCs from either D1/dyn (DMS^pdyn^) or D1(−) neurons (**Fig. 4D**). We found that BLA terminal stimulation evoked oEPSCs in both populations, albeit with a significantly higher optically-evoked amplitude in DMS^pdyn^ neurons (**Fig. 4E**). Furthermore, to measure if these terminals are directly modulated by KOR, we bath applied the KOR agonist U69 and found that U69 inhibited oEPSCs equivalently from both neuronal populations (**Fig. 4F,G**). Given the known role of Gi-coupled GPCRs in coupling to inhibition of transmitter release via beta-gamma-mediated calcium channel inhibition (Al-Hasani and Bruchas 2011, Corder et al. 2018), this result suggests that pre-synaptic KOR acts to negatively regulate glutamatergic release from BLA inputs to the DMS.

Next, we asked what selectively removing KOR from the BLA would do to performance of animals in goal-directed behavior. We injected KOR^lox/lox^ mice with AAV5-Cre recombinase in the BLA (BLA^KOR-cKO^) (**Fig. 4H**) and observed that with a similar effect of to DMS^pdyn-cKO^ (**Fig 3**), these mice consumed the same amount of sucrose in the homecage (**Fig. S4H**), yet consumed fewer rewards during Pavlovian conditioning (**Fig. S4I**). Moreover, BLA^KOR-cKO^ mice were slower to learn, sustain goal-directed behavior and extinguished their operant responding with a slower rate as compared to the control group (**Fig. 4I, S4J,K**). These results indicate that glutamatergic neurons from the BLA terminals in the DMS are regulated by presynaptic KORs which are necessary for maintaining goal-directed behavior.

### BLA-DMS terminals are necessary and sufficient for goal-directed behavior

Our results suggest a mechanism of regulation of BLA-DMS terminals during goal-directed behavior wherein: (i) BLA terminal activation of DMS dyn neurons during action causes (ii) retrograde DMS dyn release during anticipation of reward, causing (iii) dyn-KOR negative modulation and Gi-coupled GPCR mediated inhibition of glutamate release at BLA terminals, thereby promoting goal-directed behavior (**Fig. 5A, hypothesis**). To test this hypothesis in vivo, we first measured the activity of Vglut1-expressing BLA-DMS terminals (BLA^vglut1^-DMS) using fiber photometry, by injecting Vglut1-cre mice with AAVDJ-EF1a-DIO-GCaMP6s and implanting a 400 um optic fiber in the DMS (**Fig. 5B, S5A**). As mice progressed through Pavlovian conditioning (**Fig. S5B**), we observed a significant inhibition of BLA^vglut1^-DMS fluorescence upon reward delivery (**Fig. S5C-E**). As animals learned operant conditioning (**Fig. 5D**), we observed a similar reduction in fluorescence during reward (**Fig. 5C,E**), but strikingly, a significant ramping of GCaMP activity as the animals performed nosepokes to obtain rewards (**Fig. 5C,E,G**). Interestingly, we also found a significant reduction in BLA^vglut1^-DMS fluorescence at the time of cue delivery (**Fig. 5G**), following action that we previously did not observe during Pavlovian conditioning (**Fig. S5E**). Importantly, the activation of BLA^vglut1^-DMS fluorescence during action and inhibition during cue and reward were eliminated during extinction, where reward delivery was omitted (**Fig. 5F,G**).

**FIGURE 5.**
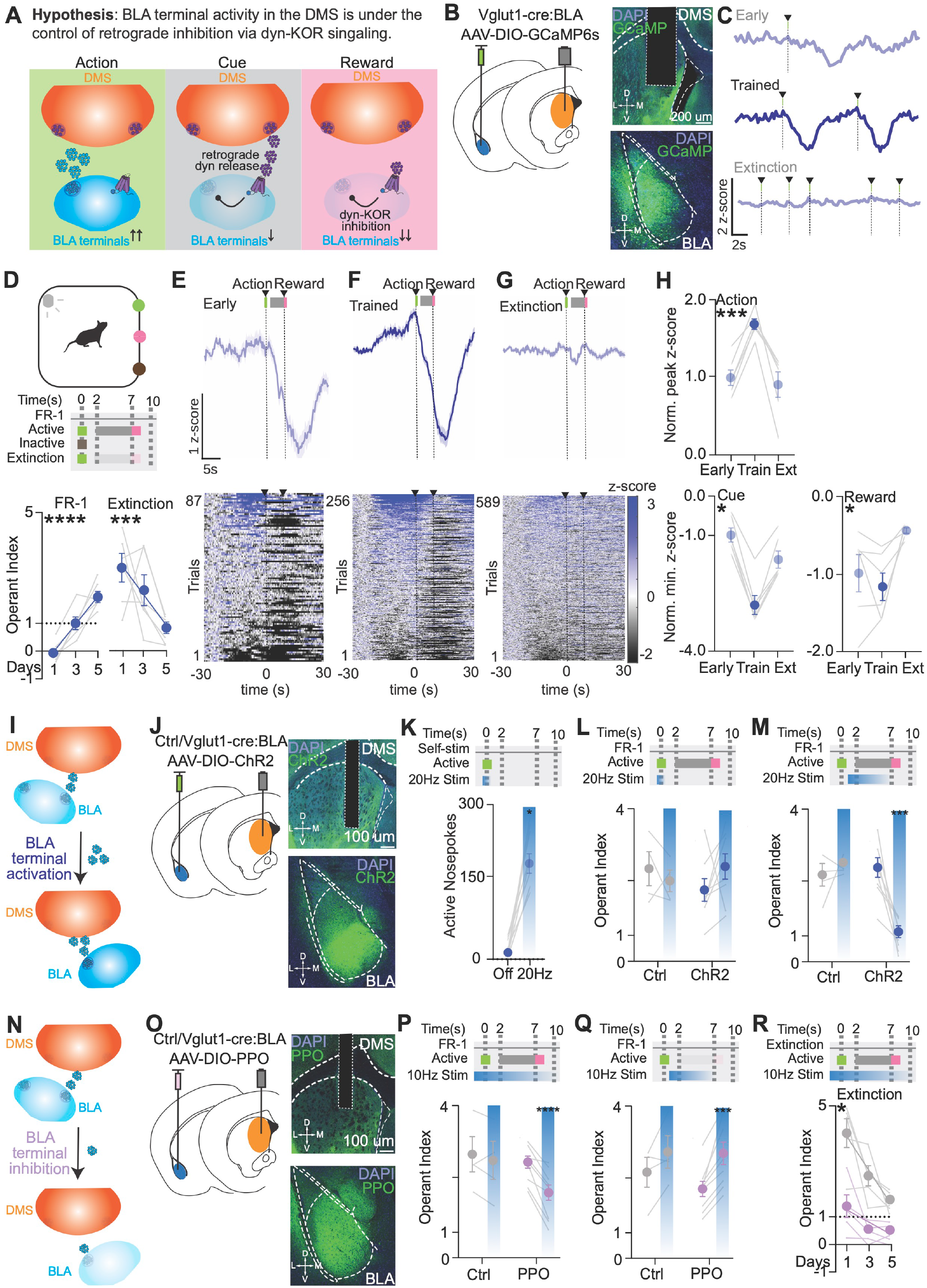
BLA-DMS terminals are engaged during and necessary, and sufficient for goal-directed behavior. (**A**) Schematic of experimental hypothesis to test whether dyn-KOR signaling at BLA-DMS terminals promotes goal-directed behavior. (**B**) Left - Schematic of viral injection in the BLA and optic fiber implantation in the DMS. Right top - 20X Confocal image of optic fiber implant in the DMS and cells stained with DAPI, with Vglut1 terminals expressing GCaMP6s. Right bottom - 20X Confocal image of BLA and cells stained with DAPI, with Vglut1 cells expressing GCaMP6s. (**C**) Mean Fluorescence during early (top) and trained (bottom) operant behavior time-locked to reinforced active nosepokes from a representative animal. (**D**) Top - Operant behavior schedule during photometry. Bottom - Operant learning and extinction. Left – Operant Learning (n=6mice; Simple Linear Regression, R^2^=0.7781, *p<0*.*0001\*\*\*\**, F=56.11). Right – Operant Extinction (n=6mice; Simple Linear Regression, R^2^=0.4251, *p=0*.*0033\*\**, F=11.83). (**E**) Mean fluorescence and heatmap raster plots during early operant behavior (n=6mice). (**F**) Mean fluorescence and heatmap raster plots during trained operant behavior (n=6mice). (**G**) Mean fluorescence and heatmap raster plots during operant extinction behavior (n=6mice). (**H**) Top – Normalized Peak z-score values of early, trained and extinction during **action** period (−20-0s) (n=6mice; One Way ANOVA, *p=0*.*0026*\*\*, F(1.307,6.535)=19.94. Multiple comparisons – early vs. trained, *p=0*.*0005\*\*\**, trained vs. extinction, *p=0*.*0092\*\**, early vs. extinction, *p>0*.*05*). Bottom Left – Normalized Minimum z-score values of early, trained and extinction during **cue** period (2-7s) (n=6mice; One Way ANOVA, *p=0*.*0002*\*\*\*, F(1.454,7.272)=42.8. Multiple comparisons – early vs. trained, *p=0*.*0002\*\*\**, trained vs. extinction, *p=0*.*0027\*\**, early vs. extinction, *p>0*.*05*). Bottom Right – Normalized Minimum z-score values of early, trained and extinction during **reward** period (7-30s) (n=6mice; *p=0*.*0222*\*, F(1.649,8.245)=6.648. Multiple comparisons – early vs. trained, *p>0*.*05*, trained vs. extinction, *p=0*.*0269\**, early vs. extinction, *p>0*.*05*). (**I**) Schematic of experimental design to test whether BLA-DMS activity via optogenetics is sufficient for goal-directed behavior. (**J**) Left - Schematic of viral injection in the BLA and optic fiber implantation in the DMS. Right top - 20X Confocal image of optic fiber implant in the DMS and cells stained with DAPI, with Vglut1 terminals expressing ChR2. Right bottom - 20X Confocal image of the BLA and cells stained with DAPI, with Vglut1 cells expressing ChR2. (**K**) Active nosepokes for 1 second self-stimulation (n=8 vglut1-cre mice; paired t test, *p<0*.*0001\*\*\*\**, t=8.548, df=7). (**L**) Operant behavior during optogenetic stimulation at **action** in Ctrl and vglut1-cre mice (n=4 Ctrl,8 vglut1-cre mice; Two Way ANOVA, stim x genotype *p>0*.*05*, F(1,10)=1.662). (**M**) Operant behavior during optogenetic stimulation at **cue and reward** in Ctrl and Vglut1-cre mice (n=4 Ctrl,8 Vglut1-cre mice; Two Way ANOVA, stim x genotype *p=0*.*005\*\**, F(1,10)=12.84. Multiple comparisons – ctrl off vs. on, *p>0*.*05;* vglut1-cre off vs. on, *p=0*.*0008\*\*\**). (**N**) Schematic of experimental design to test whether BLA-DMS activity via optogenetics is necessary for goal-directed behavior. (**O**) Left - Schematic of viral injection in the BLA and optic fiber implantation in the DMS. Right top - 20X Confocal image of optic fiber implant in the DMS and cells stained with DAPI, with Vglut1 terminals expressing PPO. Right bottom - 20X Confocal image of the BLA and cells stained with DAPI, with Vglut1 cells expressing PPO. (**P**) Operant behavior during optogenetic inhibition at **whole session** in Ctrl and vglut1-cre mice (n=4 Ctrl,9 vglut1-cre mice; Two Way ANOVA, stim x genotype *p=0*.*0079\*\**, F(1,11)=10.48. Multiple comparisons – ctrl off vs. on, *p>0*.*05;* vglut1-cre off vs. on, *p<0*.*0001\*\*\*\**). (**Q**) Operant behavior during optogenetic inhibition at **cue and reward** in Ctrl and vglut1-cre mice (n=4 Ctrl,9 vglut1-cre mice; Two Way ANOVA, *stim p=0*.*0005\*\*\**, F(1,11)=23.95, stim x genotype *p>0*.*05*. Multiple comparisons – ctrl off vs. on, *p>0*.*05;* vglut1-cre off vs. on, *p=0*.*0002\*\*\**). (**R**) Extinction behavior during optogenetic inhibition at **whole session** (n=5 vglut1-cre mice; Off - Simple Linear Regression, R^2^=0.6151, *p=0*.*0005\*\*\**, F=20.77. On – Simple Linear Regression, R^2^=0.2822, *p=0*.*0416\**, F=5.11. Difference in Slopes – *p=0*.*0261\**).

Since BLA^vglut1^-DMS terminals displayed a bimodal pattern of activity during an action-outcome sequence, we then attempted to parse each of these differing contributions to a respective behavior via optogenetic manipulation (**Fig. 5I,N**). First, we injected Vglut1-cre mice with AAVDJ-EF1a-DIO ChR2-YFP in the BLA and implanted 200 um optical fibers in the DMS to photo-activate BLA^vglut1^-DMS terminals (**Fig. 5I,J, S5A**). Remarkably, sucrose- and operant-naïve mice nosepoked repeatedly just for BLA^vglut1^-DMS terminal stimulation, suggesting that their activity is reinforcing (**Fig. 5K**). We then trained these mice on operant conditioning, following which mice either received 20 Hz, 5 ms pulse-width, 465nm light delivery into the DMS triggered by the nosepoke to enhance BLA^vglut1^-DMS activity, or during cue and reward delivery, to disrupt operant behavior. Surprisingly, although animals receiving nosepoke-triggered stimulation made more active nosepokes (**Fig. S5F**), they consumed the same number of rewards, resulting in no net difference in their operant index (**Fig. 5L**). Conversely, stimulating BLA^vglut1^-DMS terminals during cue and reward delivery to disrupt the decrease in BLA^vglut1^-DMS activity, resulted in a reduction in their operant index (**Fig. 5M, S5G**). In parallel, we injected Vglut1-Cre mice with Cre-dependent parapinopsin (PPO) (Copits et al. 2021) in the BLA and implanted 200mm optical fibers in the DMS to spatiotemporally mimic Gi-coupled GPCR inhibition of BLA^vglut1^-DMS terminals using optogenetics (**Fig. 5N,O, S5A**). Optical inhibition during the entire session resulted in a significant reduction in the operant index (**Fig. 5P, S5G**). Furthermore, inhibition of BLA^vglut1^-DMS terminals during cue and reward delivery to mimic the KOR-mediated reduction in their activity (**Fig 4**) caused a significant increase in operant behavior (**Fig. 5Q, S5I**). Next, we asked whether BLA^vglut1^-DMS terminal inhibition upon cue and reward delivery during extinction impacts mice’s ability to extinguish operant behavior. We found that mimicking terminal Gi/o-mediated inhibition significantly decreased extinction behavior in PPO animals compared to controls (**Fig. 5R, S5J**). Altogether, our results show that BLA^vglut1^-DMS terminal activity is engaged upon learning, necessary and sufficient for goal-directed behavior, and is under the control of Gi/o GPCR-mediated inhibition during outcome, thereby invigorating action.

### Dynorphin-KOR signaling at BLA-DMS terminals is necessary for acquiring and maintaining goal-directed behavior

To determine whether BLA-DMS terminal activity is under dyn-KOR control, we first i.p injected Vglut1-Cre mice with a low dose of the non-selective opioid receptor antagonist naloxone (2mg/kg; higher doses result in a reduction in sucrose consumption; (Castro et al. 2021)) while monitoring BLA^vglut1^-DMS fluorescence using fiber photometry from the same mice in **Fig. 5** (**Fig. 6A**). Opioid receptor antagonism at this dose did not impact operant behavior (**Fig. 6B, S6B**), as we previously observed with the KOR antagonist aticaprant (**Fig. 2K, S2I**). However, we did observe a significant reduction in BLA^vglut1^-DMS fluorescence dynamics (**Fig. 6C,D**) wherein naloxone injected animals showed a decrease in the magnitude of GCaMP activity during action and reduced inhibition during reward compared to when they were treated with the vehicle (**Fig. 6E**). Next, we tested the contributions of dyn expression in DMS neurons on BLA-DMS fluorescence during goal-directed behavior (**Fig. 6F**). We injected pdyn^lox/lox^ mice with AAV5-Cre recombinase in the DMS bilaterally to conditionally delete pdyn in DMS neurons and AAVDJ-CaMKIIa-GCaMP6s in the BLA, and implanted a 400 mm optic fiber in the DMS unilaterally (BLA^CaMKII^-DMS^pdyn-cKO^) (**Fig. 6F, S6E**). Corroborating our prior studies (**Fig. S3G,H**), mice lacking pdyn consumed the same amount of sucrose under food-restriction in their homecage, they consumed fewer pellets during Pavlovian conditioning (**Fig. S6F**). Additionally, we saw a similar inhibition of BLA^CaMKII^-DMS fluorescence upon reward delivery (**Fig. S6G-I**), as we previously observed in BLA^vglut1^-DMS terminals *in vivo* (**Fig. S5C-E**). During operant conditioning, we observed similar prior deficits (**Fig. 3H, S3I**) in goal-directed behavior in these BLA^CaMKII^-DMS^pdyn-cKO^ mice relative to controls (**Fig. 6G, S6J**). Strikingly, we found a concomitant drastic reduction in the magnitude of BLA^CaMKII^-DMS^pdyn-cKO^ fluorescence to both action and outcome (**Fig. 6H-J**), suggesting that the elimination of dyn from the DMS significantly impacted BLA^CaMKII^-DMS activity. Collectively, these results lead us to conclude that dyn-KOR signaling at BLA-DMS terminals is necessary for promoting directed time-locked modulation of their activity to consequently shape the learning and maintenance of goal-directed behavior.

**FIGURE 6.**
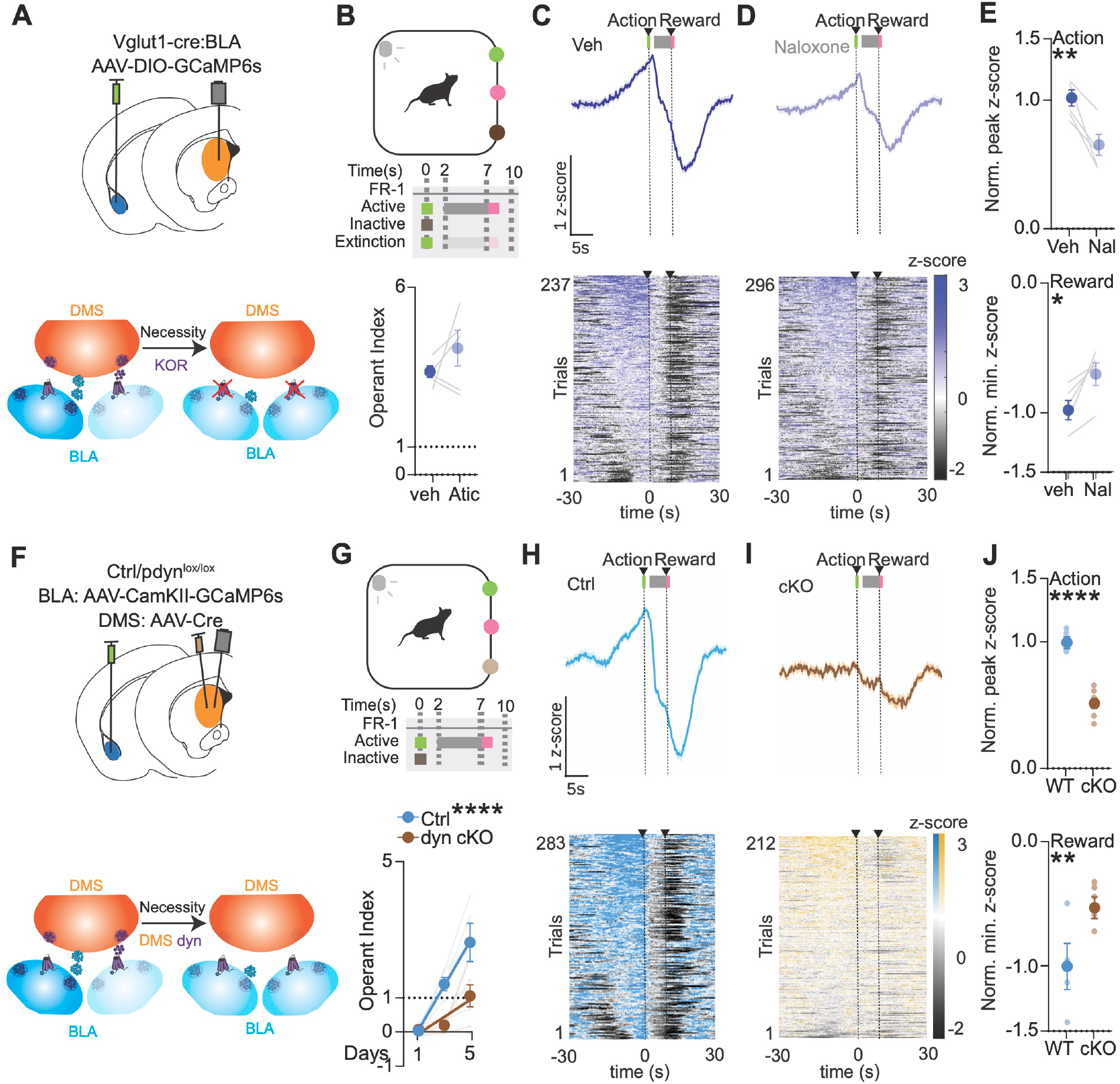
Dynorphin-KOR signaling at BLA-DMS terminals is necessary for goal-directed behavior. (**A**) Top - Schematic of viral injection in the BLA and optic fiber implantation in the DMS. Bottom - Schematic of experimental hypothesis to test whether KOR signaling at BLA-DMS terminals is necessary for goal-directed behavior. (**B**) Top - Operant behavior schedule during photometry and naloxone injection. Bottom - Operant behavior during vehicle vs. Naloxone (n=5mice; paired t test, *p>0*.*05*). (**C**) Mean fluorescence and heatmap raster plots following vehicle injection during operant behavior (n=5mice). (**D**) Mean fluorescence and heatmap raster plots following naloxone injection during operant behavior (n=5mice). (**E**) Top - Normalized Peak z-score values of vehicle vs. Naloxone during **action** period (−20-0s) (n=5mice; paired t test, *p=0*.*0092\*\**, t=4.632, df=4). Bottom - Normalized Minimum z-score values of vehicle vs. Naloxone during **reward** period (7-30s) (n=5mice; paired t test, *p=0*.*0201\**, t=3.744, df=4). (**F**) Top - Schematic of viral injection in the BLA and optic fiber implantation in the DMS. Bottom - Schematic of experimental hypothesis to test whether dyn release at BLA-DMS terminals is necessary for goal-directed behavior. (**G**) Top - Operant behavior schedule during photometry. Bottom - Operant learning (n=4 Ctrl, 7 DMS^pdyn-cKO^ mice; Ctrl - Simple Linear Regression, R^2^=0.8551, *p<0*.*0001\*\*\*\**, F=59. DMS^pdyn-cKO^ – Simple Linear Regression, R^2^=0.3656, *p=0*.*0037\*\**, F=10.95. Difference in Slopes – *p<0*.*0001\*\*\*\**). (**H**) Mean fluorescence and heatmap raster plots for Ctrl during trained operant behavior (n=4mice). (**I**) Mean fluorescence and heatmap raster plots for DMS^pdyn-cKO^ during trained operant behavior (n=7mice). (**J**) Top - Normalized Peak z-score values of trained Ctrl vs. DMS^pdyn-cKO^ during **action** period (−20-0s) (n=4 Ctrl, 7 DMS^pdyn-cKO^ mice; unpaired t test, *p<0*.*0001\*\*\*\**, t=7.462, df=9). Bottom - Normalized Minimum z-score values of early, trained and extinction during **reward** period (7-30s) (n=4 Ctrl, 7 DMS^pdyn-cKO^ mice; unpaired t test, *p=0*.*0077\*\**, t=3.413, df=9).

### BLA afferents induce DMS dynorphin neuron activity, retrograde dynorphin release and dynorphin-KOR signaling to sculpt goal-directed behavior

Our results thus far suggest that DMS dyn release during the anticipation of reward leads to dyn-KOR signaling at BLA-DMS terminals to shape and sustain goal-directed behaviors. To determine if DMS dyn neuron activity and subsequent dyn release is due to BLA terminal activation, we injected D1R-TdTomato mice we used in **Fig.1** with AAV9-CamKIIa-rsChRmine (rs-ChRmine) in the BLA that is activated at 1040 nm wavelength during two-photon imaging to stimulate BLA terminals while imaging activity from the DMS (**Fig. 7A**). We performed sequential, spiral stimulation at discrete points across the field of view (spirals of 5-10 mm in diameter at 20Hz, 5ms pulse-widths for 500 ms, every 10 seconds at 10 mW of laser power, sequentially tiling the entire FOV), and in counterbalanced sessions, applied the same stimulation protocol with the laser switched off, resulting only in the shutter opening (Baseline sessions). To classify neurons that were active immediately following stimulation, we grouped neurons in the stim session that displayed activity higher than that during the baseline session as “active”, and the rest as “inactive” to BLA terminal stimulation. Using this protocol, we identified 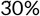 of DMS neurons that were active in response to BLA terminal stimulation, showing increases in fluorescence in the stim sessions markedly higher than during the baseline sessions (**Fig. 7A, S7A**). Further corroborating our sorting methods, collectively, “active” DMS neurons showed significantly higher activity immediately following stimulation compared to “inactive” and “baseline” (**Fig. 7B,C**). We then determined the relative proportions of these BLA-stimulated neurons based on their behavioral cluster classification during learned operant behavior. Interestingly, we found that while “active” DMS neurons were represented in each behavioral cluster (action, cue, reward and null), they were significantly enriched in the cue cluster (**Fig. 7D**). Furthermore, “inactive” DMS neurons showed no similar enrichment and were evenly distributed across all of the behavioral clusters (**Fig. S7B**). This data suggests that DMS neurons that receive BLA terminal input are preferentially activated during the cue delivery (**Fig. 7E**).

**FIGURE 7.**
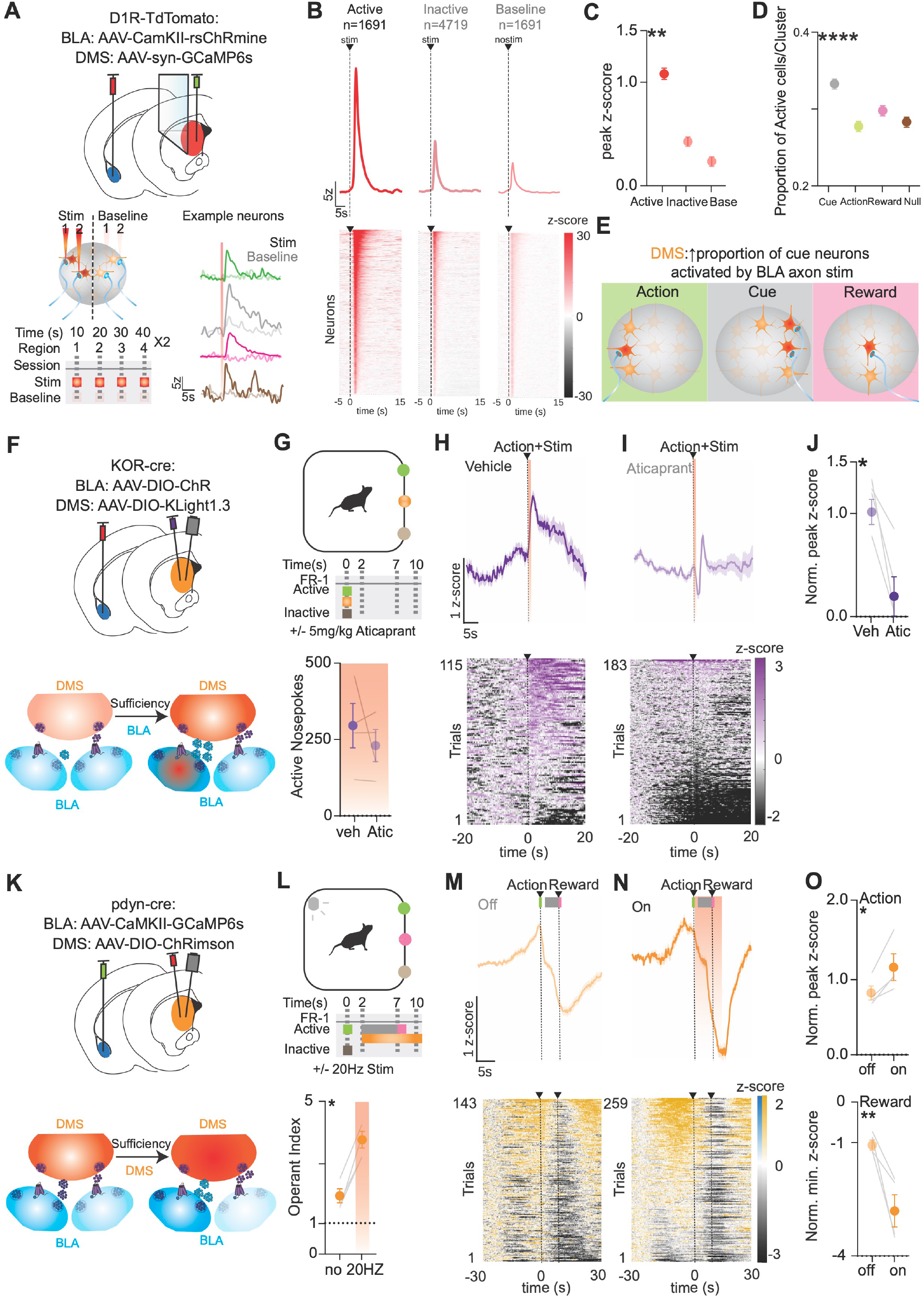
BLA terminal activity engenders DMS dynorphin neuron activity, dynorphin release and subsequent dynorphin-KOR signaling to promote goal-directed behavior. (**A**) Top - Schematic of viral injection in the BLA and DMS, and microprism implantation in the DMS. Bottom left - Schematic of experimental design to test whether BLA terminals are sufficient to activate DMS neurons. Bottom right – Traces from individual neurons in the “Stim” and “Baseline” conditions. (**B**) Classification of neurons based on activity following BLA axon stimulation showing fluorescence activity traces (top) and heat maps (bottom) of cells for “Active”, “Inactive” and “Baseline” conditions (n=2mice). (**C**) Peak z-score quantification of active, inactive and baseline neurons (n=2mice; One Way ANOVA, *p=0*.*0095\*\*\*\**. Multiple comparisons – active vs. inactive, *p=0*.*0129\**, active vs. baseline, *p=0*.*0116\**, inactive vs. baseline, *p>0*.*05*). (**D**) Proportion of active DMS cells enriched in behavioral clusters (n=2mice; One Way ANOVA, *p<0*.*0001\*\*\*\**, F(3,6406)=28.77. Multiple comparisons – Cue vs. Reward, *p<0*.*0001\*\*\*\**, Cue vs. Null, *p<0*.*0001\*\*\*\**, Action vs. Reward, *p=0*.*0057\*\**). (**E**) Summary schematic of interpretation of results. (**F**) Top - Schematic of viral injection in the BLA and optic fiber implantation in the DMS. Bottom - Schematic of experimental design to test whether BLA-DMS terminal stimulation causes dyn release. (**G**) Top – Self stimulation behavior schedule during photometry and aticaprant injection. Bottom - Operant behavior during vehicle vs. Aticaprant (n=4mice; paired t test, *p>0*.*05*). (**H**) Mean fluorescence and heatmap raster plots following vehicle injection during operant behavior (n=4mice). (**I**) Mean fluorescence and heatmap raster plots following aticaprant injection during operant behavior (n=4mice). (**J**) Normalized Peak z-score values of vehicle vs. Aticaprant following **stim** period (2-7s) (n=4mice; paired t test, *p=0*.*0186\**, t=4.662, df=3). (**K**) Top - Schematic of viral injection in the BLA and DMS, and optic fiber implantation in the DMS. Bottom - Schematic of experimental design to test whether DMS dyn stimulation modulates BLA-DMS activity. (**L**) Top – Operant behavior schedule during photometry and optogenetic stimulation. Bottom - Operant behavior during optogenetic stimulation (n=4mice; paired t test, *p=0*.*0027\**, t=9.188, df=3). (**M**) Mean fluorescence and heatmap raster plots during operant behavior with no stimulation (n=4mice). (**N**) Mean fluorescence and heatmap raster plots during operant behavior with optogenetic stimulation (n=4mice). (**O**) Top - Normalized Peak z-score values of off vs. on during **action** period (−20-0s) (n=4mice; paired t test, *p=0*.*0354\**, t=3.654, df=3). Bottom - Normalized Minimum z-score values of off vs. on during **reward** period (7-30s) (n=4mice; paired t test, *p=0*.*0059\**, t=7.039, df=3).

We then determined if BLA terminal activity was sufficient for DMS dyn release by multiplexing BLA terminal stimulation with fiber photometry to measure kLight1.3a fluorescence *in vivo* (**Fig. 7F**). We injected KOR-cre mice with AAV5-EF1a-DIO-ChRimson-TdTomato in the BLA and AAV5-EF1a-DIO-kLight1.3a in the DMS, and implanted 400 mm optic fibers (**Fig. 7F, S7C**). Since BLA^vglut1^-DMS terminal stimulation is reinforcing, we trained mice to nosepoke into an active port for 635 nm light stimulation in counterbalanced sessions, treated with vehicle or aticaprant (5 mg/kg, i.p). Mice displayed robust nosepoking for BLA^KOR^-DMS terminal stimulation in both vehicle and aticaprant conditions (**Fig. 7G**). Importantly, here we observed a significant increase in kLight fluorescence immediately following a nosepokes, as they were engaged in a nosepoke bout, and that KOR antagonism eliminated this upward change in fluorescence dynamics (**Fig. 7H-J**).

Our results thus far suggest that BLA terminal activity promotes DMS^pdyn^ activity and subsequent dyn release. Next, we ascertained its conseuqences, specifically whether DMS dyn release and aubsequent retrograde dyn-KOR signaling at BLA-DMS terminals can potentiate KOR-mediated BLA terminal inhibition and thus shape goal-directed behavior. Hence, we multiplexed DMS dyn neuron stimulation with fiber photometry to measure BLA-DMS terminal fluorescence *in vivo* (**Fig. 7K**). We injected pdyn-cre mice with AAVDJ-CamKIIa-GCaMP6s in the BLA and AAV5-EF1a-DIO-ChRimson-TdTomato in the DMS, and implanted 400 mm optic fibers (**Fig. 7K, S7D**). Before conditioning, we stimulated DMS^pdyn^ neurons for 60s with 20Hz, 635 nm wavelength light and found a significant inhibition of BLA^CamKII^-DMS terminal activity upon DMS^pdyn^ stimulation (**Fig. S7E,G**). Importantly we also found that KOR antagonism via aticaprant i.p injection was able to significantly decrease the magnitude of this inhibition (**Fig. S7F,G**). Following pavlovian and operant conditioning, we triggered 20 Hz, 5 ms pulse-width, 635nm laser stimulation for 5 seconds following active nosepokes to mimick when we previously observed dyn release (**Fig. 2**). Similar to our prior results, pdyn-cre mice enhanced their operant behavior during stimulation (**Fig. 7L, S7H**). Strikingly, DMS^pdyn^ stimulation enhanced the magnitude of BLA^CamKII^-DMS activation during action (**Fig. 7M,O**) and the magnitude of inhibition during cue, and reward (**Fig. 7N,O**). Altogether, our results lead to the conclusion that BLA terminal activity stimulates DMS dyn neuron activity, release and subsequent retrograde KOR signaling at BLA-DMS terminals to acquire and maintain goal-directed behavior. Specifically, dyn release during cued anticipation and subsequent dyn-KOR signaling during outcome potentiates the Gi/o GPCR-mediated inhibition of BLA-DMS terminals, thereby invigorating goal-directed behavior.

## 2 DISCUSSION

It is well-established that neuropeptide-GPCR signaling can shape neural activity to have a sustained impact on behavior. However, the timescales of their action and a mechanistic understanding of how they control neural activity is unclear. In this manuscript, we uncover a novel endogenous neuropeptidergic GPCR-mediated mechanism for the control of neural activity to sculpt goal-directed behavior across multiple timescales (**Fig. 8A**). As animals learn the predictive relationship between actions and outcomes, BLA terminal activity promotes the recruitment of DMS^pdyn^ neurons to encode action and cued anticipation and subsequent DMS dyn release across days (Dlearning, slow). Specifically, BLA terminal activation during action results the activation of cue-responsive DMS^pdyn^ neurons, and in dyn release during cued anticipation, thereby causing dyn-KOR signaling to inhibit BLA-DMS terminal activity during outcome (Daction-outcome, fast). Importantly, this retrograde inhibition of BLA-DMS terminal activity via dyn-KOR signaling invigorates subsequent action-outcome sequences.

**FIGURE 8.**
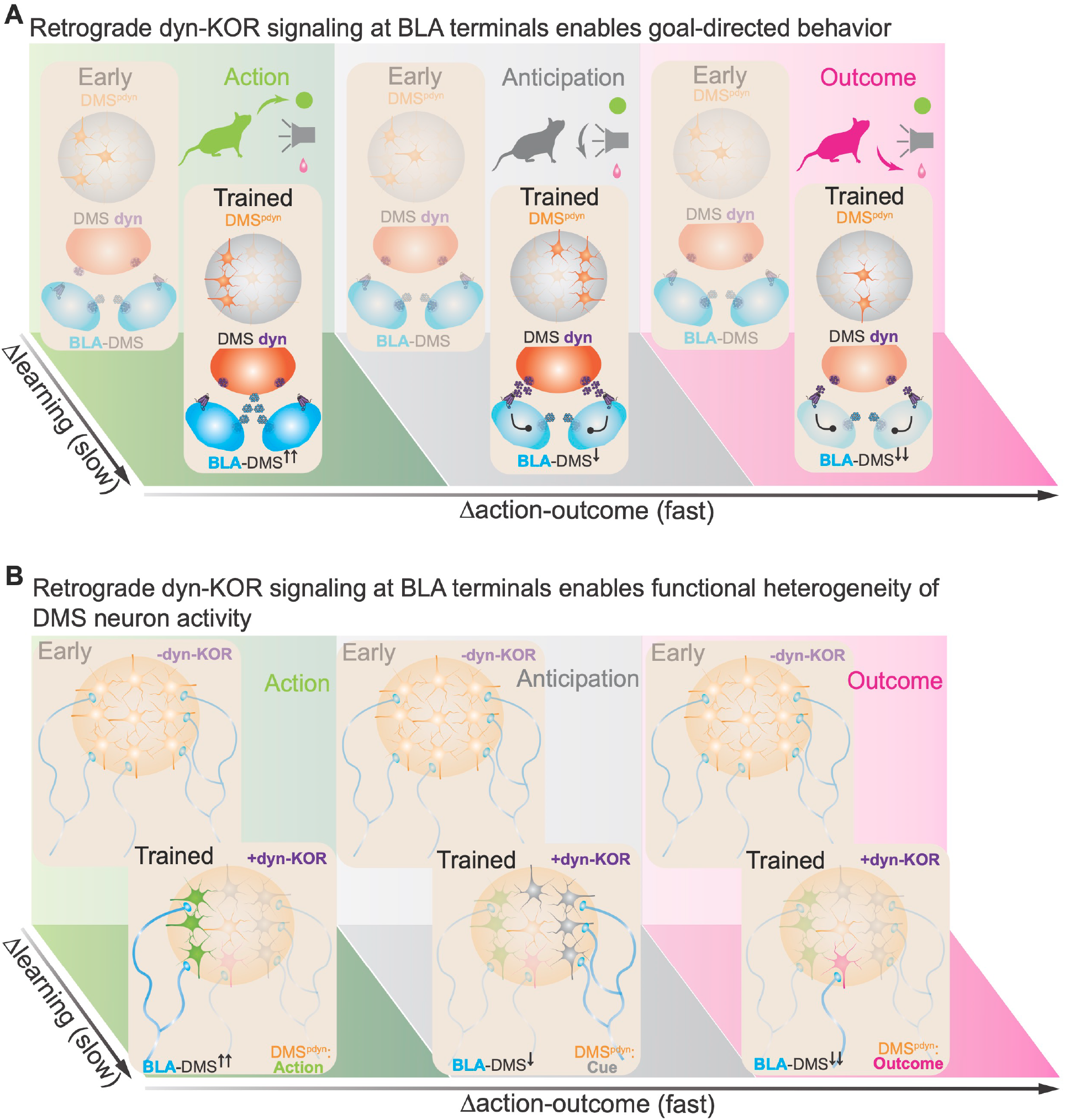
Endogenous opioid dynamics in the dorsal striatum sculpt neural activity to control goal-directed action. (**A**) Cartoon summarizing results showing that retrograde dyn-KOR signaling at BLA-DMS terminals sculpts goal-directed behavior. (**B**) Cartoon summarizing hypothesis showing that retrograde dyn-KOR signaling across learning enables functional heterogeneity of DMS neuron activity.

Dorsal striatal D1 SPNs are known to be involved in movement initiation (Kravitz et al. 2010, Kønig et al. 2019), motor learning (Jin et al. 2014) and action-outcome control (Tai et al. 2012, Matamales et al. 2020, Peak et al. 2020, Bloem et al. 2022). Yet, these prior studies have lacked the spatiotemporal precision to study how D1 SPN encoding of goal-directed behavior evolves, or to assess the distinct contributions they make to specific components of the behavior, such as actions, cued anticipation or outcomes. Here, we used a very recently published technology we developed using microprisms implanted in the DMS (Hjort et al. 2024), allowing us to image and track >14,000 neurons over days and weeks in a single behavioral task (**Fig. 1**). Using this approach, we demonstrated that DMS neuron activity evolves functional heterogeneity; while DMS neurons appear agnostic to behavioral variables early in learning, they segregate into clusters defined by action, cued anticipation and outcome as they learn behavior. While studies prior have reported behavior-selective activity in striatal SPNs (Bloem et al. 2022), the evolution of such patterns during goal-directed learning has only been observed in cortical areas (Namboodiri et al. 2019, Reinert et al. 2021, Ottenheimer et al. 2023). Furthermore, using D1R-tdTomato mice and co-registering td-Tomato positive cells in the field of view after every session, we were able to identify 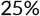 of all DMS neurons as D1/dyn neurons. This percentage is lower than that reported using *in situ* hybridization techniques (Gerfen et al. 1990), likely attributable to a combination of the efficiency of viral transduction (GCaMP6s), genetically-encoded fluorophore expression and limited detection using two-photon imaging with td-Tomato’s efficiency. Indeed, our ISH revealed a very similar % of DRD1 mRNA in the DMS and a high degree of overlap between pdyn and DRD1 mRNA. Importantly, we showed that D1/dyn neurons preferentially contribute to action and cued anticipation, owing to their enrichment in these behavioral clusters (**Fig. 1K,L**). Previous studies have shown that D1 SPNs encode action-outcome learning (Peak et al. 2020) and a high degree of cue-responsive neurons in the striosomal compartments of the dorsal striatum (Bloem et al. 2017). Thus, our studies corroborate previous findings and helped to inform and guide the future experiments that followed.

Despite having known for decades that a large percentage of D1 SPNs express dyn in the DMS (Reiner and Anderson 1990), dyn has largely been used as a marker for these cells, and has not been given a definitive functional role within these canonical neural circuits. Furthermore, only recently have we been able to detect dynorphin levels (Al-Hasani et al. 2018), allowing for the possibility of definitive attributions to the role of dyn-KOR signaling to behavior. To address the role of endogenous opioid signaling in naturalistic, goal-directed behavior, we recently developed a suite of novel, genetically-encoded opioid biosensors, including a dyn biosensor (kLight1.3a) (Rappleye et al. 2022, Tian et al. 2023, Zhou et al. 2023). Using kLight1.3a, we found that dyn is released in the DMS upon cued anticipation of reward, at strikingly fast timescales, suggesting a hitherto-unknown contribution of dyn release to action-outcome sequences (**Fig. 2**). Importantly, we do not see an appreciable increase in dyn to cues during Pavlovian conditioning, suggesting that learning action-outcome behavior is necessary for dyn release to evolve. Of note, we also observe a reduction kLight fluorescence below baseline following reward delivery that increases in magnitude with Pavlovian conditioning, but remains relatively stable thereafter (**Fig. S2H,I, 2E-G**). We posit that this reduction, perhaps biological, may be artifactual in nature owing to our observations that aticaprant blocked elevations in dyn during cued anticipation, yet the reductions in fluorescence remained intact. Moreover, our prior studies show that stimulated release in animals lacking dyn (Sharifi et al. 2001) precludes kLight fluorescence. The increase in dyn during cued anticipation, along with the functional specialization of DMS dyn neurons after goal-directed learning, suggest that dyn neuron activity and subsequent dyn release are recruited to shape goal-directed behavior.

We subsequently explored this role for dyn in goal-directed behavior, during learning across slower timescales and in sustained action-outcome behavior across faster timescales (**Fig. 3**). Stimulating dyn neuron activity via optogenetics, for the period we observe kLight fluorescence resulted in a significant increase in goal-directed behavior. Furthermore, ongoing dyn neuron stimulation during the cue prevented animals from extinguishing their operant responding. Additionally, KOR antagonism resulted in a reduction of behavior to control levels of operant responding. Interestingly, while animals on aticaprant elevated their nosepokes to the same degree, they showed deficiencies in reward consumption (**Fig. 3E, S3E**), suggesting that while dyn-KOR signaling following an action is necessary for promoting action-outcome sequences. Eliminating pdyn from the DMS resulted in a significant blunting of goal-directed learning and inflexibility in adapting to changes in operant contingencies such as reversal learning and fixed-ratio 3 responding, suggesting that an ongoing, perhaps growing tone of dyn at slower timescales is essential for goal-directed behavior. Remarkably, this is further evidenced by the rescue of these deficits in behavior by an acute i.p injection of a KOR agonist (**Fig. 3L, S3M**). These studies drawilar parallels to that observed with drug self-administration, wherein theories have suggested that this is due to a growing dyn tone, owing to drug craving (Wee and Koob, 2010). Indeed, previous studies have showed slow and sustained elevations in pdyn mRNA levels following drug self-administration behavior (Hurd et al. 1992, Hurd and Herkenham 1993, Fagergren et al. 2003), and consequentially, activating (Redila and Chavkin 2008, Ehrich et al. 2014), or antagonizing (Wee et al. 2009, Walker et al. 2011) KOR impacts drug-seeking and reinstatement.

It is important to note that recent studies have provided results that present alternative findings to those in this manuscript. One study showed that exposure to norBNI systemically, a long-lasting KOR antagonist, results in enhanced operant learning (Farahbakhsh et al. 2023). Another study reported that animals lacking pdyn in all D1R-expressing neurons from birth resulted in increased flexibility of operant behavior, possibly due to long-term changes in plasticity at D1-SPNs via dyn-KOR signaling in the dorsal striatum (Yang et al. 2023). In our work, dyn expression was manipulated only in the DMS during adulthood. Several brain regions express pdyn and KOR, and multiple other brain regions in addition to the dorsal striatum show dyn/D1R overlap (Drago et al. 1994, Perreault et al. 2010, Kim et al. 2017, Wang et al. 2024). Furthermore, whereas infusion of KOR agonists into multiple brain regions produced aversion, infusion into the dorsal striatum did not (Bals-Kubik et al. 1993), suggesting a putative different role for dyn-KOR control in the dorsal striatum. Ultimately, these reported findings do not conflict with the results in this study and serve to further implicate endogenous opioid signaling in the modulation of adaptive behavior, which only a few studies have done (Abraham et al. 2021, Wang et al. 2024). Indeed, future studies warrant a targeted understanding of the cell-types, circuits and time window of action of dyn-KOR signaling.

We next asked what the locus of action of dyn-KOR signaling was and relied on prior evidence that opioids may function to dampen neuronal circuit activity in a retrograde manner (Ehrich et al. 2014, Tejeda et al. 2017, Castro et al. 2021), although most of these studies were predominantly in *ex-vivo* preparations (**Fig. 4**). Deleting KOR from inputs into the DMS also resulted in deficits in goal-directed behavior. Among the regions that project to the DMS, the BLA has been implicated as a hub for motivated behaviors (Wassum and Izquierdo 2015, Namburi et al. 2016). The existence of amygdalar projections across the striatum has been known for decades (Kelley et al. 1982), but only recently have studies begun to assess their functional role (Corbit et al. 2013, Courtin et al. 2022, Giovanniello et al. 2023). Furthermore, the BLA is enriched in KOR expression (Crowley et al. 2016, Nygard et al. 2016). We found a significant population of KOR^+^ DMS-projecting BLA neurons that preferentially project to D1/dyn neurons in the DMS and are under dyn-KOR control. KOR deletion from the BLA resulted in a profound reduction in goal-directed learning and extinction. Despite evidence for the BLA in such behaviors, studies prior have only looked at the contribution of KOR in the BLA for aversive behaviors (Knoll et al. 2011) or for the reinstatement of drug preference (Nygard et al. 2016). We then asked if and when BLA-DMS projections are active during goal-directed behavior (**Fig. 5**). Again, we found BLA-DMS engagement during behavior across multiple timescales. General engagement during behavior grew as animals learned behavior, while the terminals displayed a bimodal pattern of activity across action-outcome sequences (**Fig. 5E-G**). These fluctuations in activity were also specific to behavior, as they markedly decreased in magnitude upon extinction. Whereas increases in GCaMP6s from axon terminals has been widely reported, studies observing reductions in fluorescence have been less prominent. Such reductions perfectly capture possible dampening of neuronal activity by Gai GPCR-coupled mechanisms, either via the reduction of cAMP production and subsequent decrease in intracellular calcium (slow), or via the inhibition of calcium channels (fast). Indeed, studies are already underway utilizing cAMP sensors to ascertain the impact of GPCR signaling on neuronal activity and behavior (Zhang et al. 2023). While future studies are required to uncover what component of behavior or behavioral state this reduction is causal to, our subsequent studies that disrupt (with ChR2; activation) or enhance (with PPO; inhibition) this inhibition suggest that the magnitude of this reduction contributes to outcome consumption and subsequent action-outcome behavior. Interestingly, studies have reported similar reductions in BLA terminal activity in the NAc to reward consumption during pavlovian and operant conditioning (Reed et al. 2018), but they did not report any increase in activity during action suggesting that BLA terminals in the DMS are differentially engaged in goal-directed behavior. It is also important to note that studies observing activity in this projection have only been conducted at the level of the cell bodies in the BLA (Courtin et al. 2022, Giovanniello et al. 2023). While both studies report an engagement in activity, they did not observe reductions in activity of bulk fluorescence, or populations of single soma. This may suggest that BLA terminals may be under differential refinement in activity, particularly during outcome consumption, independent of BLA soma activity.

We then determined whether this inhibition of BLA-DMS terminal activity was due to dyn-KOR signaling (**Fig. 6**). Strikingly, loss of dyn from DMS neurons drastically impacted the magnitudes of these signals at the BLA-DMS terminals, while also affecting goal-directed behavior. These results, along with our earlier experiments on the impact of BLA KOR on behavior, strongly suggest that BLA-DMS terminals are under dyn-KOR control, via retrograde release of dyn and KOR modulation of BLA terminal activity. These results may also lend support to the possibility that neuropeptide signaling uniquely sculpts axonal activity, independent of cell body activity, thereby resulting in refinement of behavior. Whereas retrograde release of dyn has been posited as a mechanism of action to modulate dopamine release in the striatum (Ehrich et al. 2014, Tejeda and Bonci 2019, Gordon-Fennell et al. 2023b), this is the first study exploring its impact on circuits important for reward processing beyond dopamine. Of note, while our results thus far strongly suggest that BLA-DMS terminals are modulated by dyn-KOR signaling, the nature of DMS dyn neuron activation and subsequent dyn release is as yet unclear (**Fig. 7**). Hence, we used *in vivo* two-photon calcium imaging multiplexed with spatial light modulation to stimulate spatial subsets of BLA axon terminals in the DMS. We found 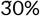 of DMS neurons are active upon BLA axon stimulation and a small but significant enrichment of activated DMS neurons in the cue cluster. These results suggest a preponderance for BLA axons to preferentially activate DMS neurons that encode cues. Next we assayed the ability of BLA axons to induce dyn release. We found that when mice nosepoke for BLA terminal stimulation, there is a significant increase in dyn release *in vivo*. Finally, to determine the sufficiency for this dyn release during cued anticipation to promote action-outcome behavior, we stimulated dyn release during behavior. Indeed, we observed a marked increase in the magnitudes of both the peak and the minimum of BLA-DMS activity during action and outcome, respectively.

Our results raise important questions as to the nature of neuropeptide GPCRs and their role in sculpting behavior, across multiple timescales. It is well-established that neuropeptide-GPCR signaling can exert their effects on pre- and post-synaptic activity by inhibiting ion channels (fast) and downstream signaling mechanisms (slow). Our study uniquely demonstrates the synergy between these mechanisms resulting in the stabilization of neural activity, affording the refinement of goal-directed behavior. Furthermore, we propose that dyn-KOR signaling at BLA terminals enables the stability and segregation of DMS neuron activity during distinct behavioral epochs (**Fig. 8B**). Herein, (i) BLA terminal excitation during action promotes the activity of cue-selective DMS^pdyn^ neurons and subsequent release of DMS^dyn^ during cued anticipation, (ii) thereby inhibiting BLA terminals via retrograde dyn-KOR inhibition, and (iii) affording the stability of DMS^pdyn^ neuron selectivity to encode distinct variables during goal-directed behavior.

More specifically, our study raises key questions to follow up - What is the functional role for dyn-KOR signaling in goal-directed behavior? Why is dyn released during cued anticipation and what component of an action-outcome behavior does it contribute to? As we observed an increase in dyn across learning (**Fig. 2**), our results suggest that dyn could be signaling the salience of the context animals find themselves in to obtain rewards. Alternatively, dyn may contribute to determining outcome value in the context of action-outcome learning. At the circuit level, it is particularly interesting to delve deeper into why the retrograde inhibition of neural activity by dyn-KOR signaling is critical for acquiring robust learning and sustaining goal-directed behavior (**Fig. 6**). We propose a model whereby dyn-KOR mediated inhibition of BLA-DMS activity during outcome may (i) proffer (switch the animal) to the experience of an outcome and (ii) thereby negatively regulate the action to ensure the completion of the action-outcome sequence. This is particularly apparent in our experiments wherein optogenetic inhibition during extinction prevented the animals from extinguishing their operant responding. The inhibition of information from a circuit normally thought to regulate appetitive behavior upon achieving the outcome lends credence to this possibility; however, future studies are required to tease this apart.

In summary, we ascribe a unique dynamic role for the neuropeptide GPCR signaling via dynorphin in the acquisition of learning and in shaping refinement of goal-directed behavior across multiple timescales (**Figure 8**). Our studies describe a novel, retrograde presynaptic GPCR inhibition mechanism whereby dyn-KOR control of BLA projections to the DMS promote behavior. These efforts illuminate a window for dyn-KOR action during the acquisition and maintenance of these behaviors and provide necessary insight to guide the much anticipated therapeutic development for the treatment of neuropsychiatric disorders.

## 3 METHODS

### Animals

Adult (18–35 g) male and female Ai14 x DrD1-Cre, *Oprk1*-Cre (KOR-cre), wildtype (WT), Pdyn-IRES-Cre (*Pdyn*-Cre), *Pdyn*^*fl/f*^ (Pdyn cKO) and *Oprk1*^*fl/f*^ (KOR cKO) mice were group housed, given access to food pellets and water ad libitum, and maintained on a 12 hr:12 hr light:dark cycle (lights off at 9:00 AM, lights on at 9:00 PM). All mice were kept in a sound-attenuated, isolated holding facility one week prior to surgery, post-surgery, and throughout the duration of the behavioral assays to minimize stress. For cell-type conditional deletion and optogenetic experiments we used age-matched Cre- cage, littermate and WT controls. Unless otherwise noted, animals had *ad libitum* access to food and water. Any variation from these approaches was due to behavioral attrition from off-target injections/implants or headcap failures. All animals were drug and test naive, individually assigned to specific experiments as described, and not involved with other experimental procedures. Statistical comparisons did not detect any significant differences between male and female mice, and were therefore combined to complete final group sizes. All animals were monitored for health status daily and before experimentation for the entirety of the study. All procedures were approved by the Animal Care and Use Committee of the University of Washington, and conformed to US National Institutes of Health guidelines.

### Stereotaxic Surgery

All coordinates, viruses and implant type for experiments are listed in Table S1. After mice were acclimated to the holding facility for at least seven days, the mice were anaesthetized in an induction chamber (1%-4% isoflurane) and placed into a stereotaxic frame (Kopf Instruments, model 1900) where they were mainlined at 1%-2% isoflurane. For mice receiving viral injections followed by microprism implants, we used a Nanoject II (Drummond Scientific) to inject 4 x 300 nL of virus at a rate of 100 nL/min. For all other viral injections, a blunt needle (86200, Hamilton Company) syringe was used to deliver 400 nL of virus at a rate of 100 nL/min either in the DMS or BLA. For mice receiving intracranial implants (i.e., microprism implants, fiber photometry or optogenetic optic fibers), a hole was drilled above the site of interest and the implant was slowly lowered to the coordinates. Cannulas were secured to the skull using one bone screw and super glue (Lang Dental). All other implants were secured using MetaBond (C & B Metabond). For mice undergoing head-fixed experiments, a head-ring was placed over the implant before securing them and the implants with MetaBond. For further detail on 1.5 x 1.5 x 8mm microprism (OptoSigma) implantation, please refer to (Hjort et al. 2024).

### Head-fixed Operant Behavior

Experiments were performed as previously described (Gordon-Fennell et al. 2023a). In brief, animals were signal checked for GcaMP6s dynamics 4-6 weeks after surgery by securing them in the OHRBETS platform. Animals were water-restricted for 1 week and maintained at 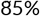 body weight. Animals then underwent sipper training for 10% sucrose for 3 days, followed by Pavlovian conditioning for 3 days to associate a 3s tone followed by 3s access to sucrose via sipper extension, then operant conditioning to rotate a wheel for 8 days. Rotating the wheel a half-turn in the “active” contingency yielded the tone and sucrose availability, the “inactive” contingency yielded nothing. Tone delivery also resulted in a brake applied to the wheel. This was followed by operant extinction where active rotations resulted in the tone and sipper extension, but no sucrose for 2 days.

### Operant Index for Head-fixed Behavior

To accurately capture all components of an action-outcome behavior, we devised a summary metric called the operant index taking into account action discrimination (Active – Inactive wheel rotations), action vigor (total wheel rotations) and outcomes consumed (every trial the animal consumed sucrose). This is represented as

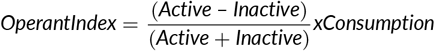

We then normalized this value to a hypothetical index under ideal conditions involving the active rotation contingencies required to achieve the maximum outcomes in Pavlovian conditioning (40 trials), half the number of inactive rotation contingencies to achieve discrimination and the maximum outcomes consumed in Pavlovian conditioning (40 trials). For extinction, we used the number of trials the animals performed a lick at the sipper instead of outcomes consumed. For transparency, active-inactive wheel rotation data is reported in the supplemental figures.

### Two Photon Imaging

For details on how imaging and longitudinal tracking was performed, please refer to (Hjort et al. 2024). Prior to each behavior session, animals were placed in the OHRBETS platform outfitted with a Thorlabs goniometer (TTR001/M, Thorlabs) to facilitate levelling of the imaging plane. Imaging was conducted on a Bruker 2p+ (Bruker) at 920 nm using the Cousa objective (20mm working distance; Pacifica Optics). Following identification of the same field of view (FOV) to enable longitudinal tracking, animals underwent 15 minute sessions of lick training, Pavlovian conditioning, Operant conditioning or Operant extinction where imaging was acquired at 7.5 Hz using resonant galvos (4 frame averaging). At the end of each session, a 5 minute static recording was acquired at at 7.5 Hz using resonant galvos (4 frame averaging) at 1080 nm wavelength in the same FOV to register td-Tomato postitive cells. Imaged cells were tracked longitudinally by concatenating recording sessions and running them through Suite2p (HHMI, Janelia Research Campus). Sorted cells were manually verified as well. GcaMP6s and Td-Tomato cells were overlaid in ImageJ/Fiji and quantified for overlap. Spectral clustering, peak analysis, RNN and GLM analyses were performed using custom Python code based on (Namboodiri et al. 2019). All data was was z-scored using the mean fluorescence and standard deviation of a predefined time window (5s prior active wheel rotation). Data are presented as z-score across −10s to 7s, with 0s signifying the start of tone delivery.

For targeted sequential photostimulation experiments of BLA axons in conjunction with DMS two-photon imaging, a second laser path with a 1040nm high powered femtosecond laser (Spirit One, SpectraPhysics) was used with a pair of galvanometers to generate spiral montages across the FOV, similar to (Yang et al. 2018, Piantadosi et al. 2024). Spirals of 5-10 um in diameter were generated at 20Hz, 5ms pulse-widths for 500 ms, every 10 seconds at 10 mW of laser power, sequentially tiling the entire FOV. In parallel, animals also underwent “baseline” sessions resulting in shutter opening for the same frequency, but with the laser off. Data was normalized to a 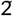 minute baseline prior to stimulation. “Active” cells were determined based on whether their activity was higher than the peak activity in the “baseline” sessions. These sessions were then concatenated with the recordings from the trained operant session and analyzed in Suite2p for tracking to determine the identities of the neurons enriched in behavioral clusters. Point spread functions to assess the utility for microprims for spatial stimulation were assessed in (Hjort *et al*., 2024). All data was was z-scored using the mean fluorescence and standard deviation of a predefined time window (1s prior to laser stimulation). Data are presented as z-score across −5s to 5s, with 0s signifying the start of laser stimulation.

### RNN Decoding

To determine whether cell identities could be decoded from trace responses, we used a Recurrent Neural Network (RNN) Long Short-Term Memory Network (LSTM). This allowed us to additionally capture temporal dependencies in the decoding analysis. Cells were labeled based on the period (baseline, action, cue, reward) of their maximal activity. We used data from Day 8 to train the model, holding out 20% of the cells for evaluation. The accuracy for Day 8 was computed as the proportion of the held-out cells for which the prediction matched the label. We then utilized the same model weights to predict the identities of cells across each session, allowing us to directly compare the model’s differential decoding ability across sessions. In addition, we trained the model with baseline data from each cell, as well as separately with a single random shuffle of cells and labels to determine whether the model’s decoding ability was significantly stronger than expected by chance.

### GLM Encoding

To determine changes in fluorescence traces caused by the behavioral variables, we used a Gaussian generalized linear model (GLM). First, we pre-processed the raw fluorescence traces, by applying a Butterworth lowpass filter, with a cutoff set at 20% of the Nyquist frequency, followed by z-scoring across the trace. Since we are primarily interested in determining changes in activity responses to behavioral variables, we defined explanatory variables as spanning multiple frames with respect to the fluorescence trace, as opposed to a time varying kernel approach (Namboodiri et al. 2019). This implies that the coefficient for each term represents the change in activity attributable to the explanatory variable represented. We defined the opening of the solenoid, the presence of the cue, an active wheel rotation, and an interaction term between the action period (ref to previous figure/mention) and an active wheel rotation as explanatory variables. This was because we are interested in understanding the relationship between wheel rotations during the action period, as opposed to during the entire trial, which would be the case if we simply used the coefficient for the active wheel rotation term. The final GLM equation was as follows:

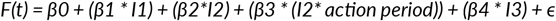

*F(t)*represents pre-processed fluorescence on frame *t. β* represents the coefficients and *I* represent indicator variables corresponding to each behavioral input. More specifically, I1 corresponds to the solenoid, I2 corresponds to active wheel rotations, and I3 corresponds to the presence of the cue. The action period corresponds to the 5 second window prior to the instance of the cue. To minimize any possible overlap between the action and cue periods, we elected to use the first 3 seconds of the action window. Each of these variables was coded as 0 for frames in which they did not occur, and as 1 on frames where they did occur. *B*0 is the intercept term and *ϵ* is the error term. To determine cells that may be encoding information related to specific explanatory variables, we first matched the cluster identity of each cell with an explanatory variable (cluster 2 (reward) = solenoid, cluster 1 (action) = active wheel rotation/ action period interaction term, cluster 0 (cue) = tone). As we were interested in understanding encoding patterns across the training paradigm in cells that encoded this information on the trained day (day 8), we then subset the dataset to include cells with GLM coefficients which differed significantly (p <0.05) from 0 on the trained day. To account for inhibitory modulation, we took the absolute value of the GLM coefficients from subset cells, after which we considered changes in the coefficients for these cells across different sessions. This allows us to quantify the extent to which the cells that encode information on the trained day encode that same information at different points in the training paradigm.

### Freely Moving Operant Behavior

For freely moving experiments, animals were food-restricted 6-8 weeks following surgery to maintain their body weight at 85%. If undergoing self-stimulation experiments, animals were tethered and placed in an operant chamber (MedPC). Animals nosepoked into illuminated nose-poke ports, randomly designated “active” or “inactive”, with the active poke resulting in a 1s or 7s delivery of 465 nm (1-5mW laser power) or 635 nm (1-2mW laser power) laser light at 20Hz, 5 ms pulse-width. For experiments involving sucrose, animals were tethered (if tethering was required), placed in an operant chamber (MedPC) and underwent magazine training for random delivery of sucrose pellets (BioServ) in a receptacle for 3 days. Following this, animals underwent Pavlovian conditioning to associate a 5s houselight succeeded by sucrose pellet delivery for 5 days. Mice then underwent operant conditioning, where they nosepoked into illuminated nosepoke ports, randomly designated “active” or “inactive”, with the active poke resulting in a 2s timeout, then 5s houselight, followed by sucrose delivery for 5 days. When specified, animals then underwent operant reversal, where the nosepoke ports were reversed, fixed ratio-3, where they performed 3 nosepokes instead of 1 for cue and reward delivery of the same time, and progressive ratio testing where the nosepokes required escalated exponentially as we previously described (Parker et al. 2019). If undergoing PR, animals were returned to fixed ratio-1 for 3 days until stable responding was achieved, and then underwent operant extinction for 5 days. Here, an active nosepoke yielded cue delivery, but no sucrose delivery.

### Operant Index for Freely Moving Behavior

Similar to head-fixed behavior, we devised an operant index taking into account action discrimination (Active – Inactive nosepokes), action vigor (total nosepokes) and sucrose pellets consumed. This was normalized to a hypothetical index under ideal conditions involving the active nosepokes required to achieve the maximum outcomes in Pavlovian conditioning (40 trials), half the number of inactive nosepokes to achieve discrimination and the maximum outcomes consumed in Pavlovian conditioning (40 trials). For extinction, we used the number of approaches the animal made to the sucrose hopper instead of outcomes consumed. For transparency, all active and inactive nosepokes, rewards consumed and approaches to hopper are quantified in the supplemental figures.

### *In Vivo* Fiber Photometry

Fiber photometry recordings were made throughout the entirety of 60-minute Operant testing. Prior to recording, an optic fiber was attached to the implanted fiber using a ferrule sleeve (Doric, ZR_2.5). Two LEDs were used to excite GCaMP6s. A 531-Hz sinusoidal LED light (Thorlabs, LED light: M470F3; LED driver: DC4104) was bandpass filtered (470 ± 20 nm, Doric, FMC4) to excite GCaMP6s and evoke Ca^2+^-dependent emission. A 211-Hz sinusoidal LED light (Thorlabs, LED light: M405FP1; LED driver: DC4104) was bandpass filtered (405 ± 10 nm, Doric, FMC4) to excite GCaMP6s and evoke Ca^2+^-independent isosbestic control emission. Prior to recording, a 120 s period of GCaMP6s excitation with 405 nm and 470 nm light was used to remove the majority of baseline drift. Laser intensity for the 470 nm and 405 nm wavelength bands were measured at the tip of the optic fiber and adjusted to 30 μW before each day of recording. GCaMP6s fluorescence traveled through the same optic fiber before being bandpass filtered (525 ± 25 nm, Doric, FMC4), transduced by a femtowatt silicon photoreceiver (Newport, 2151) and recorded by a real-time processor (TDT, RZ10). The envelopes of the 531-Hz and 211-Hz signals were extracted in real-time by the TDT program Synapse at a sampling rate of 1017.25 Hz. For the ChrimsonR stimulation experiments, a 635 nm laser was used with a custom filter cube (Doric) at 1-2 mW intensity to deliver red light through the tip of the same optic fiber used to excite GCaMP6s similar to (Al-Hasani et al. 2021). For operant behavior, data are presented as z-score across −30s to 30s, with 0s signifying a nosepoke. For Pavlovian, data are presented as z-score across −30s to 30s, with 0s signifying the start of cue delivery.

### Photometry Analysis

Custom MATLAB scripts were developed for analyzing fiber photometry data in context of mouse behavior and can be accessed via GitHub. The isosbestic 405 nm excitation control signal was scaled to the 470nm excitation signal, then this refitted 405nm signal was subtracted from the 470 nm excitation signal to remove movement artifacts from intracellular Ca^2+^-dependent GCaMP6s fluorescence. Baseline drift was evident in the signal due to slow photobleaching artifacts, particularly during the first several minutes of each hour-long recording session. A double exponential curve was fit to the raw trace and subtracted to correct for baseline drift. After baseline correction, the photometry trace was z-scored relative to the mean and standard deviation of the entire session. The post-processed fiber photometry signal was analyzed in the context of animal behavior.

### Patch-Clamp Electrophysiology

Mice were anesthetized with pentobarbital (50 mg/kg) before transcardial perfusion with ice-cold sucrose cutting solution containing the following (in mM): 75 sucrose, 87 NaCl, 1.25 NaH_2_P0_4_, 7 MgCl_2_, 0.5 CaCl_2_, 25 NaHCO_3_, 306-308 mOsm. Brains were then rapidly removed, and coronal sections 300 μm thick were taken using a vibratome (Leica, VT 1200). Sections were then incubated in aCSF (32°C) containing the following (in mM): 126 NaCl, 2.5 KCl, 1.2 NaH_2_P0_4_, 1.2 MgCl_2_, 2.4 CaCl_2_, 26 NaHCO_3_, 15 glucose, 305 mOsm.. After an hour of recovery, slices were constantly perfused with aCSF (32°C) and visualized using differential interference contrast through a 40x water-immersion objective mounted on an upright microscope (Olympus BX51WI). Whole-cell patch-clamp recordings were obtained using borosilicate pipettes (4– 5.5 MΩ) back-filled with internal solution containing the following (in mM): 117 Cs-Methanesulfonate, 20 HEPES, 0.4 EGTA, 2.8 NaCl, 5 TEA, 5 ATP, and 0.5 GTP (pH 7.35, 285 mOsm). To assess connectivity between BLA and DMS, voltage clamp recordings were performed from cells located near eYFP-expressing axons within the NAC. 5 ms blue light pulses were delivered through the objective while holding each cell at −70 mV to assess glutamatergic input, respectively. U69 or vehicle washes were conducted after collecting stable evoked EPSCs for 20 minutes. Data acquisition occurred at 10 kHz sampling rate through a MultiClamp 700B amplifier connected to a Digidata 1440A digitizer (Molecular Devices). Data were processed using Clampfit v11.0.3.03 (Molecular Devices) and analyzed using GraphPad Prism v8.3.0. All tests were two-sided and corrected for multiple comparisons or unequal variance where appropriate.

### Tissue Processing

Unless otherwise stated, animals were transcardially perfused with 0.1 M phosphate-buffered saline (PBS) and then 40 mL 4% paraformaldehyde (PFA). Brains were dissected and post-fixed in 4% PFA overnight and then transferred to 30% sucrose solution for cryoprotection. Brains were sectioned at 30 mM on a microtome and stored in a 0.01M phosphate buffer at 4ºC prior to immunohistochemistry and tracing experiments. For behavioral cohorts, viral expression and optical fiber placements were confirmed before inclusion in the presented datasets.

### RNAscope Fluorescent *In Situ* Hybridization

Following rapid decapitation of WT, *Pdyn*^fl/fl^ or *Oprk1*^fl/fl^ mice brains were rapidly frozen in 100mL −50°C isopentane and stored at −80°C. Coronal sections corresponding to the site of interest or injection plane used in the behavioral experiments were cut at 20uM at −20°C and thaw-mounted onto SuperFrost Plus slides (Fisher). Slides were stored at −80°C until further processing. Fluorescent in situ hybridization was performed according to the RNAscope 2.0 Fluorescent Multiple Kit User Manual for Fresh Frozen Tissue (Advanced Cell Diagnostics, Inc.). Briefly, sections were fixed in 4% PFA, dehydrated, and treated with pretreatment 4 protease solution. Sections were then incubated for target probes for mouse calcium calmodulin kinase II (CamKIIa, accession number NM_009792.3), vesicular glutamate transporter 1 (*slc17a7*, accession number NM_182993.2), dopamine D1 receptor (*DrD1*, accession number NM_010076.3) kappa opioid receptor (*Oprk1*, accession number NM_001204371.1) and prodynorphin (*Pdyn*, accession number NM_018863.3). All target probes were obtained from Advanced Cell Diagnostics. Following probe hybridization, sections underwent a series of probe signal amplification steps followed by incubation of fluorescently labeled robes designed to target the specific channel associated with the probes. Slides were counterstained with DAPI, and coverslips were mounted with Vectashield Hard Set mounting medium (Vector Laboratories. Images were obtained on an Olympus Fluoview 3000 confocal microscope and analyzed with HALO software. To analyze the images, each image was opened in the HALO software. DAPI positive cells were then registered and used as markers for individual cells. A positive cell consisted of an area within the radius of a DAPI nuclear staining that measured at least 3 positive pixels for receptor probes, or 10 total positive pixels for neurotransmitter probes. Two - three separate slices from the DMS or BLA were used for each animal and that total is presented in the data.

### Statistical Analyses

All data collected were averaged and expressed as mean ± SEM. Statistical significance was taken as ^⍰^p < 0.05, ^⍰⍰^p < 0.01, and ^⍰⍰⍰^p < 0.001 and ****p < 0.0005 as determined by Pearson’s correlation, Student’s t test, one-way ANOVA or a two-way repeated-measures ANOVA followed by post hoc tests as appropriate. Statistical analyses were performed in GraphPad Prism 8.0 (Graphpad, La Jolla, CA) and MATLAB 9.6 (The MathWorks, Natick, MA).

## Abbreviations

DMS: dorsomedial striatum
dyn: dynorphin
KOR: kappa opioid receptor
BLA: basolateral amygdala

## AUTHOR CONTRIBUTIONS

R.G and M.R.B conceptualized, designed the experiments. R..G. collected and analyzed data. R.G. and M.R.B. wrote the manuscript. M.H helped with experimental design performed the analysis for the *in vivo* two photon calcium imaging. P.S and D.S assisted with analysis for the *in vivo* two photon calcium imaging. Z.C.Z assisted with two photon imaging hardware setup. A.A.G helped with head-fixed operant behavior. C.D and L.T conceptualized and designed biosensor development. A.J.E, S.E.S, J.V.T, K.A, D.S, K.M and V.L performed experiments and collected data. D.J.M performed the electrophysiology experiments and collected, and analyzed the data. M.R.B helped lead the design, analysis, oversight of experiments, discuss results, provide resources, and write the manuscript.

## ACKNOWLEDGMENTS

We thank Taylor Hobbs, Carina Pizzano and Bailey Wells for animal colony maintenance. We thank Azra Suko for lab management and virus preparation. We thank Scott Ng-Evans for operations support and Jazmyne Fosha, and Lusine Eyde for programmatic support. We thank Vijay Namboodiri for original Python code, and Christian Pederson for original MATAB code for analyses. We thank Larry Zweifel, Daniel Castro and Benjamin Land for valuable discussions regarding and comments on the manuscript.

## DATA AVAILABILITY

The data supporting this study’s findings are available upon request from the authors.

## SUPPORTING INFORMATION

Additional supporting information may be found in the online version of the article.

---

**Supplementary Figure 1:**
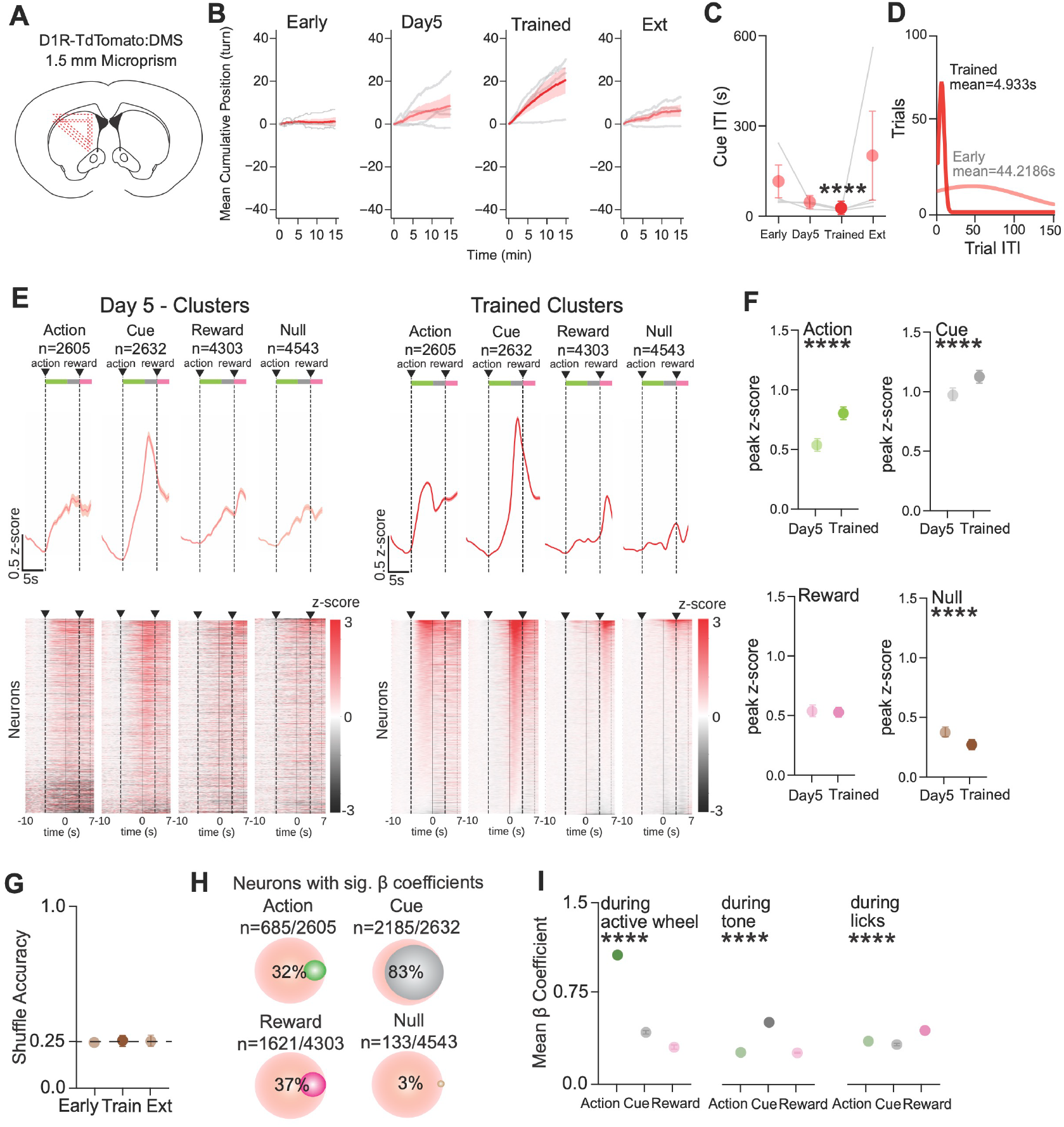
DMS^pdyn^ neurons preferentially encode goal-directed actions and cues. (**A**) Placement maps of prism implantation in the DMS. (**B**) Mean cumulative rotations of active-inactive rotations across early, day5, trained and extinction operant sessions (n=4mice). (**C**) Mean inter-trial interval between cues across early, day5, trained and extinction operant sessions (n=4mice; One Way ANOVA, *p=0*.*0156\**, Kruskal-Wallis statistic=9.154. Multiple comparisons – early vs. trained, *p=0*.*0286\**). (**D**) Trial vs. Trial ITI plot for a representative animal across early and trained operant sessions. (**E**) Spectral clustering classification of day5 operant data (left) and trained operant data (right) across 4 mice showing fluorescence activity traces (top) and heat maps (bottom) of cells from all clusters across time (n=4mice). (**F**) Peak z-score quantification of **Action** clusters (n=4mice; Unpaired t test, *p<0*.*0001\*\*\*\**, t=7.54, df=5208). Peak z-score quantification of **Cue** clusters (n=4mice; Unpaired t test, *p<0*.*0001\*\*\*\**, t=5.63, df=5262). Peak z-score quantification of **Reward** clusters (n=4mice; Unpaired t test, *p>0*.*05*). Peak z-score quantification of **Null** clusters (n=4mice; Unpaired t test, *p<0*.*0001\*\*\*\**, t=6.279, df=9084). (**G**) RNN accuracy for shuffled neural data (n=4mice; One Way ANOVA, p>0.05). (**H**) Summary quantification of % of neurons with significant mean b coefficients (n=4mice). (**I**) Generalized Linear Regression Modeling (GLM) quantification on the trained operant session comparing mean b coefficients of **action, cue and reward** clusters during **active wheel rotations** (n=4mice; One Way ANOVA, *p<0*.*0001\*\*\*\**, F(2,1653)=196217. Multiple comparisons – action vs. cue, *p<0*.*0001\*\*\*\**, action vs. reward, *p<0*.*0001\*\*\*\**), during **tone** (n=4mice; One Way ANOVA, p<0.0001****, F(2,6552)=730332. Multiple comparisons – cue vs. action, *p<0*.*0001\*\*\*\**, cue vs. reward, *p<0*.*0001\*\*\*\**), and during **licks** (n=4mice; One Way ANOVA, *p<0*.*0001\*\*\*\**, F(2,4860)=63753. Multiple comparisons – reward vs. action, *p<0*.*0001\*\*\*\**, reward vs. cue, *p<0*.*0001\*\*\*\**).

**Supplementary Figure 2:**
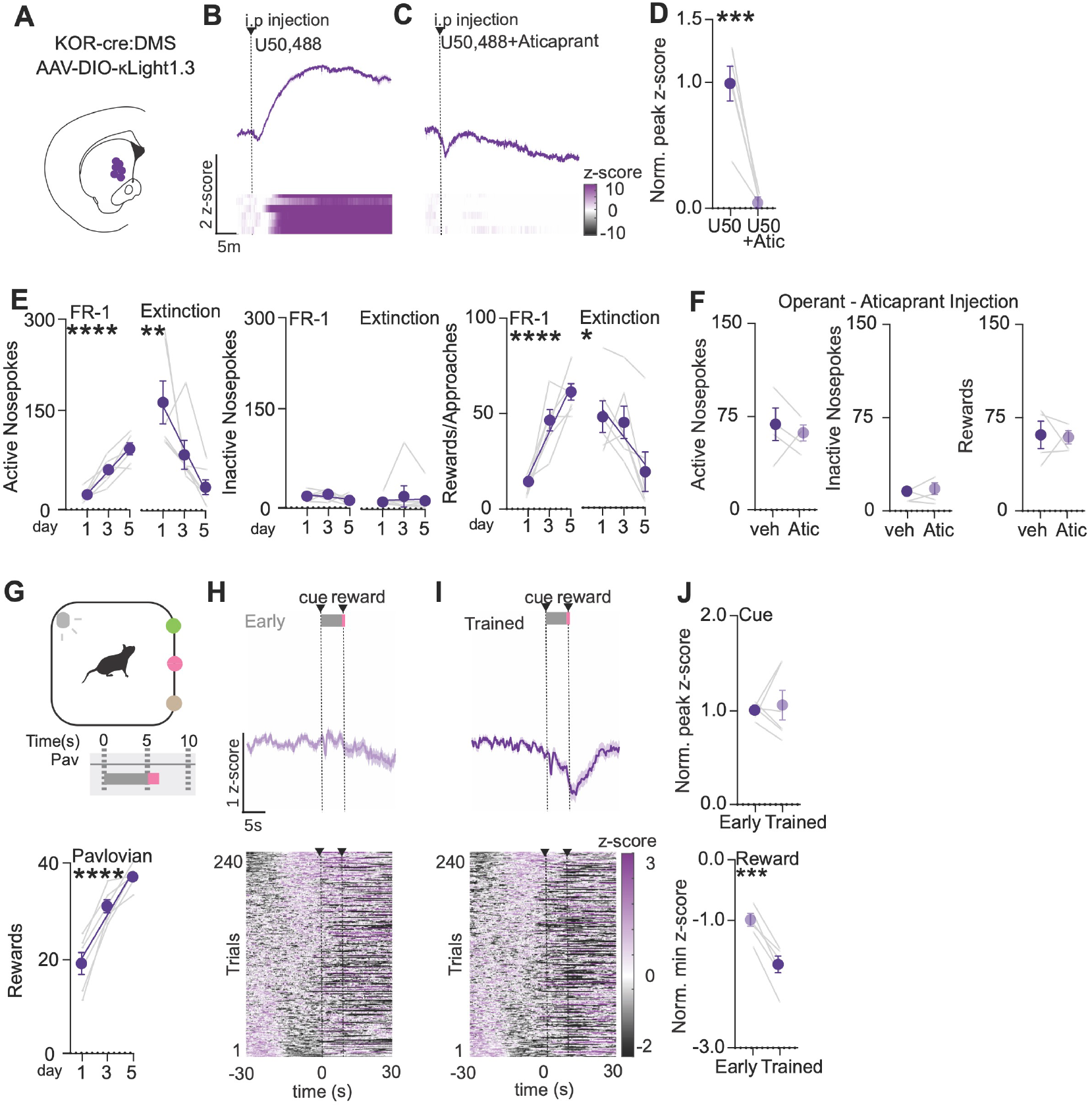
DMS dynorphin is released upon cued anticipation of reward during goal-directed behavior. (**A**) Placement map of optic fiber implantation in the DMS. (**B**) Mean fluorescence and heatmap raster plots of i.p injection of 10 mg/kg U50,488 (n=6mice). (**C**) Mean fluorescence and heatmap raster plots of i.p injection of 10 mg/kg U50,488 and 5 mg/kg Aticaprant (n=6mice). (**D**) Normalized Peak AUC values of U50 vs. U50+Aticaprant (n=6mice; paired t test, *p=0*.*0138\**, t=3.714, df=5). (**E**) Operant behavior during photometry. Left – Active nosepokes during Operant Learning (n=6mice; Simple Linear Regression, R^2^=0.7422, *p<0*.*0001\*\*\*\**, F=46.06) and Operant Extinction (n=6mice; Simple Linear Regression, R^2^=0.4918, *p=0*.*0012\*\**, F=15.48); Middle – Inactive nosepokes during Operant Learning (n=6mice; Simple Linear Regression, R^2^=0.1034, *p>0*.*05*, F=1.846) and Operant Extinction (n=6mice; Simple Linear Regression, R^2^=0.0007, *p>0*.*05*, F=0.01205); Right – Rewards consumed during Operant Learning (n=6mice; Simple Linear Regression, R^2^=0.7748, *p<0*.*0001\*\*\*\**, F=55.04) and Approaches to the reward port during Operant Extinction (n=6mice; Simple Linear Regression, R^2^=0.2364, *p=0*.*0408\**, F=4.953). (**F**) Operant behavior during photometry and aticaprant injection: Left – Active nosepokes during vehicle vs. Aticaprant (n=4mice; paired t test, *p>0*.*05*); Middle - Inactive nosepokes during vehicle vs. Aticaprant (n=4mice; paired t test, *p>0*.*05*); Right – Rewards consumed during vehicle vs. Aticaprant (n=4mice; paired t test, *p>0*.*05*). (**G**) Top - Pavlovian behavior schedule during photometry. Bottom – Rewards consumed during Pavlovian learning (n=6mice; Simple Linear Regression, R^2^=0.7798, *p<0*.*0001\*\*\*\**, F=56.67). (**H**) Mean fluorescence and heatmap raster plots during early pavlovian behavior (n=6mice). (**I**) Mean fluorescence and heatmap raster plots during trained pavlovian behavior (n=6mice). (**J**) Top – Normalized Peak z-score values of early and trained during **cue** period (2-7s) (n=6mice; paired t test, *p>0*.*05*). Bottom – Normalized Minimum z-score values of early and trained during **reward** period (7-30s) (n=6mice; paired t test, *p=0*.*0003\*\*\**, t=8.684, df=5).

**Supplementary Figure 3:**
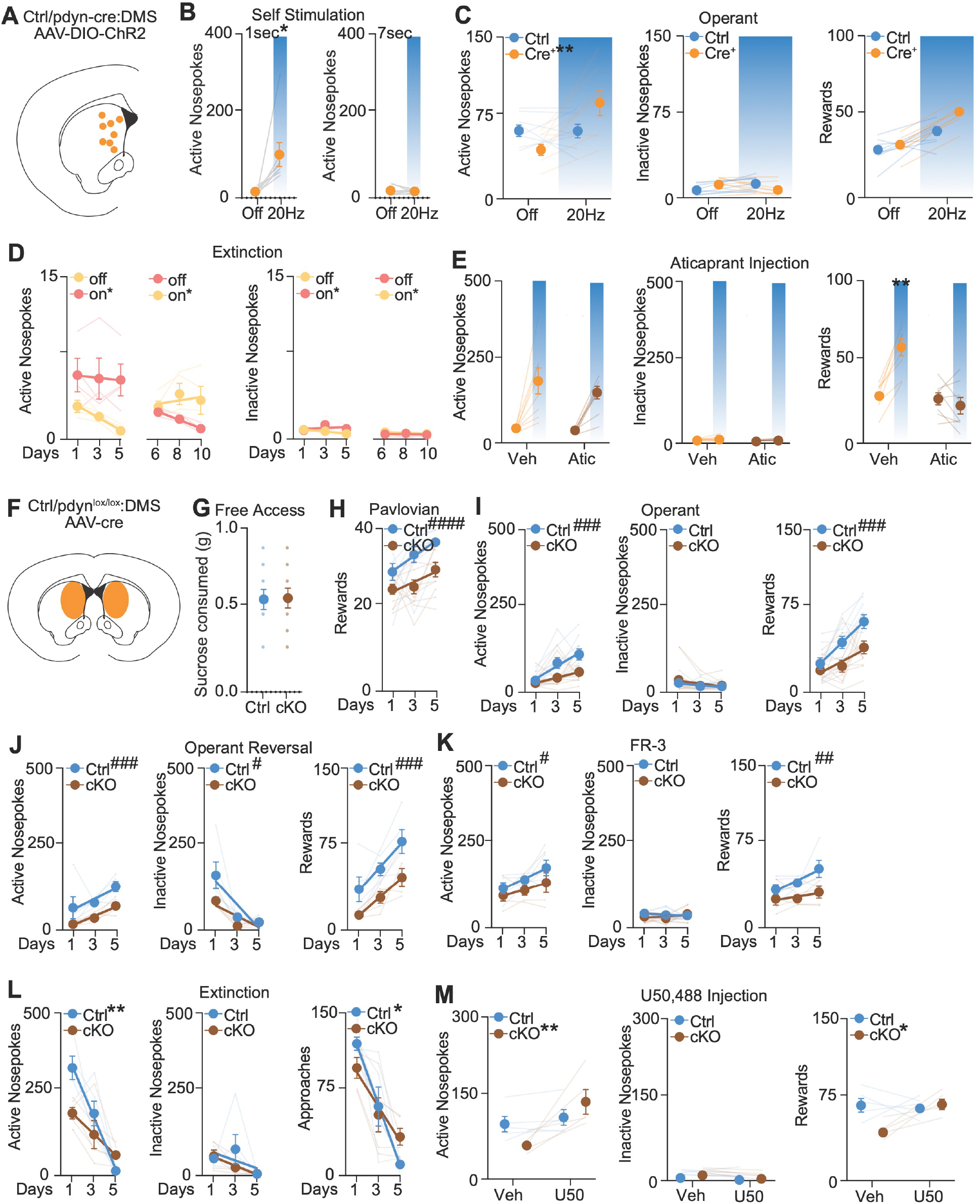
DMS dynorphin is sufficient and necessary for goal-directed behavior. (**A**) Placement maps of optic fiber implants in the DMS. (**B**) Left - Active nosepokes for 1 second self-stimulation (n=8 DMS^pdyn^ mice; paired t test, *p=0*.*0213\**, t=2.9572, df=7). Right - Active nosepokes for 7 second self-stimulation (n=8 DMS^pdyn^ mice; paired t test, *p>0*.*05)*. (**C**) Left – Active Nosepokes during optogenetic stimulation in Ctrl and DMS^pdyn^ mice (n=8 Ctrl,8 dyn-cre mice; Two Way ANOVA, stim x genotype *p=0*.*0036\*\**, F(1,14)=12.00. Multiple comparisons – ctrl off vs. on, *p>0*.*05;* DMS^pdyn^ off vs. on, *p=0*.*0004\*\*\**). Middle – Inactive Nosepokes during optogenetic stimulation in Ctrl and dyn-cre mice (n=8 Ctrl,8 DMS^pdyn^ mice; Two Way ANOVA, stim x genotype *p>0*.*05*). Right – Rewards consumed during optogenetic stimulation in Ctrl and DMS^pdyn^ mice (n=8 Ctrl,8 dyn-cre mice; Two Way ANOVA, stim x genotype *p>0*.*05*). (**D**) Left – Active Nosepokes during extinction days 1-5 (n= 4mice; Off - Simple Linear Regression, R^2^=0.6540, *p=0*.*0014\*\**, F=18.91, On – Simple Linear Regression, R^2^=0.004103, *p>0*.*05*, F=0.04120; Difference in intercepts – *p=0*.*0006*^*##*^), Active Nosepokes during extinction days 1-5 (n=4mice; Off - Simple Linear Regression, R^2^=0.6540, *p=0*.*0014\*\**, F=18.91, On – Simple Linear Regression, R^2^=0.004103, *p>0*.*05*, F=0.04120; Difference in intercepts – *p=0*.*0006*^*##*^). (**E**) Left – Active Nosepokes during optogenetic stimulation in DMS^pdyn^ mice with aticaprant injection (n=8 dyn-cre mice; Two Way ANOVA, stim x treatment *p>0*.*05*). Middle – Inactive Nosepokes during optogenetic stimulation in DMS^pdyn^ mice with aticaprant injection (n=8 DMS^pdyn^ mice; Two Way ANOVA, stim x treatment *p>0*.*05*). Right – Rewards consumed during optogenetic stimulation in DMS^pdyn^ mice with aticaprant injection (n=8 DMS^pdyn^ mice; Two Way ANOVA, stim x treatment *p<0*.*0018\*\**, F(1,7)=23.9. Multiple comparisons – veh off vs. on, *p=0*.*0010**;* aticaprant off vs. on, *p>0*.*05*). (**F**) Schematic of injection site in the DMS. (**G**) Sucrose consumed in homecage (n=9 Ctrl, 12 DMS^pdyn-cKO^ mice; unpaired t test, *p>0*.*05)*. (**H**) Rewards consumed during Pavlovian learning (n=9 Ctrl, 12 DMS^pdyn-cKO^ mice; Ctrl - Simple Linear Regression, R^2^=0.2758, *p<0*.*0049\*\**, F=9.520. DMS^pdyn-cKO^ – Simple Linear Regression, R^2^=0.1292, *p=0*.*0399\**, F=4.6. Difference in Intercepts – *p<0*.*0001*^*##*^). (**I**) Left – Active Nosepokes during operant learning (n=9 Ctrl, 12 DMS^pdyn-cKO^ mice; Ctrl - Simple Linear Regression, R^2^=0.3825, *p<0*.*0006\*\*\**, F=15.49. DMS^pdyn-cKO^ – Simple Linear Regression, R^2^=0.1388, *p=0*.*0252\**, F=5.479. Difference in Intercepts – *p=0*.*0006*^*##*^). Middle – Inactive Nosepokes during operant learning (n=9 Ctrl, 12 DMS^pdyn-cKO^ mice; Ctrl - Simple Linear Regression, R^2^=0.0978, *p>0*.*05*, F=2.691. DMS^pdyn-cKO^ – Simple Linear Regression, R^2^=0.05205, *p>0*.*05*, F=1.867). Right – Rewards consumed during operant learning (n=9 Ctrl, 12 DMS^pdyn-cKO^ mice; Ctrl - Simple Linear Regression, R^2^=0.4804, *p<0*.*00001\*\*\*\**, F=23.11. DMS^pdyn-cKO^ – Simple Linear Regression, R^2^=0.2117, *p=0*.*0048\*\**, F=9.129. Difference in Intercepts – *p=0*.*0002*^*##*^). (**J**) Left – Active Nosepokes during operant reversal learning (n=5 Ctrl, 5 DMS^pdyn-cKO^ mice; Ctrl - Simple Linear Regression, R^2^=0.2660, *p<0*.*0491\**, F=4.710. DMS^pdyn-cKO^ – Simple Linear Regression, R^2^=0.6869, *p=0*.*0001\*\*\*\**, F=28.52. Difference in Intercepts – *p=0*.*0003*^*##*^). Middle – Inactive Nosepokes during operant learning (n=9 Ctrl, 12 DMS^pdyn-cKO^ mice; Ctrl - Simple Linear Regression, R^2^=0.5330, *p=0*.*002\*\**, F=14.84. DMS^pdyn-cKO^ – Simple Linear Regression, R^2^=0.05205, *p=0*.*0028\*\**, F=13.53. Difference in Intercepts – *p=0*.*0465*^*#*^). Right – Rewards consumed during operant learning (n=9 Ctrl, 12 DMS^pdyn-cKO^ mice; Ctrl - Simple Linear Regression, R^2^=0.4578, *p<0*.*0056\*\**, F=10.98. DMS^pdyn-cKO^ – Simple Linear Regression, R^2^=0.5903, *p=0*.*0008\*\*\**, F=18.73. Difference in Intercepts – *p=0*.*0001*^*##*^). (**K**) Left – Active Nosepokes during operant Fixed Ratio 3 learning (n=5 Ctrl, 5 DMS^pdyn-cKO^; Ctrl - Simple Linear Regression, R^2^=0.3544, *p=0*.*0192\**, F=7.317. DMS^pdyn-cKO^ – Simple Linear Regression, R^2^=0.1146, *p>0*.*05*, F=1.683. Difference in Intercepts – *p=0*.*04*^*#*^). Middle – Inactive Nosepokes during operant learning (n=9 Ctrl, 12 DMS^pdyn-cKO^ mice; Ctrl - Simple Linear Regression, R^2^=0.06995, *p>0*.*05* F=0.9777). DMS^pdyn-cKO^ – Simple Linear Regression, R^2^=0.04547, *p>0*.*05* F=0.6193). Right – Rewards consumed during operant learning (n=9 Ctrl, 12 DMS^pdyn-cKO^ mice; Ctrl - Simple Linear Regression, R^2^=0.3460, *p=0*.*0211\**, F=6.878. dyn-cKO – Simple Linear Regression, R^2^=0.06238, *p>0*.*05*, F=0.8649. Difference in Intercepts – *p=0*.*0012*^*#*^). (**L**) Left – Active Nosepokes during extinction learning (n=5 Ctrl, 5 DMS^pdyn-cKO^ mice; Ctrl - Simple Linear Regression, R^2^=0.8160, *p<0*.*0001\*\*\*\**, F=57.65. DMS^pdyn-cKO^ – Simple Linear Regression, R^2^=0.4654, *p=0*.*0051\*\**, F=11.32. Difference in Slopes – *p=0*.*0026\*\**). Middle – Inactive Nosepokes during operant learning (n=9 Ctrl, 12 DMS^pdyn-cKO^ mice; Ctrl - Simple Linear Regression, R^2^=0.09804, *p>0*.*05*, F=1.413. DMS^pdyn-cKO^ – Simple Linear Regression, R^2^=0.4345, *p=0*.*0075\** F=9.989). Right – Rewards consumed during operant learning (n=9 Ctrl, 12 DMS^pdyn-cKO^ mice; Ctrl - Simple Linear Regression, R^2^=0.8084, *p<0*.*0001\*\*\*\**, F=54.84. DMS^pdyn-cKO^ – Simple Linear Regression, R^2^=0.5465, *p=0*.*0016\*\**, F=15.67. Difference in Slopes – *p=0*.*0389\**). (**M**) Left – Active Nosepokes in Ctrl and DMS^pdyn-cKO^ mice with U50,488 injection (n=5 Ctrl, 5 DMS^pdyn-cKO^ mice; Two Way ANOVA, genotype x treatment *p<0*.*0266\**, F(1,8)=7.358. Multiple comparisons – Ctrl veh vs. U50, *p>0*.*05;* DMS^pdyn-cKO^ veh vs. U50, *p=0*.*0039\*\**). Middle – Inactive Nosepokes in Ctrl and DMS^pdyn-cKO^ mice with U50,488 injection (n=5 Ctrl, 5 DMS^pdyn-cKO^ mice; Two Way ANOVA, genotype x treatment *p>0*.*05*). Right – Rewards consumed in Ctrl and DMS^pdyn-cKO^ mice with U50,488 injection (n=5 Ctrl, 5 DMS^pdyn-cKO^ mice; Two Way ANOVA, genotype x treatment *p<0*.*0318\**, F(1,8)=6.739. Multiple comparisons – Ctrl veh vs. U50, *p>0*.*05;* DMS^pdyn-cKO^ veh vs. U50, *p=0*.*0214\**).

**Supplementary Figure 4:**
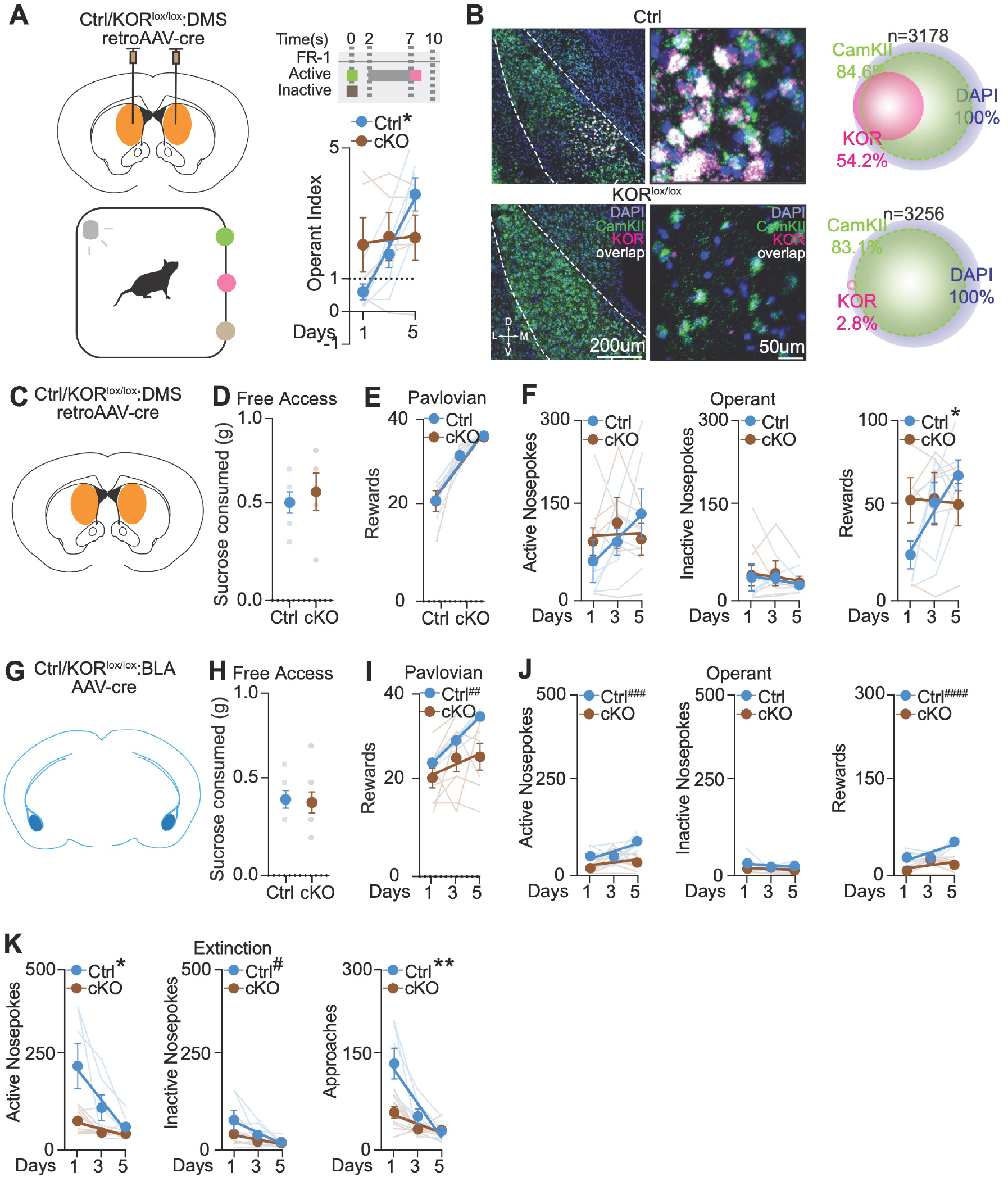
BLA KOR-expressing neurons preferentially project to DMS D1/dyn neurons and are necessary for goal-directed behavior. (**A**) Left - Schematic of viral injection in the DMS and schematic of operant behavior. Right - Operant learning (n=6 Ctrl, 5 DMS^retroKOR-cKO^ mice; Ctrl - Simple Linear Regression, R^2^=0.6427, *p<0*.*0001\*\*\*\**, F=28.78. DMS^retroKOR-cKO^ – Simple Linear Regression, R^2^=0.003458, *p>0*.*05*, F=0.04629. Difference in Slopes – *p=0*.*0193\**). (**B**) Top - 20X (left) and 40X (middle) Confocal image of **Control** BLA section with ISH for DAPI, CamKII, KOR and overlap, and (right) Quantification of ISH. Bottom - 20X (left) and 40X (middle) Confocal image of **DMS**^**retroKOR-cKO**^ BLA section with ISH for DAPI, CamKII, KOR and overlap, and (right) Quantification of ISH. (**C**) Schematic of injection site in the DMS. (**D**) Sucrose consumed in homecage (n=6 Ctrl, 5 DMS^retroKOR-cKO^ mice; unpaired t test, *p>0*.*05)*. (**E**) Rewards consumed during Pavlovian learning (n=5 Ctrl, 6 DMS^retroKOR-cKO^ mice; Ctrl - Simple Linear Regression, R^2^=0.8757, *p<0*.*0001\*\*\*\**, F=112.7. DMS^retroKOR-cKO^ – Simple Linear Regression, R^2^=0.1292, *p=0*.*0001\*\*\*\**, F=44.93). (**F**) Left – Active Nosepokes during operant learning (n=5 Ctrl, 6 DMS^retroKOR-cKO^ mice; Ctrl - Simple Linear Regression, R^2^=0.1497, *p>0*.*05*, F=2.817. DMS^retroKOR-cKO^ – Simple Linear Regression, R^2^=0.0005158, *p>0*.*05*, F=0.00678). Middle – Inactive Nosepokes during operant learning (n=5 Ctrl, 6 DMS^retroKOR-cKO^ mice; Ctrl - Simple Linear Regression, R^2^=0.02413, *p>0*.*05*, F=0.3957). DMS^retroKOR-cKO^ – Simple Linear Regression, R^2^=0.02019, *p>0*.*05*, F=0.2679). Right – Rewards consumed during operant learning (n=5 Ctrl, 6 DMS^retroKOR-cKO^ mice; Ctrl - Simple Linear Regression, R^2^=0.4161, *p<0*.*0038\*\**, F=11.40. DMS^retroKOR-cKO^ – Simple Linear Regression, R^2^=0.001594, *p>0*.*05*, F=0.02076. Difference in Slopes – *p=0*.*0394\**). (**G**) Schematic of injection site in the BLA. (**H**) Sucrose consumed in homecage (n=6 Ctrl, 8 BLA^KOR-cKO^ mice; unpaired t test, *p>0*.*05)*. (**I**) Rewards consumed during Pavlovian learning (n=6 Ctrl, 8 BLA^KOR-cKO^ mice; Ctrl - Simple Linear Regression, R^2^=0.9110, *p<0*.*0001\*\*\*\**, F=163.9. BLA^KOR-cKO^ – Simple Linear Regression, R^2^=0.06236, *p>0*.*05*, F=1.463. Difference in Intercepts – *p=0*.*0066*^*#*^). (**J**) Left – Active Nosepokes during operant learning (n=6 Ctrl, 8 KOR-cKO mice; Ctrl - Simple Linear Regression, R^2^=0.3383, *p=0*.*0113\**, F=8.180. Difference in Intercepts – *p=0*.*0002*^*##*^). Middle – Inactive Nosepokes during operant learning (n=6 Ctrl, 8 BLA^KOR-cKO^ mice; Ctrl - Simple Linear Regression, R^2^=0.04630, *p>0*.*05*, F=0.7773. KOR-cKO – Simple Linear Regression, R^2^=0.03250, *p>0*.*05*, F=0.7389). Right – Rewards consumed during operant learning (n=5 Ctrl, 6 BLA^KOR-cKO^ mice; Ctrl - Simple Linear Regression, R^2^=0.4868, *p<0*.*0013\*\**, F=15.17. KOR-cKO – Simple Linear Regression, R^2^=0.1290, *p>0*.*05*, F=3.259. Difference in Intercepts – *p<0*.*0001*^*##*^). (**K**) Left – Active Nosepokes during operant learning (n=6 Ctrl, 8 BLA^KOR-cKO^ mice; Ctrl - Simple Linear Regression, R^2^=0.3358, *p=0*.*0117\**, F=8.088. BLA^KOR-cKO^ – Simple Linear Regression, R^2^=0.3369, *p=0*.*0029\**, F=11.18. Difference in Slopes – *p=0*.*0154\**). Middle – Inactive Nosepokes during operant learning (n=6 Ctrl, 8 BLA^KOR-cKO^ mice; Ctrl - Simple Linear Regression, R^2^=0.2867, *p=0*.*0220\**, F=6.430. BLA^KOR-cKO^ – Simple Linear Regression, R^2^=0.2991, *p=0*.*0057\**, F=9.389. Difference in Intercepts – *p=0*.*0432*^*#*^). Right – Approaches during extinction learning (n=6 Ctrl, 8 BLA^KOR-cKO^ mice; Ctrl - Simple Linear Regression, R^2^=0.5617, *p<=0*.*0003\*\*\**, F=20.51. BLA^KOR-cKO^ – Simple Linear Regression, R^2^=0.3075, *p=0*.*0049\*\**, F=9.770. Difference in Slopes – *p=0*.*0013\*\**).

**Supplementary Figure 5:**
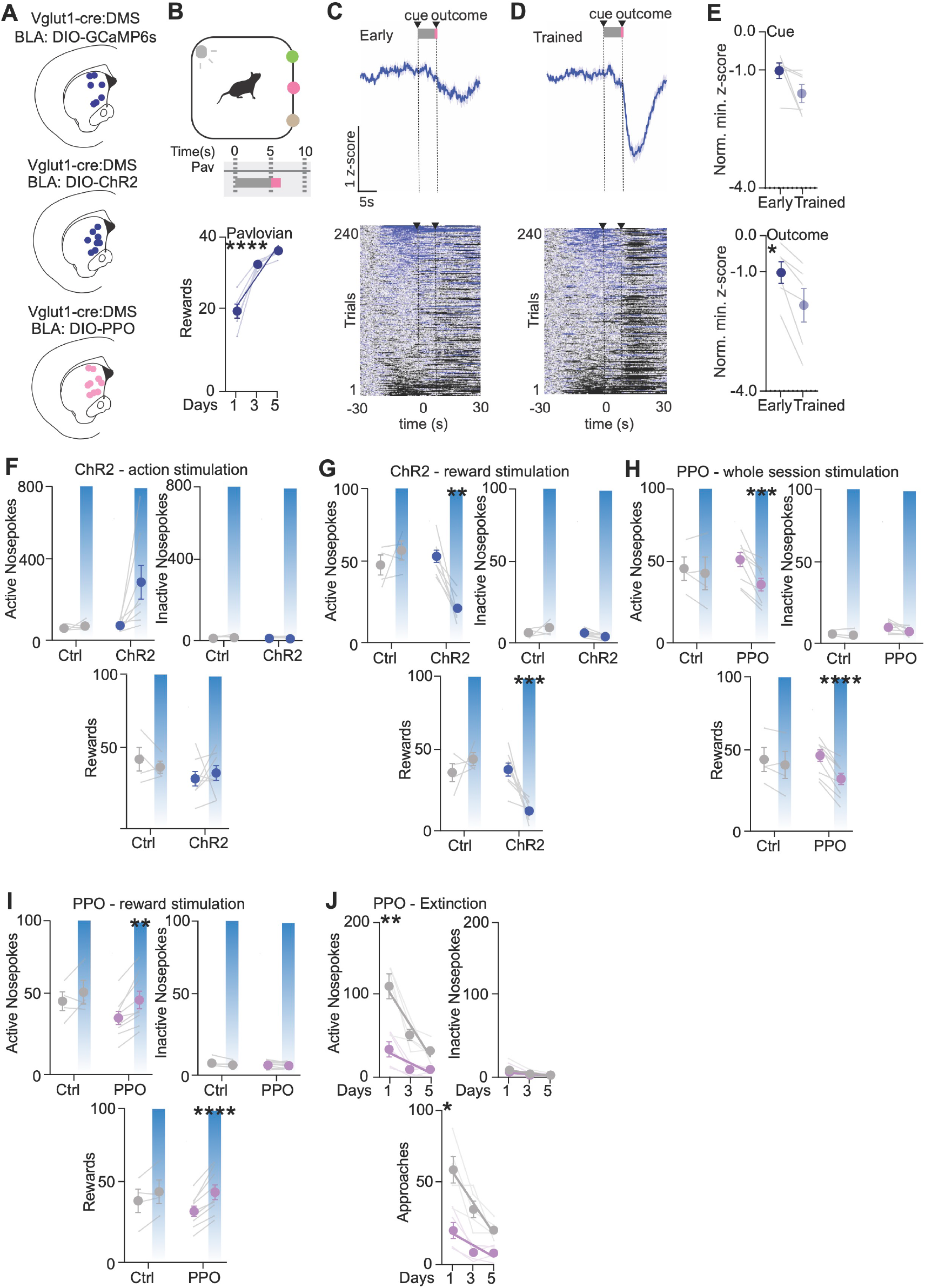
BLA-DMS terminals are engaged during and necessary, and sufficient for goal-directed behavior. (**A**) Top – Placement map of optic fiber implants for photometry in the DMS. Middle – Placement map of optic fiber implants for photoactivation in the DMS. Bottom – Placement map of optic fiber implants for photoinhibition in the DMS. (**B**) Top - Pavlovian behavior schedule during photometry. Bottom – Rewards consumed during Pavlovian learning (n=6mice; Simple Linear Regression, R^2^=0.7959, *p<0*.*0001\*\*\*\**, F=62.41). (**C**) Mean fluorescence and heatmap raster plots during early pavlovian behavior (n=6mice). (**D**) Mean fluorescence and heatmap raster plots during trained pavlovian behavior (n=6mice). (**E**) Top - Normalized Minimum z-score values of early and trained during **cue** period (0-5s) (n=6mice; paired t test, *p>0*.*05*). Bottom - Normalized Minimum z-score values of early and trained during **reward** period (7-30s) (n=6mice; *p=0*.*0392*\*, paired t test, t=2.774, df=5). (**F**) Left - Active Nosepokes during optogenetic stimulation at **action** in Ctrl and vglut1-cre mice (n=4 Ctrl,8 vglut1-cre mice; Two Way ANOVA, stim x genotype *p>0*.*05*). Right - Inactive Nosepokes during optogenetic stimulation at **action** in Ctrl and vglut1-cre mice (n=4 Ctrl,8 vglut1-cre mice; Two Way ANOVA, stim x genotype *p>0*.*05*). Bottom - Rewards consumed during optogenetic stimulation at **action** in Ctrl and vglut1-cre mice (n=4 Ctrl,8 vglut1-cre mice; Two Way ANOVA, stim x genotype *p>0*.*05*). (**G**) Left - Active Nosepokes during optogenetic stimulation at **cue and reward** in Ctrl and vglut1-cre mice (n=4 Ctrl,8 vglut1-cre mice; Two Way ANOVA, stim x genotype *p=0*.*0015\*\**, F(1,10)=18.59. Multiple comparisons – ctrl off vs. on, *p>0*.*05;* vglut1-cre off vs. on, *p=0*.*0003\*\*\**). Right - Inactive Nosepokes during optogenetic stimulation at **cue and reward** in Ctrl and vglut1cre mice (n=4 Ctrl,8 vglut1-cre mice; Two Way ANOVA, stim x genotype *p>0*.*05*). Bottom – Rewards consumed during optogenetic stimulation at **cue and reward** in Ctrl and vglut1-cre mice (n=4 Ctrl,8 vglut1-cre mice; Two Way ANOVA, stim x genotype *p=0*.*0081\*\**, F(1,10)=10.83. Multiple comparisons – ctrl off vs. on, *p>0*.*05;* vglut1-cre off vs. on, *p=0*.*0031\*\**). (**H**) Left - Active Nosepokes during optogenetic inhibition at **whole session** in Ctrl and vglut1-cre mice ((n=4 Ctrl,9 vglut1-cre mice; Two Way ANOVA, stim x genotype *p=0*.*0384\**, F(1,11)=5.529. Multiple comparisons – ctrl off vs. on, *p>0*.*05;* vglut1-cre off vs. on, *p=0*.*0006\*\*\**). Right - Inactive Nosepokes during optogenetic stimulation at **whole session** in Ctrl and vglut1- cre mice (n=4 Ctrl,9 vglut1-cre mice; Two Way ANOVA, stim x genotype *p>0*.*05*). Bottom - Rewards consumed during optogenetic stimulation at **whole session** in Ctrl and vglut1-cre mice ((n=4 Ctrl,9 vglut1-cre mice; Two Way ANOVA, stim x genotype *p=0*.*0082\*\**, F(1,11)=10.34. Multiple comparisons – ctrl off vs. on, *p>0*.*05;* vglut1-cre off vs. on, *p<0*.*0001\*\*\*\**). (**I**) Left - Active Nosepokes during optogenetic stimulation at **cue and reward** in Ctrl and vglut1-cre mice (n=4 Ctrl,9 vglut1-cre mice; Two Way ANOVA, stim *p=0*.*0044\*\**, F(1,11)=12.77, stim x genotype *p>0*.*05*. Multiple comparisons – ctrl off vs. on, *p>0*.*05;* vglut1-cre off vs. on, *p=0*.*0026\*\**). Right - Inactive Nosepokes during optogenetic stimulation at **cue and reward** in Ctrl and vglut1-cre mice (n=4 Ctrl,9 vglut1-cre mice; Two Way ANOVA, stim x genotype *p>0*.*05*). Bottom – Rewards consumed during optogenetic stimulation at **cue and reward** in Ctrl and vglut1-cre mice (n=4 Ctrl,9 vglut1-cre mice; Two Way ANOVA, stim *p=0*.*0003\*\*\**, F(1,11)=28.09, stim x genotype *p>0*.*05*. Multiple comparisons – ctrl off vs. on, *p>0*.*05;* vglut1-cre off vs. on, *p<0*.*0001\*\*\*\**). (**J**) Left – Active Nosepokes during optogenetic inhibition for extinction at **whole session** (n=5 vglut1-cre mice; Off - Simple Linear Regression, R^2^=0.6730, *p=0*.*0002\*\*\**, F=26.75. On – Simple Linear Regression, R^2^=0.3776, *p=0*.*0148\**, F=7.886. Difference in Slopes – *p=0*.*0048\*\**). Right – Inactive Nosepokes during optogenetic inhibition for extinction at **whole session** (n=5 vglut1-cre mice; Off - Simple Linear Regression, R^2^=0.2174, *p>0*.*05*, F=3.611. On – Simple Linear Regression, R^2^=0.1300, *p>0*.*05*, F=1.942). Bottom – Approaches to reward port during optogenetic inhibition for extinction at **whole session** (n=5 vglut1- cre mice; Off - Simple Linear Regression, R^2^=0.6210, *p=0*.*0005\*\*\**, F=21.30. On – Simple Linear Regression, R^2^=0.3793, *p=0*.*0145\**, F=7.945. Difference in Slopes – *p=0*.*0209\**).

**Supplementary Figure 6:**
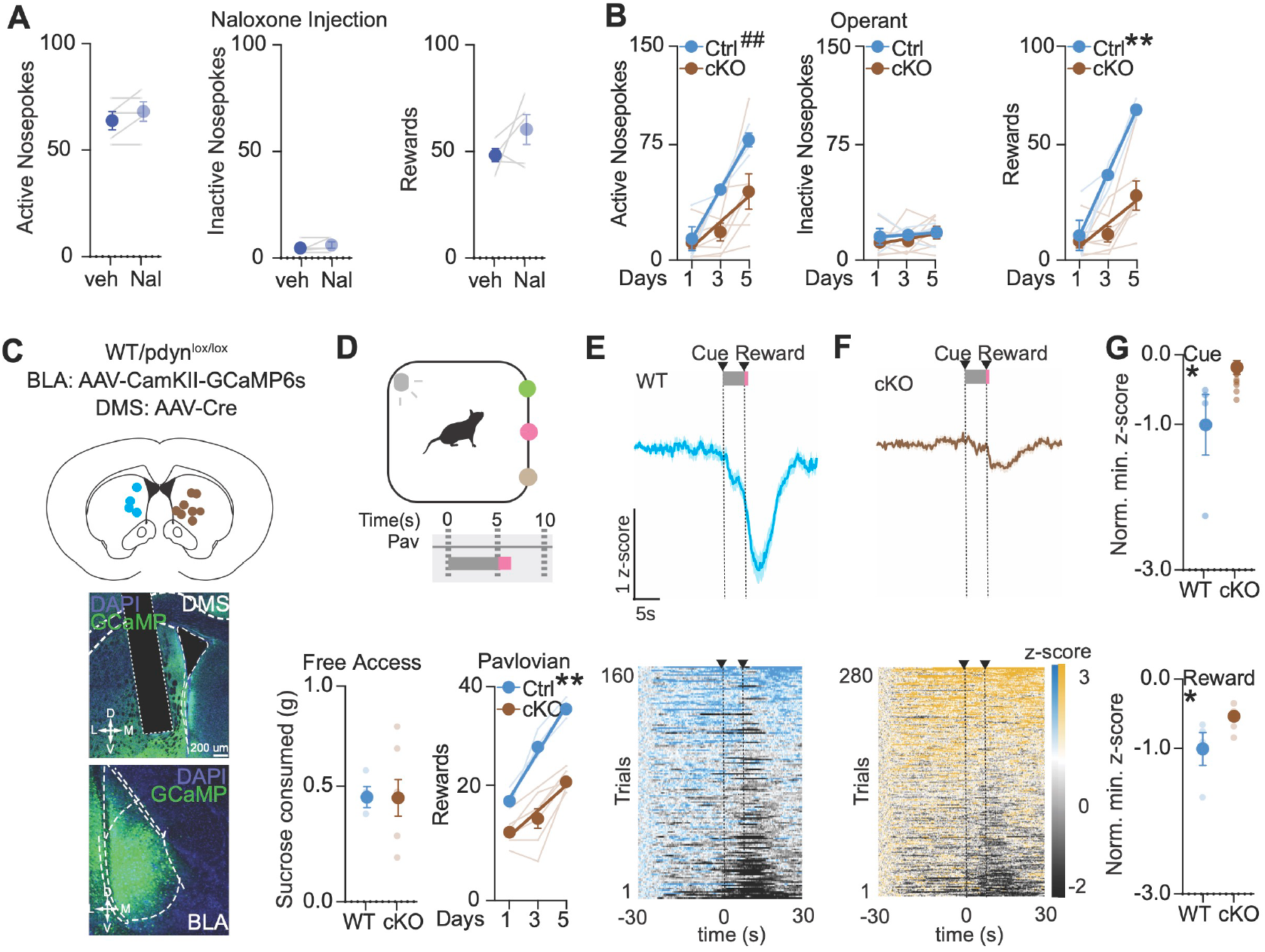
Dynorphin-KOR signaling at BLA-DMS terminals is necessary for goal-directed behavior. (**A**) Left - Active nose- pokes during vehicle vs. Naloxone (n=5mice; paired t test, *p>0*.*05*). Middle - Inactive nosepokes during vehicle vs. Naloxone (n=5mice; paired t test, *p>0*.*05*). Right – Rewards consumed during vehicle vs. Naloxone (n=5mice; paired t test, *p>0*.*05*). (**B**) Top – Placement maps of optic fiber implants in the DMS. Middle - 20X Confocal image of optic fiber implant in the DMS and cells stained with DAPI, with BLA expressing GCaMP6s. Bottom - 20X Confocal image of BLA and cells stained with DAPI, with projection cells expressing GCaMP6s. (**C**) Top - Pavlovian behavior schedule during photometry. Bottom left – Sucrose consumed in homecage (n=4 Ctrl, 7 DMS^pdyn-cKO^ mice; unpaired t test, *p>0*.*05)*. Bottom right - Rewards consumed during Pavlovian learning (n=4 Ctrl, 7 DMS^pdyn-cKO^ mice; Ctrl - Simple Linear Regression, R^2^=0.9423, *p<0*.*0001\*\*\*\**, F=163.3. dyn-cKO – Simple Linear Regression, R^2^=0.5853, *p<0*.*0001\*\*\*\**, F=26.82. Difference in Slopes – *p=0*.*0061\*\**). (**D**) Mean fluorescence and heatmap raster plots of Ctrl during trained pavlovian behavior (n=4mice). (**E**) Mean fluorescence and heatmap raster plots of DMS^pdyn-cKO^ during trained pavlovian behavior (n=7mice). (**F**) Top - Normalized Minimum z-score values of Ctrl and DMS^pdyn-cKO^ during **cue** period (0-5s) (n=4 Ctrl, 7 DMS^pdyn-cKO^ mice; *p=0*.*0389*\*, unpaired t test, t=2.416, df=9). Bottom - Normalized Minimum z-score values during **reward** period (7-30s) (n=4 Ctrl, 7 DMS^pdyn-cKO^ mice; *p=0*.*0392*\*, unpaired t test, t=2.411, df=9). (**G**) Left – Active Nosepokes during operant behavior (n=4 Ctrl, 7 DMS^pdyn-cKO^ mice; Ctrl - Simple Linear Regression, R^2^=0.8944, *p<0*.*0001\*\*\*\**, F=84.71. DMS^pdyn-cKO^ – Simple Linear Regression, R^2^=0.3190, *p=0*.*0076\*\**, F=8.900. Difference in Intercepts – *p=0*.*0038*^*#*^). Middle – Inactive Nosepokes during operant behavior (n=4 Ctrl, 7 DMS^pdyn-cKO^ mice; Ctrl - Simple Linear Regression, R^2^=0.02771, *p>0*.*05*, F=0.2875. DMS^pdyn-cKO^ – Simple Linear Regression, R^2^=0.06895, *p>0*.*05*, F=1.407). Right – Rewards consumed during operant behavior (n=4 Ctrl, 7 DMS^pdyn-cKO^ mice; Ctrl - Simple Linear Regression, R^2^=0.9036, *p<0*.*0001\*\*\*\**, F=93.72. DMS^pdyn-cKO^ – Simple Linear Regression, R^2^=0.3357, *p=0*.*0059\*\**, F=9.600. Difference in Slopes – *p=0*.*0011\*\**).

**Supplementary Figure 7:**
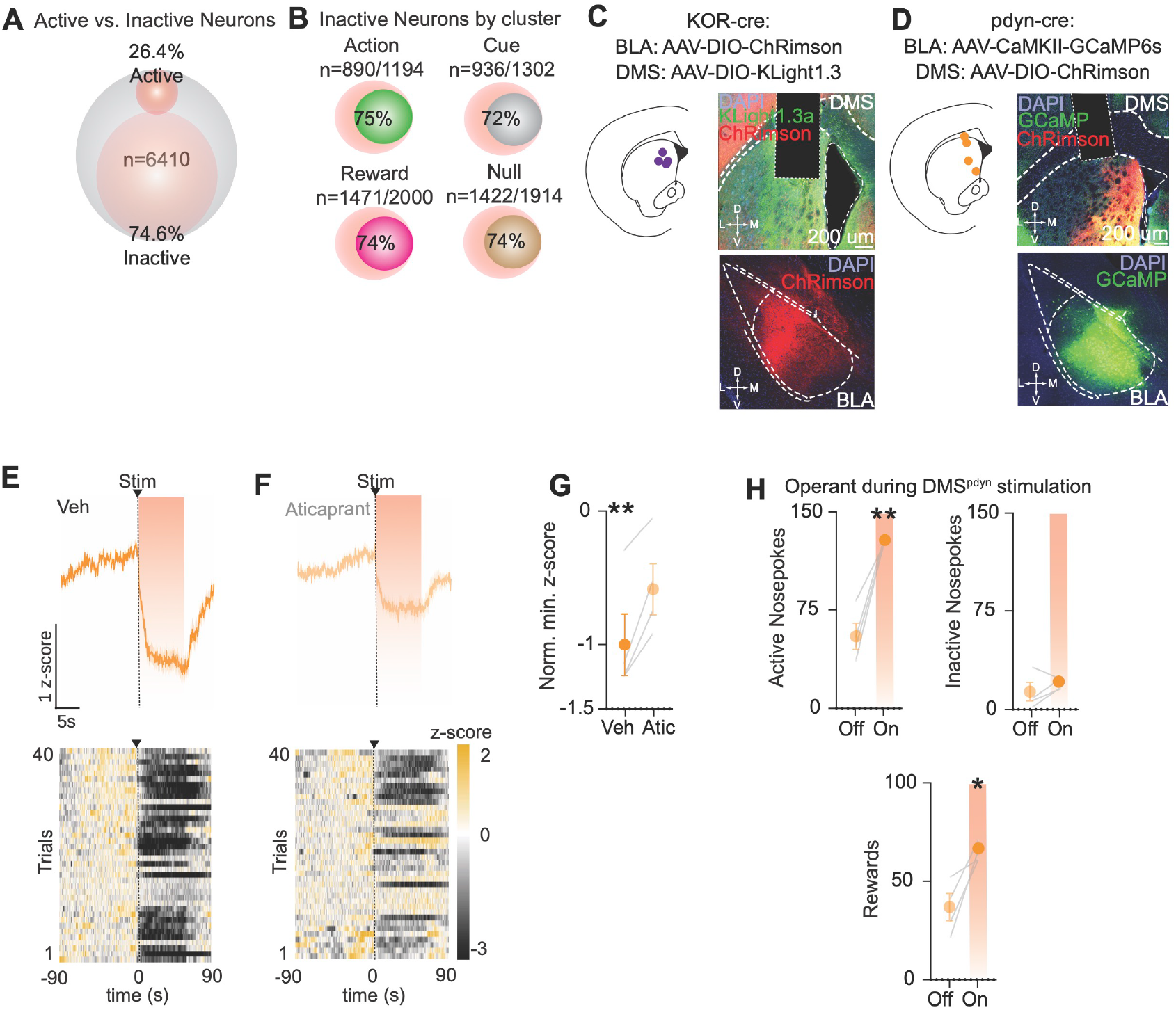
BLA terminal activity engenders DMS dynorphin neuron activity, dynorphin release and subsequent dynorphin-KOR signaling to promote goal-directed behavior. (**A**) Quantification of the % of active and inactive neurons following sequential spiral stimulation of BLA axons. (**B**) Quantification of % of inactive neurons following stimulation enriched in each cluster. (**C**) Left – Placement maps of optic fiber implants in the DMS. Right top - 20X Confocal image of optic fiber implant in the DMS and cells stained with DAPI, with DMS expressing kLight 1.3a and BLA terminals expressing ChRimson. Right bottom - 20X Confocal image of BLA and cells stained with DAPI, with projection cells expressing ChRimson. (**D**) Top – Placement maps of optic fiber implants in the DMS. Right top - 20X Confocal image of optic fiber implant in the DMS and cells stained with DAPI, with DMS dyn neurons expressing ChRimson, and BLA terminals expressing GCaMP6s. Right bottom - 20X Confocal image of BLA and cells stained with DAPI, with projection cells expressing GCaMP6s. (**E**) Mean fluorescence and heatmap raster plots following stimulation with vehicle injection (n=4mice). (**F**) Mean fluorescence and heatmap raster plots following stimulation with aticaprant injection (n=4mice). (**G**) Normalized Peak z-score values of vehicle vs. Aticaprant following **stim** period (0-60s) (n=4mice; paired t test, *p=0*.*0154\**, t=4.996, df=3). (**H**) Left - Active nosepokes following stimulation (n=4mice; paired t test, *p=0*.*0059\*\**, t=7.039, df=3). Right - Inactive nosepokes following stimulation (n=4mice; paired t test, *p>0*.*05*). Bottom – Rewards consumed following stimulation (n=4mice; paired t test, *p=0*.*031\**, t=3.893, df=3).

## REFERENCES

Abraham, A.D., Casello, S.M., Schattauer, S.S., Wong, B.A., Mizuno, G.O., Mahe, K. et al. (2021) Release of endogenous dynorphin opioids in the prefrontal cortex disrupts cognition. Neuropsychopharmacology, 46(13), 2330–2339. doi:10.1038/s41386-021-01168-2. URL https://www.ncbi.nlm.nih.gov/pmc/articles/PMC8580977/

Al-Hasani, R. & Bruchas, M.R. (2011) Molecular mechanisms of opioid receptor-dependent signaling and behavior. Anesthesiology, 115(6), 1363–1381. doi:10.1097/ALN.0b013e318238bba6.

Al-Hasani, R., Gowrishankar, R., Schmitz, G.P., Pedersen, C.E., Marcus, D.J., Shirley, S.E. et al. (2021) Ventral tegmental area GABAergic inhibition of cholinergic interneurons in the ventral nucleus accumbens shell promotes reward reinforcement. Nat Neurosci, 24(10), 1414–1428. doi:10.1038/s41593-021-00898-2.

Al-Hasani, R., Wong, J.M.T., Mabrouk, O.S., McCall, J.G., Schmitz, G.P., Porter-Stransky, K.A. et al. (2018) In vivo detection of optically-evoked opioid peptide release. eLife, 7, e36520. doi:10.7554/eLife.36520, publisher: eLife Sciences Publications, Ltd. URL https://doi.org/10.7554/eLife.36520

Atwood, B.K., Lovinger, D.M. & Mathur, B.N. (2014) Presynaptic long-term depression mediated by Gi/o-coupled receptors. Trends in Neurosciences, 37(11), 663–673. doi:10.1016/j.tins.2014.07.010, publisher: Elsevier. URL https://www.cell.com/trends/neurosciences/abstract/S0166-2236(14)00126-X

Balleine, B.W., Delgado, M.R. & Hikosaka, O. (2007) The Role of the Dorsal Striatum in Reward and Decision-Making. J Neurosci, 27(31), 8161–8165. doi:10.1523/JNEUROSCI.1554-07.2007. URL https://www.ncbi.nlm.nih.gov/pmc/articles/PMC6673072/

Bals-Kubik, R., Ableitner, A., Herz, A. & Shippenberg, T.S. (1993) Neuroanatomical sites mediating the motivational effects of opioids as mapped by the conditioned place preference paradigm in rats. J Pharmacol Exp Ther, 264(1), 489–495. publisher: American Society for Pharmacology and Experimental Therapeutics. URL https://jpet.aspetjournals.org/content/264/1/489

Baxter, M.G. & Murray, E.A. (2002) The amygdala and reward. Nat Rev Neurosci, 3(7), 563–573. doi:10.1038/nrn875, publisher: Nature Publishing Group. URL https://www.nature.com/articles/nrn875

Bloem, B., Huda, R., Amemori, K.i., Abate, A.S., Krishna, G., Wilson, A.L. et al. (2022) Multiplexed action-outcome representation by striatal striosome-matrix compartments detected with a mouse cost-benefit foraging task. Nat Commun, 13, 1541. doi:10.1038/s41467-022-28983-5. URL https://www.ncbi.nlm.nih.gov/pmc/articles/PMC8941061/

Bloem, B., Huda, R., Sur, M. & Graybiel, A.M. (2017) Two-photon imaging in mice shows striosomes and matrix have overlapping but differential reinforcement-related responses. eLife, 6, e32353. doi:10.7554/eLife.32353, publisher: eLife Sciences Publications, Ltd. URL https://doi.org/10.7554/eLife.32353

Bruchas, M.R., Land, B.B. & Chavkin, C. (2010) The dynorphin/kappa opioid system as a modulator of stress-induced and pro-addictive behaviors. Brain Res, 1314, 44–55. doi:10.1016/j.brainres.2009.08.062.

Castro, D.C., Oswell, C.S., Zhang, E.T., Pedersen, C.E., Piantadosi, S.C., Rossi, M.A. et al. (2021) An endogenous opioid circuit determines state-dependent reward consumption. Nature, 598(7882), 646–651. doi:10.1038/s41586-021-04013-0, publisher: Nature Publishing Group. URL https://www.nature.com/articles/s41586-021-04013-0

Chavkin, C., James, I.F. & Goldstein, A. (1982) Dynorphin is a specific endogenous ligand of the kappa opioid receptor. Science, 215(4531), 413–415. doi:10.1126/science.6120570.

Cho, Y.T., Ernst, M. & Fudge, J.L. (2013) Cortico–Amygdala– Striatal Circuits Are Organized as Hierarchical Subsystems through the Primate Amygdala. J Neurosci, 33(35), 14017–14030. doi:10.1523/JNEUROSCI.0170-13.2013. URL https://www.ncbi.nlm.nih.gov/pmc/articles/PMC3756751/

Copits, B.A., Gowrishankar, R., O’Neill, P.R., Li, J.N., Girven, K.S., Yoo, J.J. et al. (2021) A photoswitchable GPCR-based opsin for presynaptic inhibition. Neuron, 109(11), 1791–1809.e11. doi:10.1016/j.neuron.2021.04.026.

Corbit, L.H. & Janak, P.H. (2010) Posterior dorsomedial striatum is critical for both selective instrumental and Pavlovian reward learning. Eur J Neurosci, 31(7), 1312–1321. doi:10.1111/j.1460-9568.2010.07153.x.

Corbit, L.H., Leung, B.K. & Balleine, B.W. (2013) The Role of the Amygdala-Striatal Pathway in the Acquisition and Performance of Goal-Directed Instrumental Actions. J Neurosci, 33(45), 17682–17690. doi:10.1523/JNEUROSCI.3271-13.2013. URL https://www.ncbi.nlm.nih.gov/pmc/articles/PMC6618427/

Corder, G., Castro, D.C., Bruchas, M.R. & Scherrer, G. (2018) Endogenous and Exogenous Opioids in Pain. Annual Review of Neuroscience, 41(Volume 41, 2018), 453–473. doi:10.1146/annurev-neuro-080317-061522, publisher: Annual Reviews. URL https://www.annualreviews.org/content/journals/10.1146/annurev-neuro-080317-061522

Courtin, J., Bitterman, Y., Müller, S., Hinz, J., Hagihara, K.M., Müller, C. et al. (2022) A neuronal mechanism for motivational control of behavior. Science, 375(6576), eabg7277. doi:10.1126/science.abg7277, publisher: American Association for the Advancement of Science. URL https://www.science.org/doi/full/10.1126/science.abg7277

Crowley, N.A., Bloodgood, D.W., Hardaway, J.A., Kendra, A.M., McCall, J.G., Al-Hasani, R. et al. (2016) Dynorphin Controls the Gain of an Amygdalar Anxiety Circuit. Cell Rep, 14(12), 2774–2783. doi:10.1016/j.celrep.2016.02.069.

Drago, J., Gerfen, C.R., Lachowicz, J.E., Steiner, H., Hollon, T.R., Love, P.E. et al. (1994) Altered striatal function in a mutant mouse lacking D1A dopamine receptors. Proceedings of the National Academy of Sciences, 91(26), 12564–12568. doi:10.1073/pnas.91.26.12564, publisher: Proceedings of the National Academy of Sciences. URL https://www.pnas.org/doi/abs/10.1073/pnas.91.26.12564

Drake, C.T., Patterson, T.A., Simmons, M.L., Chavkin, C. & Milner, T.A. (1996) Kappa opioid receptor-like immunoreactivity in guinea pig brain: ultrastructural localization in presynaptic terminals in hippocampal formation. J Comp Neurol, 370(3), 377–395. doi:10.1002/(SICI)1096-9861(19960701)370:3<377::AID-CNE8>3.0.CO;2-1.

Ehrich, J.M., Phillips, P.E.M. & Chavkin, C. (2014) Kappa Opioid Receptor Activation Potentiates the Cocaine-Induced Increase in Evoked Dopamine Release Recorded In Vivo in the Mouse Nucleus Accumbens. Neuropsychopharmacol, 39(13), 3036–3048. doi:10.1038/npp.2014.157, publisher: Nature Publishing Group. URL https://www.nature.com/articles/npp2014157

Fagergren, P., Smith, H.R., Daunais, J.B., Nader, M.A., Porrino, L.J. & Hurd, Y.L. (2003) Temporal upregulation of prodynorphin mRNA in the primate striatum after cocaine self-administration. European Journal of Neuroscience, 17(10), 2212–2218. doi:10.1046/j.1460-9568.2003.02636.x, _eprint: https://onlinelibrary.wiley.com/doi/pdf/10.1046/j.1460-9568.2003.02636.x. URL https://onlinelibrary.wiley.com/doi/abs/10.1046/j.1460-9568.2003.02636.x

Farahbakhsh, Z.Z., Song, K., Branthwaite, H.E., Erickson, K.R., Mukerjee, S., Nolan, S.O. et al. (2023) Systemic kappa opioid receptor antagonism accelerates reinforcement learning via augmentation of novelty processing in male mice. Neuropsychopharmacol., 48(6), 857–868. doi:10.1038/s41386-023-01547-x, publisher: Nature Publishing Group. URL https://www.nature.com/articles/s41386-023-01547-x

Gerfen, C.R., Engber, T.M., Mahan, L.C., Susel, Z., Chase, T.N., Monsma, F.J. et al. (1990) D1 and D2 dopamine receptor-regulated gene expression of striatonigral and striatopallidal neurons. Science, 250(4986), 1429–1432. doi:10.1126/science.2147780.

Gerfen, C.R. & Surmeier, D.J. (2011) Modulation of striatal projection systems by dopamine. Annu Rev Neurosci, 34, 441–466. doi:10.1146/annurev-neuro-061010-113641.

Giovanniello, J.R., Paredes, N., Wiener, A., Ramírez-Armenta, K., Oragwam, C., Uwadia, H.O. et al. (2023) A dual-pathway architecture enables chronic stress to promote habit formation. Pages: 2023.10.03.560731 Section: New Results. URL https://www.biorxiv.org/content/10.1101/2023.10.03.560731v1

Gordon-Fennell, A., Barbakh, J.M., Utley, M.T., Singh, S., Bazzino, P., Gowrishankar, R. et al. (2023) An open-source platform for headfixed operant and consummatory behavior. eLife, 12, e86183. doi:10.7554/eLife.86183, publisher: eLife Sciences Publications, Ltd. URL https://doi.org/10.7554/eLife.86183

Gordon-Fennell, L., Farero, R.D., Burgeno, L.M., Murray, N.L., Abraham, A.D., Soden, M.E. et al. (2023) Kappa Opioid Receptors in Mesolimbic Terminals Mediate Escalation of Cocaine Consumption. Pages: 2023.12.21.572842 Section: New Results. URL https://www.biorxiv.org/content/10.1101/2023.12.21.572842v1

Gremel, C.M. & Costa, R.M. (2013) Orbitofrontal and striatal circuits dynamically encode the shift between goal-directed and habitual actions. Nat Commun, 4(1), 2264. doi:10.1038/ncomms3264, publisher: Nature Publishing Group. URL https://www.nature.com/articles/ncomms3264

Hikosaka, O., Sakamoto, M. & Usui, S. (1989) Functional properties of monkey caudate neurons. III. Activities related to expectation of target and reward. J Neurophysiol, 61(4), 814–832. doi:10.1152/jn.1989.61.4.814.

Hirokawa, J., Vaughan, A., Masset, P., Ott, T. & Kepecs, A. (2019) Frontal cortex neuron types categorically encode single decision variables. Nature, 576(7787), 446–451. doi:10.1038/s41586-019-1816-9.

Hjelmstad, G.O. & Fields, H.L. (2003) Kappa Opioid Receptor Activation in the Nucleus Accumbens Inhibits Glutamate and GABA Release Through Different Mechanisms. Journal of Neurophysiology, 89(5), 2389–2395. doi:10.1152/jn.01115.2002, publisher: American Physiological Society. URL https://journals.physiology.org/doi/full/10.1152/jn.01115.2002

Hjort, M.M., Gowrishankar, R., Tian, L., Gordon-Fennell, A., Namboodiri, V.M.K., Bruchas, M.R. et al. (2024) Microprisms enable enhanced throughput and resolution for longitudinal tracking of neuronal ensembles in deep brain structures. NPh, 11(3), 033407. doi:10.1117/1.NPh.11.3.033407, publisher: SPIE. URL https://www.spiedigitallibrary.org/journals/neurophotonics/volume-11/issue-3/033407/Microprisms-enable-enhanced-throughput-and-resolution-for-longitudinal-tracking-of/10.1117/1.NPh.11.3.033407.full

Hogarth, L. (2020) Addiction is driven by excessive goal-directed drug choice under negative affect: translational critique of habit and compulsion theory. Neuropsychopharmacol., 45(5), 720–735. doi:10.1038/s41386-020-0600-8, publisher: Nature Publishing Group. URL https://www.nature.com/articles/s41386-020-0600-8

Hurd, Y.L., Brown, E.E., Finlay, J.M., Fibiger, H.C. & Gerfen, C.R. (1992) Cocaine self-administration differentially alters mRNA expression of striatal peptides. Brain Res Mol Brain Res, 13(1-2), 165–170. doi:10.1016/0169-328x(92)90058-j.

Hurd, Y.L. & Herkenham, M. (1993) Molecular alterations in the neostriatum of human cocaine addicts. Synapse, 13(4), 357–369. doi:10.1002/syn.890130408.

Ironside, M., Amemori, K.I., McGrath, C.L., Pedersen, M.L., Kang, M.S., Amemori, S. et al. (2020) Approach-Avoidance Conflict in Major Depressive Disorder: Congruent Neural Findings in Humans and Nonhuman Primates. Biol Psychiatry, 87(5), 399–408. doi:10.1016/j.biopsych.2019.08.022.

Jin, X., Tecuapetla, F. & Costa, R.M. (2014) Basal Ganglia Subcircuits Distinctively Encode the Parsing and Concatenation of Action Sequences. Nat Neurosci, 17(3), 423–430. doi:10.1038/nn.3632. URL https://www.ncbi.nlm.nih.gov/pmc/articles/PMC3955116/

Kelley, A.E., Domesick, V.B. & Nauta, W.J. (1982) The amygdalostriatal projection in the rat–an anatomical study by anterograde and retrograde tracing methods. Neuroscience, 7(3), 615–630. doi:10.1016/0306-4522(82)90067-7.

Kim, J., Zhang, X., Muralidhar, S., LeBlanc, S.A. & Tonegawa, S. (2017) Basolateral to Central Amygdala Neural Circuits for Appetitive Behaviors. Neuron, 93(6), 1464–1479.e5. doi:10.1016/j.neuron.2017.02.034.

Knoll, A.T., Muschamp, J.W., Sillivan, S.E., Ferguson, D., Dietz, D.M., Meloni, E.G. et al. (2011) Kappa opioid receptor signaling in the basolateral amygdala regulates conditioned fear and anxiety in rats. Biol Psychiatry, 70(5), 425–433. doi:10.1016/j.biopsych.2011.03.017.

Kravitz, A.V., Freeze, B.S., Parker, P.R.L., Kay, K., Thwin, M.T., Deisseroth, K. et al. (2010) Regulation of parkinsonian motor behaviours by optogenetic control of basal ganglia circuitry. Nature, 466(7306), 622–626. doi:10.1038/nature09159, publisher: Nature Publishing Group. URL https://www.nature.com/articles/nature09159

Kønig, A.B., Ciriachi, C., Gether, U. & Rickhag, M. (2019) Chemogenetic Targeting of Dorsomedial Direct-pathway Striatal Projection Neurons Selectively Elicits Rotational Behavior in Mice. Neuroscience, 401, 106–116. doi:10.1016/j.neuroscience.2019.01.013, publisher: Elsevier. URL https://www.ibroneuroscience.org/article/S0306-4522(19)30033-8/abstract

Lalive, A.L., Lien, A.D., Roseberry, T.K., Donahue, C.H. & Kreitzer, A.C.) Motor thalamus supports striatum-driven reinforcement. eLife, 7, e34032. doi:10.7554/eLife.34032. URL https://www.ncbi.nlm.nih.gov/pmc/articles/PMC6181560/

Limoges, A., Yarur, H.E. & Tejeda, H.A. (2022) Dynorphin/kappa opioid receptor system regulation on amygdaloid circuitry: Implications for neuropsychiatric disorders. Front. Syst. Neurosci., 16. doi:10.3389/fnsys.2022.963691, publisher: Frontiers. URL https://www.frontiersin.org/articles/10.3389/fnsys.2022.963691

Lowe, S.L., Wong, C.J., Witcher, J., Gonzales, C.R., Dickinson, G.L., Bell, R.L. et al. (2014) Safety, tolerability, and pharmacokinetic evaluation of single- and multiple-ascending doses of a novel kappa opioid receptor antagonist LY2456302 and drug interaction with ethanol in healthy subjects. The Journal of Clinical Pharm-acology, 54(9), 968–978. doi:10.1002/jcph.286, _eprint: https://onlinelibrary.wiley.com/doi/pdf/10.1002/jcph.286. https://www.spiedigitallibrary.org/journals/neurophotonics/volume-11/issue-3/033407/Microprisms-enable-enhanced-throughput-and-resolution-for-longitudinal-tracking-of/10.1117/1.NPh.11.3.033407.full URL https://onlinelibrary.wiley.com/doi/abs/10.1002/jcph.286

Marder, E. (2012) Neuromodulation of neuronal circuits: Back to the future. Neuron, 76(1), 1–11. doi:10.1016/j.neuron.2012.09.010. URL https://www.ncbi.nlm.nih.gov/pmc/articles/PMC3482119/

Matamales, M., McGovern, A.E., Mi, J.D., Mazzone, S.B., Balleine, B.W. & Bertran-Gonzalez, J. (2020) Local D2-to D1-neuron transmodulation updates goal-directed learning in the striatum. Science, 367(6477), 549–555. doi:10.1126/science.aaz5751, publisher: American Association for the Advancement of Science. URL https://www.science.org/doi/10.1126/science.aaz5751

Mu, P., Neumann, P.A., Panksepp, J., Schlüter, O.M. & Dong, Y. (2011) Exposure to Cocaine Alters Dynorphin-Mediated Regulation of Excitatory Synaptic Transmission in Nucleus Accumbens Neurons. Biological Psychiatry, 69(3), 228–235. doi:10.1016/j.biopsych.2010.09.014. URL https://www.sciencedirect.com/science/article/pii/S0006322310009509

Nair, A., Karigo, T., Yang, B., Ganguli, S., Schnitzer, M.J., Linderman, S.W. et al. (2023) An approximate line attractor in the hypothalamus encodes an aggressive state. Cell, 186(1), 178–193.e15. doi:10.1016/j.cell.2022.11.027.

Namboodiri, V.M.K., Otis, J.M., van Heeswijk, K., Voets, E.S., Alghorazi, R.A., Rodriguez-Romaguera, J. et al. (2019) Single-cell activity tracking reveals that orbitofrontal neurons acquire and maintain a long-term memory to guide behavioral adaptation. Nat Neurosci, 22(7), 1110–1121. doi:10.1038/s41593-019-0408-1, publisher: Nature Publishing Group. URL https://www.nature.com/articles/s41593-019-0408-1

Namburi, P., Al-Hasani, R., Calhoon, G.G., Bruchas, M.R. & Tye, K.M. (2016) Architectural Representation of Valence in the Limbic System. Neuropsychopharmacology, 41(7), 1697–1715. doi:10.1038/npp.2015.358.

Nygard, S.K., Hourguettes, N.J., Sobczak, G.G., Carlezon, W.A. & Bruchas, M.R. (2016) Stress-Induced Reinstatement of Nicotine Preference Requires Dynorphin/Kappa Opioid Activity in the Basolateral Amygdala. J. Neurosci., 36(38), 9937–9948. doi:10.1523/JNEUROSCI.0953-16.2016, publisher: Society for Neuroscience Section: Articles. URL https://www.jneurosci.org/content/36/38/9937

O’Doherty, J.P., Deichmann, R., Critchley, H.D. & Dolan, R.J. (2002) Neural responses during anticipation of a primary taste reward. Neuron, 33(5), 815–826. doi:10.1016/s0896-6273(02)00603-7.

Ostlund, S.B. & Balleine, B.W. (2008) Differential Involvement of the Basolateral Amygdala and Mediodorsal Thalamus in Instrumental Action Selection. J. Neurosci., 28(17), 4398–4405. doi:10.1523/JNEUROSCI.5472-07.2008, publisher: Society for Neuroscience Section: Articles. URL https://www.jneurosci.org/content/28/17/4398

Ottenheimer, D.J., Hjort, M.M., Bowen, A.J., Steinmetz, N.A. & Stuber, G.D. (2023) A stable, distributed code for cue value in mouse cortex during reward learning. eLife, 12, RP84604. doi:10.7554/eLife.84604, publisher: eLife Sciences Publications, Ltd. URL https://doi.org/10.7554/eLife.84604

Pan, W.X., Mao, T. & Dudman, J.T. (2010) Inputs to the dorsal striatum of the mouse reflect the parallel circuit architecture of the forebrain. Front Neuroanat, 4, 147. doi:10.3389/fnana.2010.00147.

Parker, K.E., Pedersen, C.E., Gomez, A.M., Spangler, S.M., Walicki, M.C., Feng, S.Y. et al. (2019) A Paranigral VTA Nociceptin Circuit that Constrains Motivation for Reward. Cell, 178(3), 653–671.e19. doi:10.1016/j.cell.2019.06.034.

Parkes, S.L. & Balleine, B.W. (2013) Incentive Memory: Evidence the Basolateral Amygdala Encodes and the Insular Cortex Retrieves Outcome Values to Guide Choice between Goal-Directed Actions. J. Neurosci., 33(20), 8753–8763. doi:10.1523/JNEUROSCI.5071-12.2013, publisher: Society for Neuroscience Section: Articles. URL https://www.jneurosci.org/content/33/20/8753

Peak, J., Chieng, B., Hart, G. & Balleine, B.W. (2020) Striatal direct and indirect pathway neurons differentially control the encoding and updating of goal-directed learning. eLife, 9, e58544. doi:10.7554/eLife.58544, publisher: eLife Sciences Publications, Ltd. URL https://doi.org/10.7554/eLife.58544

Perreault, M.L., Hasbi, A., Alijaniaram, M., Fan, T., Varghese, G., Fletcher, P.J. et al. (2010) The Dopamine D1-D2 Receptor Heteromer Localizes in Dynorphin/Enkephalin Neurons. J Biol Chem, 285(47), 36625–36634. doi:10.1074/jbc.M110.159954. URL https://www.ncbi.nlm.nih.gov/pmc/articles/PMC2978591/

Piantadosi, S.C., Zhou, Z.C., Pizzano, C., Pedersen, C.E., Nguyen, T.K., Thai, S. et al. (2024) Holographic stimulation of opposing amygdala ensembles bidirectionally modulates valence-specific behavior via mutual inhibition. Neuron, 112(4), 593–610.e5. doi:10.1016/j.neuron.2023.11.007. URL https://www.sciencedirect.com/science/article/pii/S0896627323008814

Rappleye, M., Gordon-Fennel, A., Castro, D.C., Matarasso, A.K., Zamorano, C.A., Stine, C. et al. (2022) Opto-MASS: a high-throughput engineering platform for genetically encoded fluorescent sensors enabling all-optical in vivo detection of monoamines and opioids. Pages: 2022.06.01.494241 Section: New Results. URL https://www.biorxiv.org/content/10.1101/2022.06.01.494241v2

Redila, V.A. & Chavkin, C. (2008) Stress-induced reinstatement of cocaine seeking is mediated by the kappa opioid system. Psychopharmacology (Berl), 200(1), 59–70. doi:10.1007/s00213-008-1122-y.

Reed, S.J., Lafferty, C.K., Mendoza, J.A., Yang, A.K., Davidson, T.J., Grosenick, L. et al. (2018) Coordinated Reductions in Excitatory Input to the Nucleus Accumbens Underlie Food Consumption. Neuron, 99(6), 1260–1273.e4. doi:10.1016/j.neuron.2018.07.051.

Reiner, A. & Anderson, K.D. (1990) The patterns of neurotransmitter and neuropeptide co-occurrence among striatal projection neurons: conclusions based on recent findings. Brain Res Brain Res Rev, 15(3), 251–265. doi:10.1016/0165-0173(90)90003-7.

Reinert, S., Hübener, M., Bonhoeffer, T. & Goltstein, P.M. (2021) Mouse prefrontal cortex represents learned rules for categorization. Nature, 593(7859), 411–417. doi:10.1038/s41586-021-03452-z, publisher: Nature Publishing Group. URL https://www.nature.com/articles/s41586-021-03452-zShan,

Q. Ge, M., Christie, M.J. & Balleine, B.W. (2014) The acquisition of goal-directed actions generates opposing plasticity in direct and indirect pathways in dorsomedial striatum. J Neurosci, 34(28), 9196–9201. doi:10.1523/JNEUROSCI.0313-14.2014.

Sharifi, N., Diehl, N., Yaswen, L., Brennan, M.B. & Hochgeschwender, U. (2001) Generation of dynorphin knockout mice. Brain Res Mol Brain Res, 86(1-2), 70–75. doi:10.1016/s0169-328x(00)00264-3.

Shippenberg, T.S., Zapata, A. & Chefer, V.I. (2007) Dynorphin and the pathophysiology of drug addiction. Pharmacol Ther, 116(2), 306–321. doi:10.1016/j.pharmthera.2007.06.011.

Svingos, A.L., Colago, E.E. & Pickel, V.M. (1999) Cellular sites for dynorphin activation of kappa-opioid receptors in the rat nucleus accumbens shell. J Neurosci, 19(5), 1804–1813. doi:10.1523/JNEUROSCI.19-05-01804.1999.

Tai, L.H., Lee, A.M., Benavidez, N., Bonci, A. & Wilbrecht, L. (2012) Transient stimulation of distinct subpopulations of striatal neurons mimics changes in action value. Nat Neurosci, 15(9), 1281–1289. doi:10.1038/nn.3188, publisher: Nature Publishing Group. URL https://www.nature.com/articles/nn.3188

Tejeda, H.A. & Bonci, A. (2019) Dynorphin/kappa-opioid receptor control of dopamine dynamics: Implications for negative affective states and psychiatric disorders. Brain Research, 1713, 91–101. doi:10.1016/j.brainres.2018.09.023. URL https://www.sciencedirect.com/science/article/pii/S0006899318304888

Tejeda, H.A., Wu, J., Kornspun, A.R., Pignatelli, M., Kashtelyan, V., Krashes, M.J. et al. (2017) Pathway- and Cell-Specific Kappa-Opioid Receptor Modulation of Excitation-Inhibition Balance Differentially Gates D1 and D2 Accumbens Neuron Activity. Neuron, 93(1), 147–163. doi:10.1016/j.neuron.2016.12.005.

Tian, L., Dong, C., Gowrishankar, R., Jin, Y., He, X., Gupta, A. et al. (2023) Unlocking opioid neuropeptide dynamics with genetically-encoded biosensors. ISSN: 2693-5015. URL https://www.researchsquare.com/article/rs-2871083/v1

van den Pol, A.N. (2012) Neuropeptide transmission in brain circuits. Neuron, 76(1), 98–115. doi:10.1016/j.neuron.2012.09.014. URL https://www.ncbi.nlm.nih.gov/pmc/articles/PMC3918222/

Vicente, A.M., Galvão-Ferreira, P., Tecuapetla, F. & Costa, R.M. (2016) Direct and indirect dorsolateral striatum pathways reinforce different action strategies. Current Biology, 26(7), R267–R269. doi:10.1016/j.cub.2016.02.036. URL https://www.sciencedirect.com/science/article/pii/S0960982216300896

Walker, B.M., Zorrilla, E.P. & Koob, G.F. (2011) Systemic κ-opioid receptor antagonism by nor-binaltorphimine reduces dependence-induced excessive alcohol self-administration in rats. Addict Biol, 16(1), 116–119. doi:10.1111/j.1369-1600.2010.00226.x.

Wall, N.R., De La Parra, M., Callaway, E.M. & Kreitzer, A.C. (2013) Differential Innervation of Direct- and Indirect-Pathway Striatal Projection Neurons. Neuron, 79(2), 347–360. doi:10.1016/j.neuron.2013.05.014. URL https://www.sciencedirect.com/science/article/pii/S0896627313004340

Wang, H., Flores, R.J., Yarur, H.E., Limoges, A., Bravo-Rivera, H., Casello, S.M. et al. (2024) Prefrontal cortical dynorphin peptidergic transmission constrains threat-driven behavioral and network states. Pages: 2024.01.08.574700 Section: New Results. URL https://www.biorxiv.org/content/10.1101/2024.01.08.574700v1

Wassum, K.M. & Izquierdo, A. (2015) The basolateral amygdala in reward learning and addiction. Neuroscience & Biobehavioral Reviews, 57, 271–283. doi:10.1016/j.neubiorev.2015.08.017. URL https://www.sciencedirect.com/science/article/pii/S0149763415300373

Wee, S., Orio, L., Ghirmai, S., Cashman, J.R. & Koob, G.F. (2009) Inhibition of kappa opioid receptors attenuated increased cocaine intake in rats with extended access to cocaine. Psychopharmacology (Berl), 205(4), 565–575. doi:10.1007/s00213-009-1563-y.

Yang, R., Tuan, R.R.L., Hwang, F.J., Bloodgood, D.W., Kong, D. & Ding, J.B. (2023) Dichotomous regulation of striatal plasticity by dynorphin. Mol Psychiatry, 28(1), 434–447. doi:10.1038/s41380-022-01885-0.

Yang, W., Carrillo-Reid, L., Bando, Y., Peterka, D.S. & Yuste, R. (2018) Simultaneous two-photon imaging and two-photon optogenetics of cortical circuits in three dimensions. eLife, 7, e32671. doi:10.7554/eLife.32671, publisher: eLife Sciences Publications, Ltd. URL https://doi.org/10.7554/eLife.32671

Yin, H.H., Ostlund, S.B., Knowlton, B.J. & Balleine, B.W. (2005) The role of the dorsomedial striatum in instrumental conditioning. Eur J Neurosci, 22(2), 513–523. doi:10.1111/j.1460-9568.2005.04218.x.

Yoshida, K., Drew, M.R., Kono, A., Mimura, M., Takata, N. & Tanaka, K.F. (2021) Chronic social defeat stress impairs goal-directed behavior through dysregulation of ventral hippocampal activity in male mice. Neuropsychopharmacol., 46(9), 1606–1616. doi:10.1038/s41386-021-00990-y, publisher: Nature Publishing Group. URL https://www.nature.com/articles/s41386-021-00990-y

Zhang, S.X., Kim, A., Madara, J.C., Zhu, P.K., Christenson, L.F., Lutas, A. et al. (2023) Competition between stochastic neuropeptide signals calibrates the rate of satiation. Pages: 2023.07.11.548551 Section: New Results. URL https://www.biorxiv.org/content/10.1101/2023.07.11.548551v2

Zhou, X., Stine, C., Prada, P.O., Fusca, D., Assoumou, K., Dernic, J. et al. (2023) Development of a genetically-encoded sensor for probing endogenous nociceptin opioid peptide release. Pages: 2023.05.26.542102 Section: New Results. URL https://www.biorxiv.org/content/10.1101/2023.05.26.542102v2

